# Molecular Determinants of Ion Selectivity in Heavy-Metal P-type ATPases

**DOI:** 10.64898/2026.09.22.753457

**Authors:** Brisa Raíssa Bartellt Godoy, Hugo Verli

**Affiliations:** Center for Biotechnology, Federal University of Rio Grande do Sul (UFGRS), Av. Bento Gonçalves, 9500, 91501-970, Porto Alegre, Rio Grande do Sul, Brazil; Brazilian Biosciences National Laboratory - LNBio, Brazilian Center for Research in Energy and Materials (CNPEM), Giuseppe Maximo Scolfaro, 10,000, 13083-100, Campinas, São Paulo, Brazil

**Keywords:** P-type ATPase, copper transport, molecular dynamics, metadynamics, ion selectivity, membrane proteins

## Abstract

P-type ATPases maintain cellular metal homeostasis by coupling ATP hydrolysis to the active translocation of ions across biological membranes. Within this superfamily, P_1_*B* -type ATPases play a central role in the transport and detoxification of transition metals such as Cu^+^ and Ag+. Despite the availability of several crystallographic structures for bacterial ATPases, molecular determinants governing ion selectivity, coordination dynamics, and transport efficiency are not yet fully resolved, particularly for plant homologs, whose conformational landscapes have not been explored in detail at atomistic resolution. In this study, we combine molecular dynamics (MD) simulations with enhanced sampling through metadynamics to characterize the conformational dynamics and ion-binding mechanisms of representative heavy-metal P-type ATPases. By comparing bacterial and plant homologs in phosphorylated and dephosphorylated states, we examine how phosphorylation is associated with changes in structural plasticity, channel accessibility and ion coordination environments. The simulations indicate phosphorylation-dependent rearrangements of the cytosolic domains and transmembrane helices, which may alter access to the metal-binding pathway. Enhanced sampling identifies distinct conformational basins and possible transition pathways along functionally relevant collective variables, allowing characterization of the free-energy landscape associated with ion displacement within the transmembrane region. Together, these results provide an atomistic perspective on the relationship between metal coordination and protein conformational dynamics. Overall, our findings support a model in which phosphorylation state, structural dynamics, and local coordination environments collectively contribute to the differential behavior of metal ions in P_1_*B* -type ATPases.

## 1 Introduction

P-type ATPases are membrane proteins that play a central role in cellular homeostasis by transporting substrates across biological membranes through ATP hydrolysis [1, 2, 3]. Their catalytic cycle follows the canonical Post-Albers mechanism [4, 5], in which the protein alternates between E1 and E2 conformations through the phosphorylation and dephosphorylation of a highly conserved aspartate residue in the cytosolic phosphorylation (P) domain [6, 2]. In the E1 state, the cytosolic headpiece is arranged to facilitate ATP binding and substrate loading, while phosphorylation promotes the transition to E2 conformation associated with substrate release on the opposite side of the membrane [7, 8, 9]. This cycle couples ATP hydrolysis to vectorial ion transport and depends on coordinated rearrangements between the actuator (A), nucleotide-binding (N), phosphorylation (P), and transmembrane (TM) domains [7, 10, 2].

Within this superfamily of ion-pumps, P_1*B*_-type ATPases are specialized in the transport of transition metals, including monovalent ions such as Cu^+^ and Ag^+^ and divalent ions such as Zn^2+^, Cd^2+^, and Co^2+^ [11, 12, 13]. Therefore, their high substrate specificity is essential for balancing metal acquisition, detoxification, and intracellular distribution while preventing toxic accumulation. Despite the availability of high-resolution structures and extensive biochemical characterization for selected P_1*B*_-ATPases [14, 10, 15], the molecular determinants that couple phosphorylation to ion translocation and selectivity remains only partially characterized, especially in plant P_1*B*_-ATPases. In particular, it is still unclear how phosphorylation reshapes long-range interdomain communication, modulates the relative positioning of the catalytic domains, and reorganizes the transmembrane cavity during transport. These questions are especially relevant for plant heavy-metal ATPases, for which atomistic dynamic information remains much more limited than for bacterial prototypes.

The three systems investigated here were selected to enable comparison between a structurally characterized bacterial transporter and two physiologically relevant plant homologues. *Legionella pneumophila* CopA (LpCopA) was selected as a reference because it is among the best-characterized P1*B*-ATPases. Crystal structures assigned to the E2P_i_ and E2P states have provided an important structural framework for investigating heavy-metal transport by this family [14, 10]. In plants, P1*B*-ATPases comprise multiple heavy-metal transporters involved in metal homeostasis, detoxification, and distribution. The model plant *Arabidopsis thaliana* encodes eight HMA transporters, AtHMA1-AtHMA8, whereas the rice genome, *Oryza sativa*, contains nine homologues, OsHMA1-OsHMA9 [16, 17]. Although computational studies have previously examined selected plant HMAs, including the transmembrane region of AtHMA4, atomistic investigations directly focused on the conformational dynamics of rice P_1*B*_-ATPases remain limited. To our knowledge, no previous molecular dynamics study has specifically examined the phosphorylation-associated dynamics and ion interactions of OsHMA9.

Molecular dynamics simulations provide an atomistic framework for examining the structural flexibility and collective motions of membrane proteins beyond static experimental structures [18]. Enhanced-sampling approaches, including metadynamics and umbrella sampling, can additionally be used to characterize selected free-energy processes associated with conformational rearrangements and ion movement [19]. Together, these approaches enable the comparison of phosphorylation-dependent structural responses and ion-protein interactions across different P_1*B*_-ATPases.

In this work, we use molecular dynamics simulations and complementary structural analyses to compare the conformational behavior of LpCopA, AtHMA5, and OsHMA9 in dephosphorylated and phosphorylated conditions. We focus on phosphorylation-associated changes in protein flexibility, interdomain organization, collective motions, and communication between the cytosolic and transmembrane regions. Free-energy calculations are additionally used to characterize selected ion-protein interactions and ion-translocation events. By comparing a structurally characterized bacterial transporter with two plant homologues, we aim to identify conserved and system-specific dynamical responses that may contribute to the functional behavior of P_1*B*_-ATPases.

## 2 Results and Discussion

### 2.1 Global conformational changes and domain-specific flexibility

In order to characterize the general dynamical behavior of all six systems studied (LpCopA, LpCopA-P, OsHMA9, OsHMA9-P, AtHMA5, and AtHMA5-P), we initially analyzed the RMSD and RMSF profiles (Fig. 1). P_1*B*_-ATPases are known to be structurally dynamic enzymes, containing a high degree of mobility, predominantly in the cytosolic domains, which are connected to a more rigid transmembrane domain. Consequently, global backbone RMSD of proteins with this multidomain architecture should not be interpreted as an indicator of structural stability or unfolding, as it mainly reports the amplitude of large-scale domain rearrangements sampled during the MD simulations.

**Figure 1.**
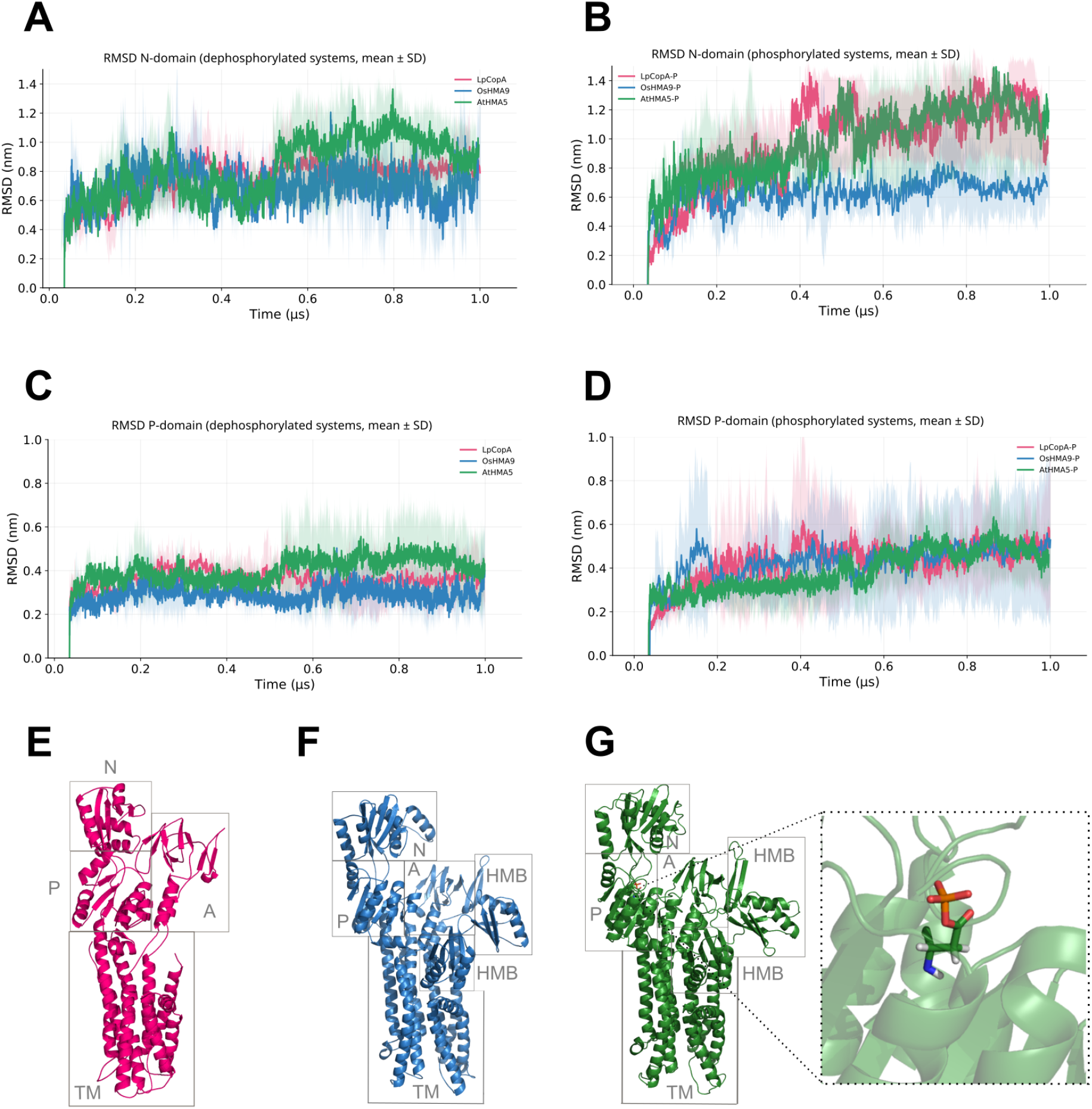
Global and domain-resolved structural dynamics of P_1*B*_-ATPases across species and phosphorylation states. **(A)** Backbone RMSD as a function of time for LpCopA, OsHMA9 and AtHMA5, both in dephosphorylated (LpCopA, OsHMA9 and AtHMA5) and phosphorylated (LpCopA-P, OsHMA9-P and AtHMA5-P) states. Curves represent averages over three independent trajectories per system; shaded regions indicate the corresponding standard deviations. **(B)** Backbone RMSF profiles showing residue-level flexibility. Rigid transmembrane segments are conserved across all systems, whereas enhanced fluctuations are localized in cytosolic loops, plant-specific insertions, and N-terminal heavy-metal–binding regions. **(C)** Domain-specific RMSD for the A, P, N, and TM domains. The TM domain remains the most rigid in all systems. Phosphorylation selectively increases fluctuations in the cytosolic headpiece, with the strongest effects observed in the N-domain of LpCopA-P and AtHMA5-P, and in the P-domain of OsHMA9-P. Plant homologs display systematically higher domain RMSD than bacterial CopA. **(D)** Representative structures of LpCopA and OsHMA9 highlighting the nucleotide-binding (N), actuator (A), phosphorylation (P), transmembrane (TM), and heavy-metal–binding (HMB) domains. Colored markers correspond to the domain definitions used in panel C.

All systems reach an apparent plateau after the initial equilibration period and remain within relatively stable RMSD ranges throughout the microsecond-long simulations, suggesting that the systems remained within a limited conformational range during the simulations. Although all systems followed a similar overall pattern, their RMSD profiles differed across homologs and phosphorylation states. The phosphorylated LpCopA and AtHMA5 systems show the largest backbone deviations, while the dephosphorylated OsHMA9 presents a smaller conformational range, suggesting lower global conformational variability in the absence of phosphorylation. A similar pattern has been reported in previous studies on different P-type ATPases, where phosphorylation has been linked to increased conformational flexibility and large-scale domain rearrangements, which are required for ion translocation [20, 21, 22]. Dephosphorylated states, however, tend to show lower structural variability and can remain in more compact conformations associated with resting or pre-activated stages of the catalytic cycle [18, 23].

To distinguish between structural loss and interdomain movements as possible sources of the global deviations observed in RMSD profiles, we analyzed the time evolution of the secondary structure of each MD replica using the Define Secondary Structure of Proteins (DSSP) algorithm [24] (Supplementary Figs. S6-8). DSSP assigns secondary structure elements based on hydrogen-bonding patterns and backbone geometry, allowing a residue-level assessment of the structural stability of the simulations [25]. Across all replicas and all six systems, the main *α*-helices and *β*-sheets remained mostly preserved throughout the trajectories, with the observed changes being small and temporary, occurring mainly in regions of loops and turns. These DSSP profiles showed no consistent loss of helical or sheet structure, indicating that the overall protein fold was maintained during the simulations, which suggests that the RMSD variations are mainly associated with domain movements rather than with structural unfolding.

A similar structural behavior has been reported for LpCopA [10], where relatively high RMSD values (>4 Å) were observed for the E2.Pi state, while the E2P state displayed lower deviations (2 Å). Importantly, these elevated RMSD values were not associated with structural instability of the transmembrane region, which remained stable and reached a plateau at approximately 2 Å after 20 ns of MD simulations. Instead, the increase in RMSD was attributed to the intrinsic flexibility of the large cytosolic headpiece and the loop regions between the cytosolic domains. Similar results have also been reported for other P-type ATPases, especially SERCA. MD studies of these proteins showed that the cytosolic A, N, and P domains are highly flexible and often reach RMSD values above 3-5 Å, while the transmembrane domain remains more stable [26, 27]. Therefore, high global RMSD values in these systems are more likely to reflect movements between domains and the natural flexibility of the cytosolic regions than protein unfolding.

The RMSF profiles also suggest that flexibility is not equally distributed across the protein structure. In all systems, the transmembrane helices show relatively low fluctuations (Fig. S5), whereas higher values are mainly observed in the cytosolic headpiece, including the loops connecting the A, P, and N domains (Fig. S4) and additional segments related to regulatory or metal-binding functions. In the plant homologs, the fluctuations appear to be spread over a larger portion of the cytosolic region, which may be related to their larger architecture and to the presence of additional heavy-metal-binding domains (HMBDs) that are absent from the bacterial transporter.

Overall, the RMSF results do not suggest that phosphorylation causes a general loss of stability, but they show local changes in flexibility that differ among the homologs and between the cytosolic and transmembrane regions. In LpCopA, the catalytic domains remain relatively compact, while the plant homologs show larger fluctuations, mainly in the cytosolic headpiece and, in some cases, in parts of the transmembrane region. These differences suggest that phosphorylation may change domain flexibility in different ways across the three systems, while the overall protein structure remains preserved.

#### 2.1.1 N-P domain distance varies with phosphorylation

To investigate how phosphorylation affects the catalytic region in terms of architecture, we monitored the distance between the centers of mass of selected residues in the N- and P-domains during the MD simulations. The selected N-domain residues are associated with the nucleotide-binding region, whereas the P-domain selection includes the conserved aspartate residue that undergoes phosphorylation. This distance provides a simple measure of the relative position of functionally important regions in the N- and P-domains, which are involved in nucleotide binding, phosphate transfer, and conformational changes in P-type ATPases. For each system, the mean distance was first calculated for each of the three independent replicas and the final mean and standard deviation were then obtained from these three replica averages (Table S2). In the dephosphorylated systems (Fig. 2B), LpCopA showed an average distance of 2.36 *±* 0.03 nm. OsHMA9 and AtHMA5 showed shorter average distances of 1.97 *±* 0.22 nm and 1.98 *±* 0.19 nm respectively. These results suggest that the selected N- and P-domain regions were more separated in LpCopA than in the two plant homologs. The low standard deviation observed for LpCopA also indicates that the mean distance was similar across the three replicas, while the larger standard deviations of OsHMA9 and AtHMA5 indicated greater variation among replicas in the dephosphorylated plant systems.

**Figure 2.**
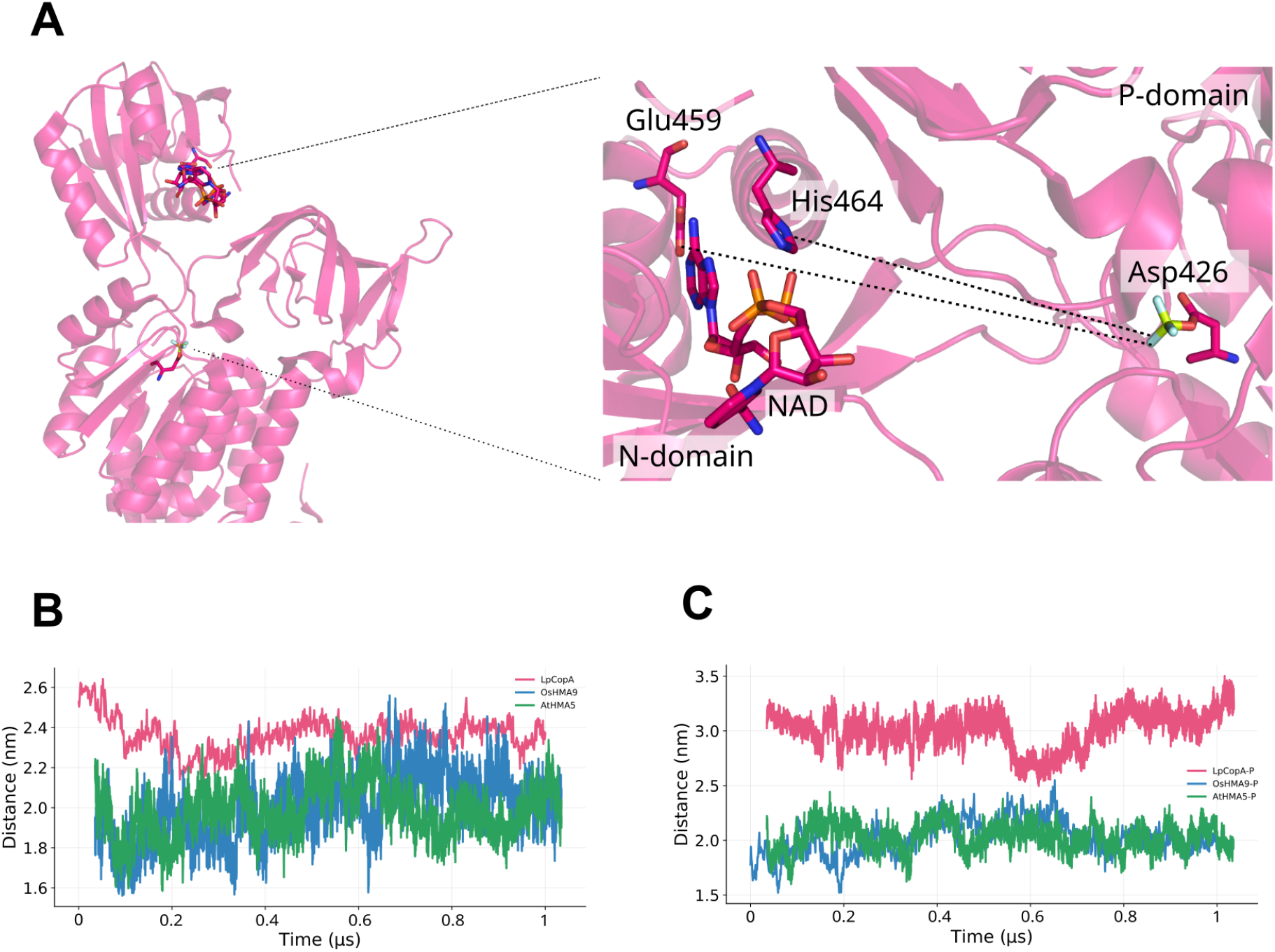
Phosphorylation-dependent modulation of the relative arrangement of selected N- and P-domain regions in P_1*B*_-ATPases. **(A)** Structural representation of the monitored distance between the centers of mass of selected residues in the nucleotide-binding region of the N-domain and the region encompassing the conserved phosphorylatable aspartate in the P-domain. The inset highlights the selected residues and their positions relative to the nucleotide-binding interface. **(B)** Time evolution of the monitored distance in the dephosphorylated systems. Mean distances were 2.36 ± 0.03 nm for LpCopA, 1.97 ± 0.22 nm for OsHMA9, and 1.98 *± ± ±* 0.19 nm for AtHMA5. **(C)** Time evolution of the same distance in the phosphorylated systems. Mean distances were 3.01 ± 0.35 nm for LpCopA-P, 2.02 ± 0.06 nm for OsHMA9-P, and 2.03 *±* 0.26 nm for AtHMA5-P. Values are reported as the mean ± sample standard deviation calculated from the mean distance of three independent replicas. Phosphorylation was associated with a pronounced separation of the monitored N- and P-domain regions in LpCopA, whereas the plant homologs retained comparatively shorter distances.

The differences became clearer when comparing the phosphorylated systems (Fig. 2C). LpCopA-P showed an average distance of 3.01 *±* 0.35 nm, representing an increase of about 0.65 nm compared with the dephosphorylated state. In contrast, OsHMA9-P and AtHMA5-P showed average distances of 2.02 *±* 0.06 nm and 2.03 *±* 0.26 nm, respectively, which were close to the values measured in the corresponding dephosphorylated systems. The low standard deviation of OsHMA9-P indicates that the mean distance was similar across replicas, whereas LpCopA-P and AtHMA5-P showed greater variation among the simulations.

These results suggest that phosphorylation can be linked to a larger separation between the selected N- and P-domain regions in LpCopA, while no similar increase was seen in the plant homologs. The greater distance observed in LpCopA-P may be related to the large movements of the cytosolic headpiece reported for phosphorylated states of several P-type ATPases, including more open arrangements of the catalytic domains in E2P conformation [28, 29, 30, 31]. In OsHMA9-P and AtHMA5-P, the selected regions showed distances similar to those of the dephosphorylated states, suggesting that phosphorylation may involve different domain arrangements in bacterial and plant P_1*B*_-ATPases. These differences may result from distinct domain arrangements and flexibility in the three proteins. Previous studies have also shown that, although the catalytic core is conserved, conformational changes can differ among P-type ATPase families and homologs [14, 10, 32, 33, 34].

### 2.2 Essential dynamics shows conserved collective motions and different responses to phosphorylation

Principal component analysis (PCA) was used in order to investigate the main collective motions sampled by the six P_1*B*_-ATPase systems (Fig. 3). Unlike local fluctuation analyses, the first principal components (PCs) describe coordinated movements involving the cytosolic domains and the transmembrane helices, capturing the main conformational changes sampled during the simulations. Among the three homologs, LpCopA showed the clearest separation between the phosphorylated and dephosphorylated states in the PCA projection, with the two conformational ensembles populating partly different regions and suggesting that phosphorylation can modify the range of conformations sampled rather than only increasing the fluctuations within the same region.

**Figure 3.**
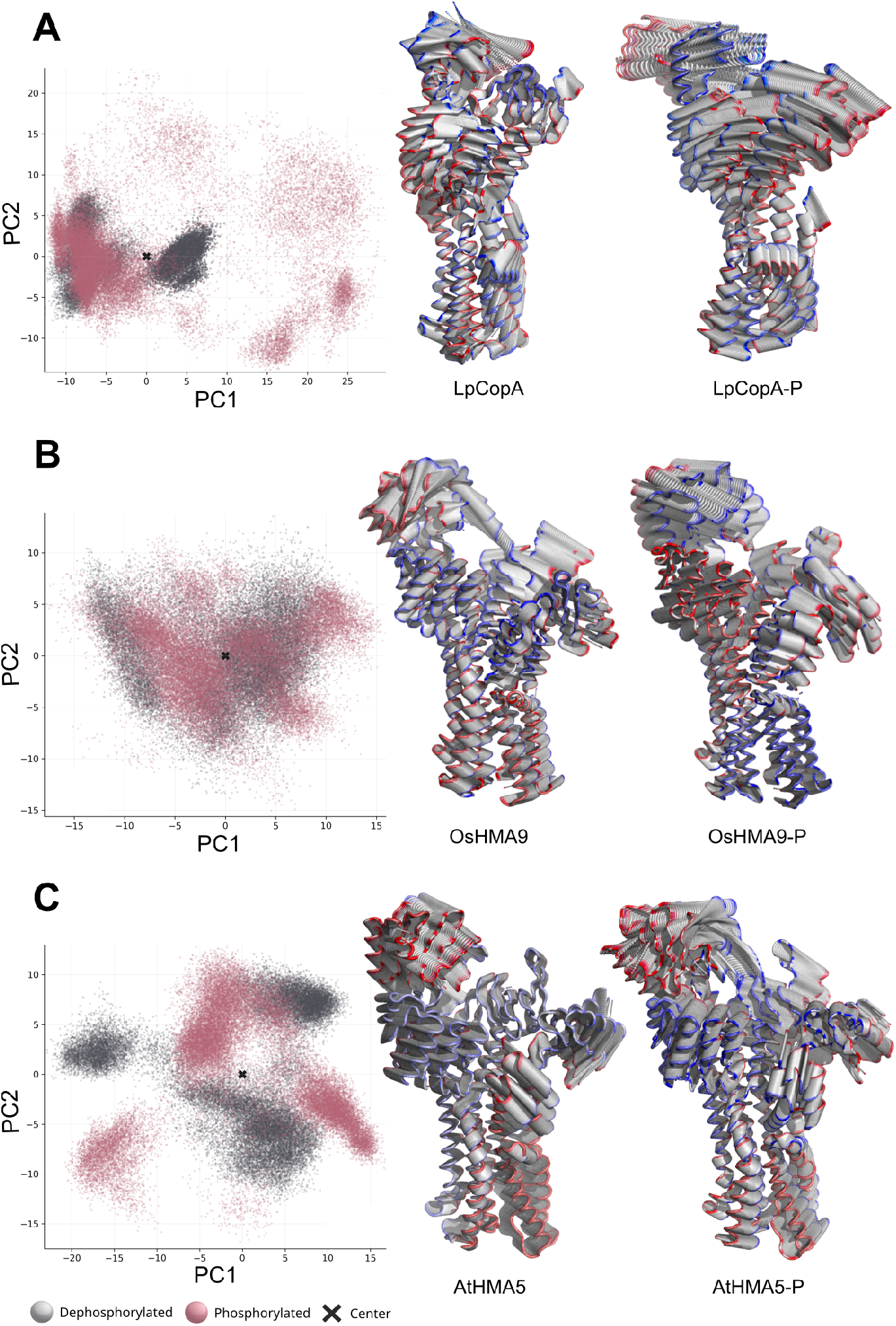
Principal component analysis (PCA) of conformational sampling in bacterial and plant P1*B*-ATPases. (A) *Legionella pneumophila* CopA (LpCopA), (B) *Oryza sativa* HMA9 (OsHMA9), and (C) *Arabidopsis thaliana* HMA5 (AtHMA5). For each homolog, PCA was performed independently using the combined ensemble of dephosphorylated and phosphorylated trajectories obtained from multiple MD replicas initiated with different initial velocities. Left panels show the projection of dephosphorylated (gray) and phosphorylated (pink) conformations onto the first two principal components (PC1 and PC2) defined for each homolog-specific conformational space; the black “X” marks the ensemble center. Right panels show representative interpolated conformations along PC1 for the dephosphorylated and phosphorylated states, illustrating the main collective rearrangements sampled within each homolog-specific PCA space. The relative overlap or separation between phosphorylation states indicates that phosphorylation reshapes the sampled conformational landscape in a homolog-dependent manner.

The plant homologs presented different patterns. In OsHMA9, the phosphorylated and dephosphorylated ensembles overlap in the PC1-PC2 projection, although the phosphorylated state showed a broader distribution within the same region of conformational space. The overlap between the two OsHMA9 states suggests that phosphorylation did not produce a major shift in the dominant conformational space, while the phosphorylated ensemble showed greater dispersion within the same region. AtHMA5 showed a more heterogeneous distribution in its PCs projection, with partially separated regions and, when compared to LpCopA and OsHMA9, this visual pattern suggests that AtHMA5 samples a less compact conformational ensemble within the simulated timescale. However, because the PCA spaces were generated independently for each homolog, this comparison should be interpreted qualitatively and within the context of each homolog-specific conformational landscape.

The structures along PC1 (right panels in Fig. 3) help illustrate the main motions represented by this component. In all three homologs, the motion involves displacement of the cytosolic domains relative to the transmembrane region, which is compatible with the large-scale domain movements described for P-type ATPases [35, 36]. However, the extent and direction of these motions differ among the systems. In LpCopA, the interpolation suggests a clearer phosphorylation-dependent repositioning of the catalytic headpiece. In OsHMA9, the motion appears more distributed, consistent with the overlap observed in PC space. In AtHMA5, the interpolated frames show larger movements of the cytosolic domains, suggesting a broader range of conformational changes during the simulations.

The main principal components described a large-scale motion involving the cytosolic catalytic domains and the transmembrane region, but the way this conformational space is sampled varies among the homologs and between phosphorylation states. In LpCopA and AtHMA5, the phosphorylated and dephosphorylated systems occupy partly different regions of the PC1-PC2 projection, which may reflect the sampling of distinct conformational states. In OsHMA9, the two ensembles overlap more strongly, suggesting that phosphorylation mainly changes the distribution of structures within a similar region of conformational space. Overall, the effect of phosphorylation was not the same in the three P_1*B*_-ATPases and appeared to depend on the conformational behavior of each homolog.

### 2.3 Dynamic networks of interdomain communication

Dynamic network analysis was performed to study correlated residue motions in the dephosphorylated and phosphorylated P_1*B*_-ATPase systems (Fig. 4 and Supplementary Figs. S9-11). In each transporter, several dynamic communities were identified across the cytosolic and transmembrane regions.

**Figure 4.**
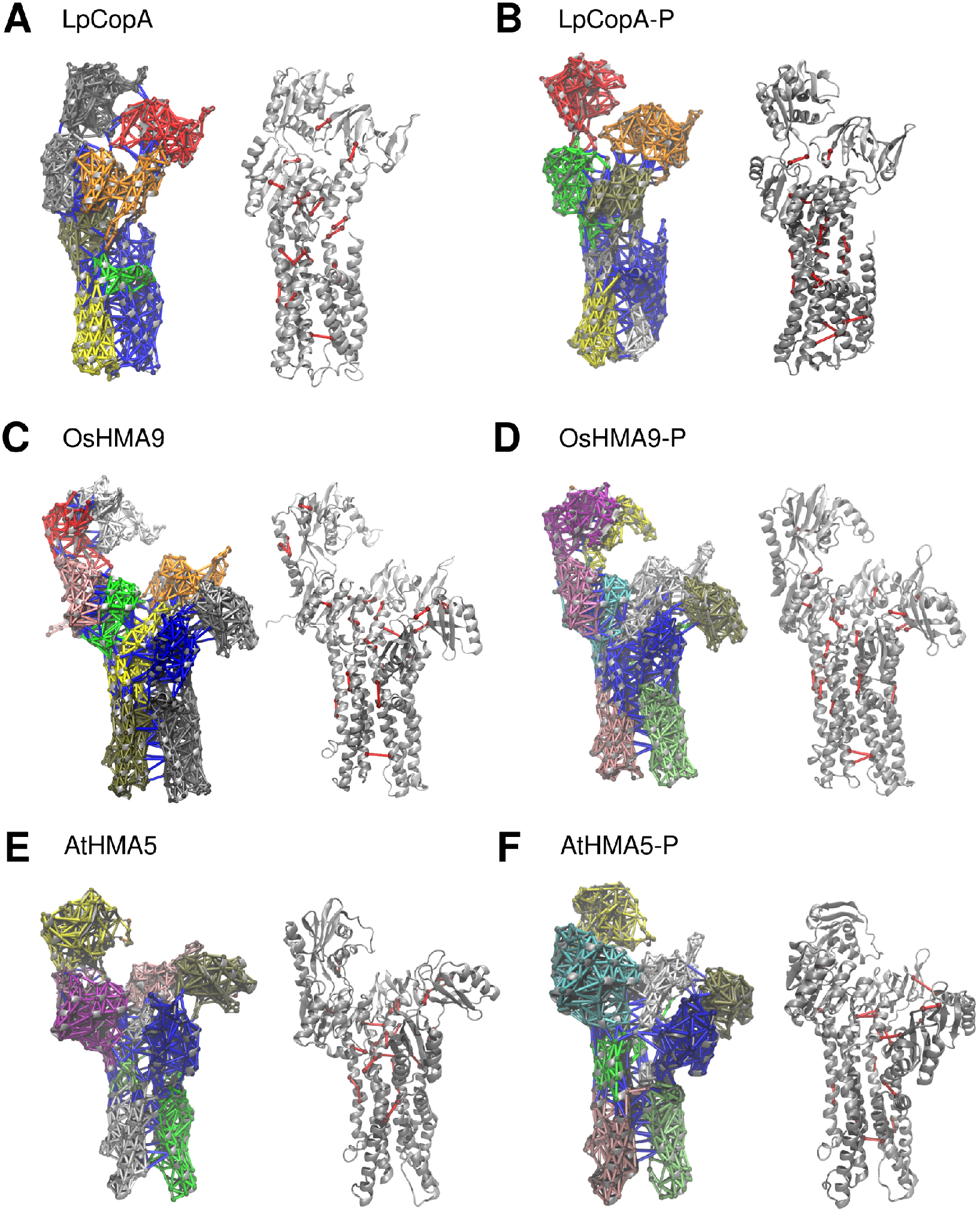
Dynamic Network Analysis (DNA) and critical nodes for representative replicas. Dynamic network communities and critical nodes for representative single replicas of each system. The complete set of three independent replicas per system is provided in Supplementary Figs. S9-11. For each system, the left subpanel shows the community decomposition of the residue-residue dynamic network, where colors indicate groups of residues undergoing correlated motions. The right subpanel displays the corresponding critical nodes mapped onto the protein structure, where red segments identify residues with higher betweenness centrality.

By comparing the two states analyzed for each system, we observed changes in community distribution (Fig. 4 A-F, left) and in the positions of critical nodes (Fig. 4 A-F, right). In the left-hand structures, the different dynamic communities are represented by distinct colors. In the right-hand structures, the proteins are shown in gray, with the critical nodes represented as red spheres, which correspond to residues that are frequently found along the shortest paths that connect different regions of the network. In LpCopA, changes were observed in both community boundaries and critical-node positions, mainly in the cytosolic headpiece and near its interface with the transmembrane region. OsHMA9 and AtHMA5 also showed changes in community organization and in the distribution of critical nodes after phosphorylation, but the number of communities did not change in the same way across the three homologs.

The clearest phosphorylation-dependent changes were observed in LpCopA, where community distribution and the position of each critical node shifted mainly in the cytosolic headpiece and near the interface between the cytosolic and transmembrane regions. Changes were also observed in OsHMA9 and AtHMA5, although no single pattern was shared by the three homologs. The number of communities, their organization and the location of critical nodes varied differently in each system. Thus, phosphorylation appears to reorganize correlated residue motions in a homolog-dependent manner rather than producing a common network response across P_1*B*_-ATPases. The complete replica analysis and quantitative network descriptors are provided in the Supplementary Information.

### 2.4 Ion-dependent free-energy landscapes

Metadynamics simulations were used to examine the free-energy landscapes associated with ion transport in bacterial and plant P_1*B*_-ATPases. For each ion, system, and phosphorylation state, the FES shown in Figs. 5-7 represents the average of three independent replicas projected onto the same CV space. Before averaging, the minimum of each individual FES was shifted to zero, and the averaged surfaces were used to describe the main energetic patterns shared across replicas, while the differences of each replica were also analyzed separately.

**Figure 5.**
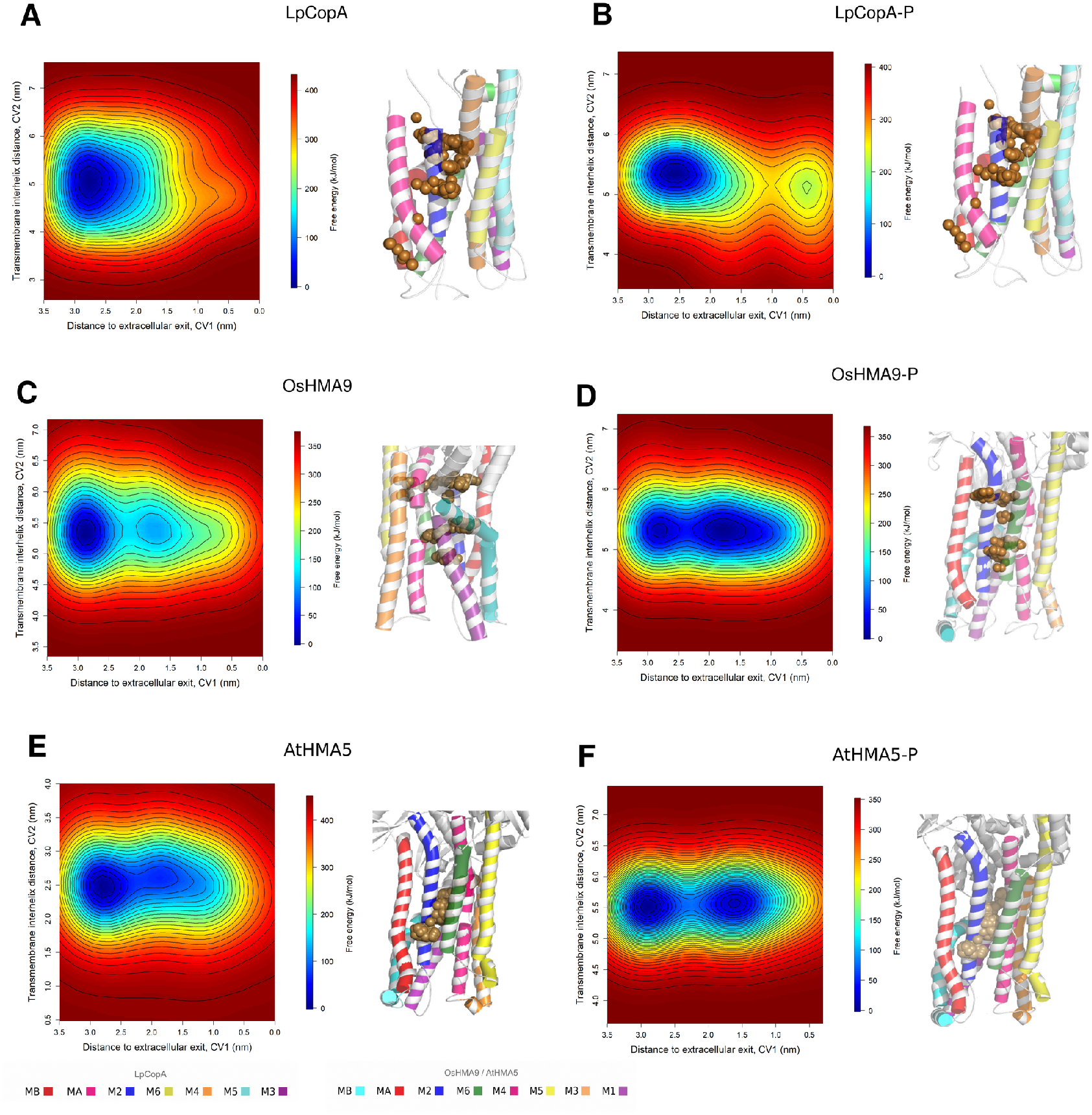
Averaged free-energy landscapes (FES) and representative ion positions for Cu^+^ transport across P1*B*-ATPases. Each two-dimensional FES represents the pointwise average of three independent metadynamics FES reconstructed for the same system and ion, after shifting the free-energy minimum of each replicate to zero. The FES were projected along the distance to the extracellular exit (CV1) and the transmembrane interhelical distance (CV2). (A) LpCopA in the dephosphorylated state. (B) LpCopA in the phosphorylated state. (C) OsHMA9 in the dephosphorylated state. (D) OsHMA9 in the phosphorylated state. (E) AtHMA5 in the dephosphorylated state. (F) AtHMA5 in the phosphorylated state. Adjacent structural panels show representative Cu^+^ positions mapped onto the transmembrane region, highlighting the main ion-sampling regions along the pore. Overall, the averaged landscapes indicate transportcompatible low-energy regions for Cu^+^, with phosphorylation reshaping basin organization along CV1 in a system-dependent manner.

The time evolution of the free-energy profiles was evaluated qualitatively by reconstructing one-dimensional FES at different stages of each metadynamics simulation. The profiles were compared based on changes in the position and shape of the main minima over time, which allowed us to follow the evolution of the main free-energy features and identify replicas with greater temporal variation. The complete replica-specific analyses are presented in the Supplementary Material. The individual replica-specific FES and the corresponding convergence analyses are presented in Supplementary Figs. S15-S50.

For Cu^+^, the averaged FES show low-energy regions within the sampled CV space for all three homologs (Fig. 5). In LpCopA, the dephosphorylated system presents a broad minimum centered at higher CV1 values, whereas the phosphorylated state shows an additional low-energy region at lower CV1 values. A similar change is observed in OsHMA9, where phosphorylation produces a broader low-energy region with two distinguishable minima along CV1. In AtHMA5, both states show extended low-energy regions, although the phosphorylated system also presents a clearer separation between two minima, indicating that phosphorylation changes the distribution of the regions of lower energy in all three systems, but the extent and shape of these changes differ among the homologs.

The structural representations beside the FES (right panels in Fig. 5) show selected Cu^+^ trajectories from one metadynamics simulation chosen for visualization for each system and ion. In LpCopA and OsHMA9, several positions are located around the central part of the transmembrane domain, while additional positions extend toward other regions of the pore. AtHMA5 also shows Cu^+^positions distributed along the transmembrane region. Together with the FES, these structures illustrate that Cu^+^samples more than one position within the pore and that the energetic distribution of these positions changes with phosphorylation.

At the residue level, the analysis of contact frequency (Fig. S12) between the ion and pore residues shows differences in Cu^+^ interactions among the three homologs and phosphorylation states. In LpCopA, Glu205 shows frequent contacts with Cu^+^ in the dephosphorylated system while Gln202 also contributes in several replicas and, after phosphorylation, the contact pattern changes with Met148 appearing more frequently among the residues interacting with the ion. Met148 and Glu205 have previously been proposed as part of the Cu^+^ entry region of LpCopA, together with Asp337, and Met148 has also been associated with a transient Met148-Cys382 coordination site during Cu^+^entry [37]. Previous work has also shown the importance of methionine for sulfur-containing residues in Cu^+^ binding and translocation in P_1*B*_-ATPases [38]. In OsHMA9, Met142, which occupies a position related to Met148 in LpCopA, also shows contacts with Cu^+^, although their frequency varies among replicas. and when comparing with AtHMA5, we observed a more dispersed contact pattern with no single residue maintaining high contact frequencies across all replicas, pointing to a more variable set of Cu^+^ interactions in AtHMA5 under the simulated conditions.

For Ag^+^, the averaged FES shows a more dispersed distribution of low-energy regions across the sampled CV space (Fig. 6). In LpCopA, the dephosphorylated system presents a main low-energy region, while phosphorylation results in two different minima along CV1. OsHMA9 shows an extended low-energy region in both states, with only small changes in its organization after phosphorylation, indicating that Ag^+^ is not restricted to a single well-defined combination of CV1 and CV2, but can occupy a range of energetically favorable configurations within the sampled space. A similar behavior is observed in AtHMA5, although the phosphorylated system presents a more extended low-energy region.

**Figure 6.**
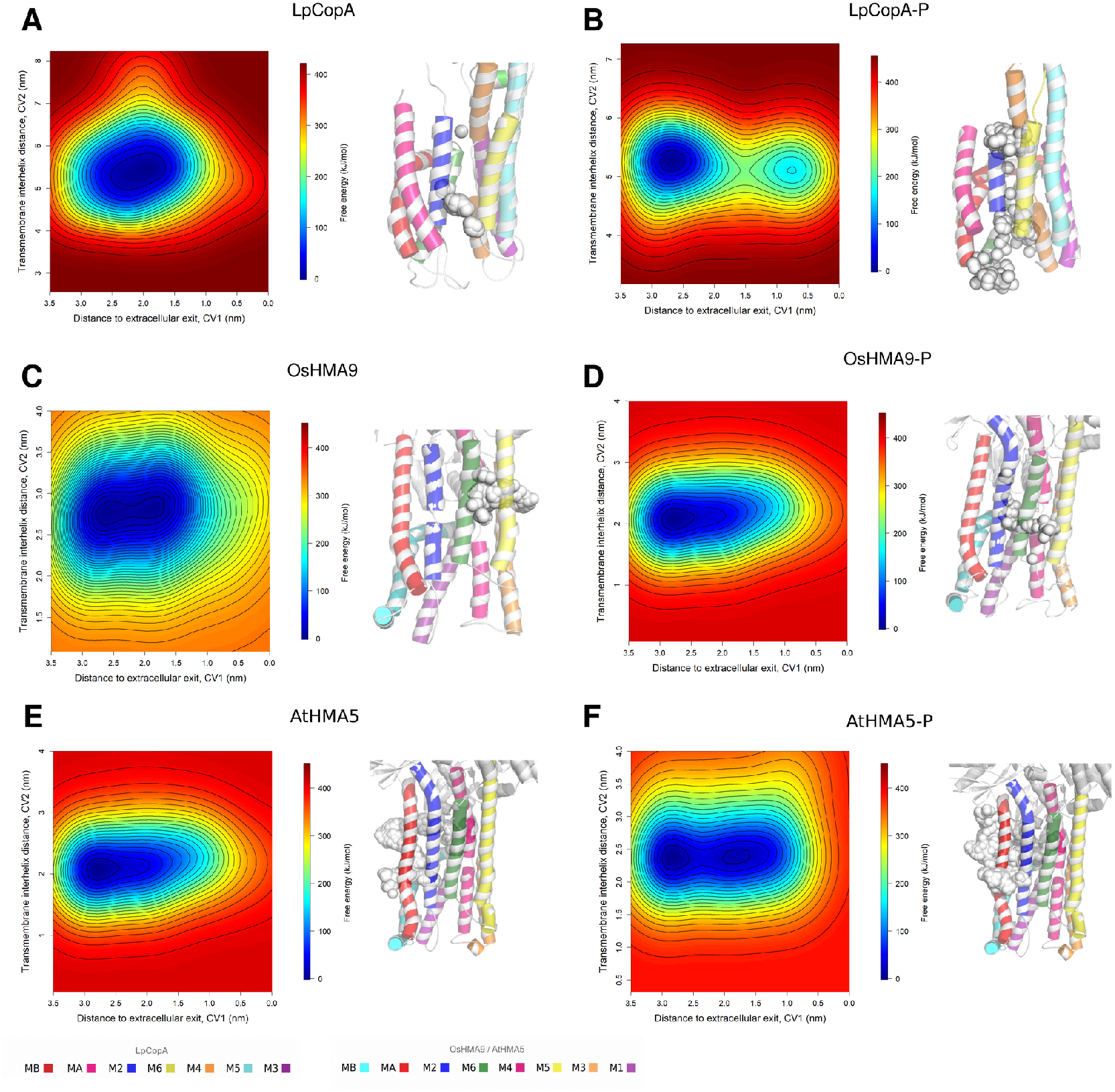
Averaged free-energy landscapes (FES) and representative ion positions for Ag+ transport across P1*B*-ATPases. Each two-dimensional FES represents the pointwise average of three independent metadynamics FES reconstructed for the same system and ion, after shifting the free-energy minimum of each replicate to zero. The FES were projected along the distance to the extracellular exit (CV1) and the transmembrane interhelical distance (CV2). (A) LpCopA in the dephosphorylated state. (B) LpCopA in the phosphorylated state. (C) OsHMA9 in the dephosphorylated state. (D) OsHMA9 in the phosphorylated state. (E) AtHMA5 in the dephosphorylated state. (F) AtHMA5 in the phosphorylated state. Adjacent structural panels show representative Ag+ positions within the transmembrane domain, indicating accessible coordination regions along the pore. Compared with Cu^+^, the averaged FES for Ag+ displays a more heterogeneous energetic landscape, with multiple low-energy regions and systemdependent pathway connectivity. Phosphorylation modulates basin organization and pathway accessibility, although the extent of this effect varies among the bacterial and plant transporters.

Overall, the Ag^+^ landscapes indicate that the ion can sample different regions of the trans-membrane pore, but the effect of phosphorylation varies among the three homologs. This behavior agrees with the chemical similarity between Cu^+^ and Ag^+^, since both are monovalent ions with affinity for soft ligands. However, Ag^+^ is not considered a physiological substrate of these Cu^+^-transporting transporters, as silver has no established biological role and is generally associated with metal toxicity rather than normal cellular metal homeostasis in P-type ATPases [39, 40].

The structural panels shown on the right in Fig. 6 complement the FES by showing Ag^+^ positions sampled across the transmembrane region. When compared with Cu^+^, the interactions between Ag^+^ and each transporter pore residues are more distributed among different residues and vary considerably between replicas. In LpCopA, residues such as Glu189, Ile194, Ala192, and neighboring positions contribute to Ag^+^ interactions in the dephosphorylated state, whereas phosphorylation changes the contact pattern and increases the contribution of residues such as Arg268, Glu189, Asp272, and Met148 (Fig. S13). A similar replica-dependent behavior is observed in OsHMA9 and AtHMA5. This more heterogeneous interaction pattern suggests that Ag^+^ can be accommodated by several local environments within the transmembrane region, but with a less defined coordination pattern than that observed for Cu^+^. This is consistent with the chemical similarity between Cu^+^ and Ag^+^, while also indicating that the pore does not interact with the two monovalent ions in exactly the same way.

Cd^2+^ displays a distinct energetic profile from the monovalent ions (Fig. 7). In the three homologs, the averaged FES is dominated by relatively localized low-energy regions, corresponding to a narrower range of favorable combinations of CV1 and CV2. This suggests that Cd^2+^ is more restricted to specific configurations within the transmembrane region, rather than sampling a more extended set of favorable states. In LpCopA, both phosphorylation states retain a compact main minimum, indicating that phosphorylation modifies the local energetic environment but does not substantially increase the range of favorable Cd^2+^ configurations. OsHMA9 shows a similar restriction, although phosphorylation shifts the position and shape of the minimum. In AtHMA5, the dephosphorylated state samples a somewhat more extended region, whereas the phosphorylated system becomes more localized. These differences indicate that phosphorylation affects Cd^2+^ energetics in a homolog-dependent manner, but does not produce the broader redistribution observed for the monovalent ions.

**Figure 7.**
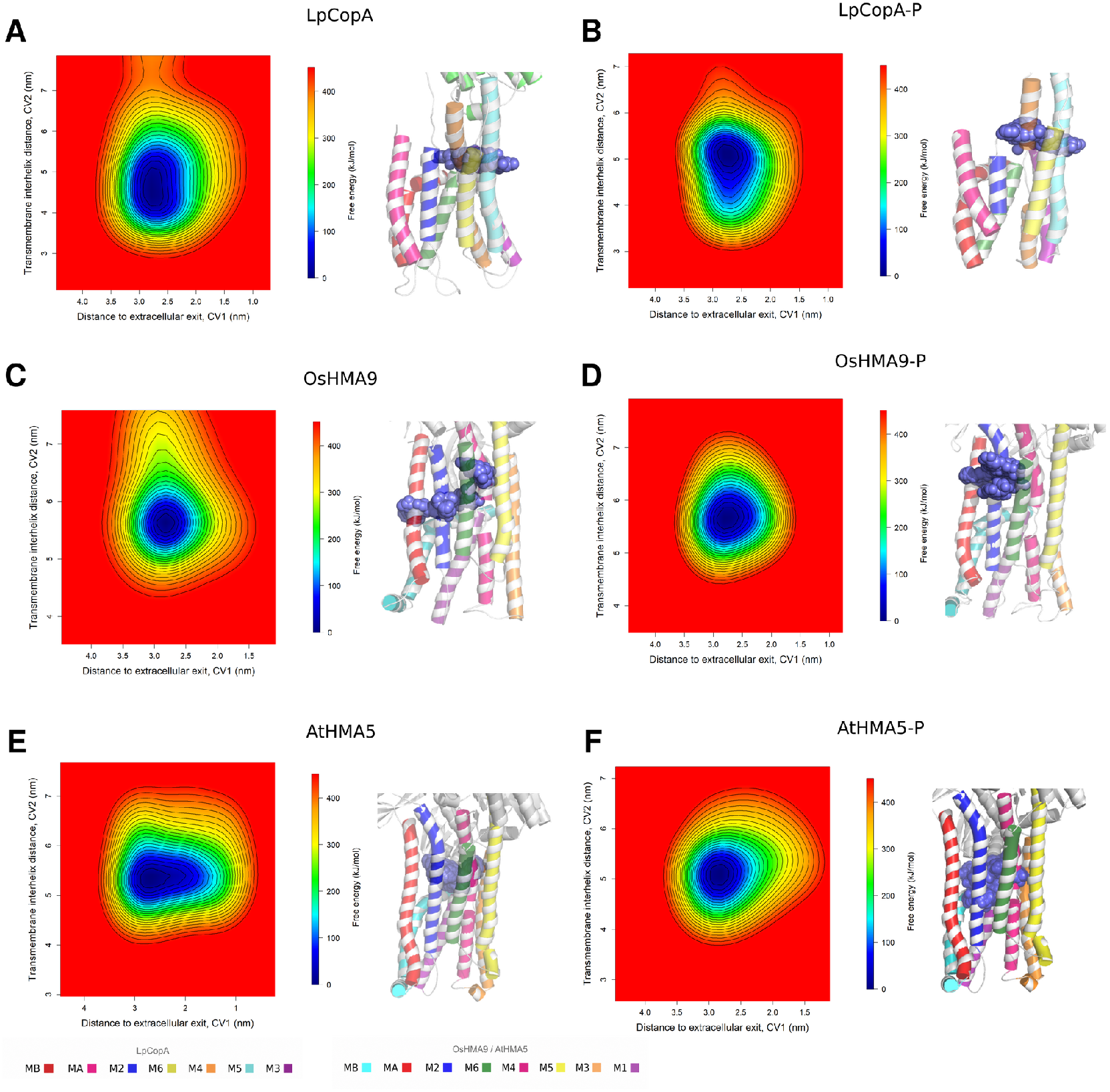
Averaged free-energy landscapes (FES) and representative ion positions for Cd^2+^ transport across P_1*B*_-ATPases. Each two-dimensional FES represents the pointwise average of three independent metadynamics FES reconstructed for the same system and ion, after shifting the free-energy minimum of each replicate to zero. The FES were projected along the distance to the extracellular exit (CV1) and the transmembrane interhelical distance (CV2). (A) Lp-CopA in the dephosphorylated state. (B) LpCopA in the phosphorylated state. (C) OsHMA9 in the dephosphorylated state. (D) OsHMA9 in the phosphorylated state. (E) AtHMA5 in the dephosphorylated state. (F) AtHMA5 in the phosphorylated state. Adjacent structural panels show representative Cd^2+^ positions mapped onto the transmembrane region, highlighting restricted sampling along the pore. The averaged energy landscapes are characterized by localized minima and reduced basin connectivity, consistent with local ion stabilization within the transmembrane pore. Phosphorylation does not generate a clearly continuous pathway for Cd^2+^, supporting reduced transport compatibility relative to monovalent substrates.

The structural panels shown on the right of Fig. 7 support this interpretation. Cd^2+^ positions remain concentrated within limited regions of the transmembrane domain, particularly in comparison with the more extended distributions observed for Ag^+^ and Cu^+^. This spatial restriction is consistent with stronger local coordination of the divalent ion. Because Cd^2+^ has a higher charge density than the monovalent ions, interactions with suitable coordinating residues can impose a stronger energetic preference for particular positions within the pore. In this context, the localized minima observed in the FES may reflect local trapping or stabilization of Cd^2+^rather than efficient progression through the transmembrane pathway.

At the residue level, Cd^2+^ interactions are more persistent and restricted than those observed for monovalent ions. The ion tends to remain coordinated to a smaller set of residues (Fig. S14), reflecting stronger local electrostatic and coordination interactions. In plant homologues, Cd^2+^ may display some positional variability within the pores, but this local mobility does not necessarily correspond to a continuous transport pathway. Instead, it suggests that increased structural plasticity can allow local exploration without overcoming the energetic constraints associated with divalent ion translocation.

Overall, the averaged FES indicates three ion-dependent energetic regimes. Cu^+^displays the most transport-compatible landscapes, with low-energy regions connected in the systems. Ag^+^ remains accessible to pores but shows a more variable landscape, consistent with partial compatibility and reduced substrate specificity. Cd^2+^ shows localized minima and limited connectivity, consistent with a less productive translocation regime. These trends suggest that the selectivity of ions in these P_1*B*_-ATPases arises from the combined effects of ion charge, coordination chemistry, conformational remodeling dependent on phosphorylation, and the plasticity of each transmembrane pore.

The comparison between bacterial and plant homologs further suggests that the basic architecture of ion passage is conserved, but the energetic response to ion identity and phosphorylation is system-dependent. LpCopA provides a more defined reference pathway, particularly for Cu^+^, while OsHMA9 and AtHMA5 display wider and more flexible sampling patterns. This increased plasticity may be relevant for plant metal transporters, which operate under variable metal availability and participate in both detoxification and metal distribution. Together, the averaged FES, FES decomposition, and structural ion-mapping analyzes suggest that ion translocation may be influenced not only by the presence of suitable coordinating residues, but also by the maintenance of accessible low-energy regions along the transmembrane pathway.

## 3 Methods

### 3.1 Molecular Modeling and System Preparation

The atomic structure of *Legionella pneumophila* copper-transporting P-type ATPase (LpCopA) in the dephosphorylated state was constructed from the Protein Data Bank (PDB ID: 3RFU; 3.2 Å resolution), which corresponds to the E2·Pi conformation [14]. The phosphorylated conformation of LpCopA in the E2P state was also obtained from the PDB (ID: 4BBJ) [10]. For the construction of *Oryza sativa* heavy metal ATPase (OsHMA9), encoded by the HMA9 gene, the protein sequence was obtained from UniProt (ID: A0A0P0×004) [41] and the *Arabidopsis thaliana* copper-transporting ATPase AtHMA5 (UniProt ID: Q9SH30) [42] were predicted using AlphaFold2 (Fig. S1-2) [43] with the LpCopA crystal structures used as templates for structural alignment. HMA9 (OsHMA9) and HMA5 (AtHMA5) were selected due to their biological relevance in metal homeostasis and detoxification in plants, while their conformational dynamics dependent on phosphorylation, inter domain communication patterns, and transport-associated structural responses remain comparatively underexplored at atomistic resolution [41, 44, 45].

Both LpCopA systems were embedded in a 1,2-dioleoyl-sn-glycero-3-phosphocholine (DOPC) lipid bilayer following the protocol described by [10], while the OsHMA9 and AtHMA5 systems were inserted into 1-palmitoyl-2-oleoyl-sn-glycero-3-phosphocholine (POPC) bilayers. All systems were solvated in a triclinic simulation box filled with TIP3P water molecules and system charges neutralized by adding counter ions. All topologies were constructed employing the CHARMM36m force field [46] parameters within CHARMM-GUI Membrane Builder [47].

### 3.2 Molecular Dynamics Simulations

Molecular dynamics simulations were performed using GROMACS [48]. Each system was subjected to steepest-descent energy minimization, followed by one NVT equilibration stage and five successive NPT stages of 5 ns each. Positional restraints applied to the protein and membrane were progressively reduced during equilibration.

Production simulations were performed for 1 *µ*s using a 2 fs integration time step. Three independent replicas were generated for each protein and phosphorylation state. Long-range electrostatic interactions were treated using the particle mesh Ewald method [49], with a real-space cutoff of 1.2 nm. Lennard-Jones interactions were switched off between 1.0 and 1.2 nm. Temperature was maintained at 303.15 K using the stochastic velocity-rescaling thermostat [50], and pressure was maintained at 1 bar using semi-isotropic Parrinello–Rahman coupling [51]. Additional simulation parameters, including metal-ion parameters (Table S1) and software versions are provided in the Supporting Information.

#### 3.2.1 Secondary Structure Analysis (DSSP)

The evolution of the secondary structure of proteins along the MD trajectories was evaluated using the Define Secondary Structure of Proteins (DSSP) algorithm [24]. Analyzes were performed with the DSSP tool implemented in GROMACS [48], which assigns secondary structure elements based on hydrogen-bonding patterns and backbone geometry. For each system, DSSP calculations were carried out over the full production trajectories of all replicas. The resulting secondary structure assignments were used to monitor the stability of key structural elements, including transmembrane helices and cytosolic domains (A, N, and P domains). To facilitate comparison between systems and phosphorylation states, secondary structure profiles were analyzed as a function of both the simulation time and the residue index. Particular attention was given to regions involved in ion transport and interdomain coupling, where local structural rearrangements may influence gating and communication pathways. The DSSP output was further processed to generate secondary structure maps and persistence profiles, allowing the identification of stable versus transient structural motifs across the trajectories.

### 3.3 Enhanced-Sampling Simulations

Well-tempered metadynamics simulations were performed using GROMACS coupled to PLUMED [52, 53]. Separate metal-containing systems were prepared for Cu^+^, Ag^+^, and Cd^2+^. The initial metal positions were defined from the putative cytoplasmic pore-entry site proposed for LpCopA by Grønberg et al. [37] and from the corresponding structurally superposed regions in OsHMA9 and AtHMA5.

Two collective variables were used. CV1 was defined as the distance between the center of mass of the transported ion and the center of mass of a selected group located near the pore exit. CV2 described the relative displacement of three selected transmembrane groups and was defined as

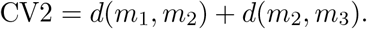

Detailed definitions of the collective variables, metal-ion placement, and biasing parameters are provided in the Supporting Information.

#### 3.3.2 FES decomposition analysis

The stability of the reconstructed metadynamics free-energy surfaces (FES) was evaluated through a time-resolved FES decomposition analysis. For each replica, the deposited HILLS file was divided into sequential time blocks and an independent FES was reconstructed for each block. The block-wise FES were inspected using the same collective variables, grid limits, and energy scale applied to the final averaged surfaces. This ensured that differences among blocks reflected changes in sampling rather than differences in the reconstruction or visualization procedure. We compared the block-wise FES to verify whether the main low-energy regions were consistently sampled over time. The FES decomposition was used to assess the temporal stability of the sampled pathways and to identify regions of the CV space that were explored more or less. This analysis was applied to all replicas and systems in order to provide a qualitative assessment of FES stability and sampling variability in the metadynamics simulations.

### 3.4 Ion–residue contact analysis

The contacts between each transported ion and the transmembrane residues were quantified by calculating the fraction of simulation frames in which the minimum distance between the ion and any heavy atom of a residue was less than 0.35 nm (3.5 Å) [54, 18]. This cutoff is in agreement with typical transition metal coordination geometries in proteins, where direct metal–ligand interactions are generally observed between approximately 2.0 and 2.7 Å [55]. The extended cutoff allows for the identification of both direct coordination and transient stabilizing interactions during ion transport. Such interactions are characteristic of P_1*B*_-type ATPases, where conserved cysteine, methionine, and histidine residues participate in metal coordination along the transport pathway [10, 56]. Contact frequencies were therefore used to identify key residues involved in ion translocation.

### 3.6 Principal Component Analysis

Principal component analysis (PCA) was performed using GROMACS tools [57, 48]. Before analysis, trajectories were corrected for periodic boundary conditions and aligned with a reference structure to eliminate global translational and rotational motions. Covariance matrices of C*α* atomic fluctuations were constructed from equilibrated trajectory segments and diagonalized to obtain eigenvalues and eigenvectors. The first principal components, corresponding to the largest collective motions were also analyzed and trajectory projections onto PC1 and PC2 were used to map the conformational space explored by each system and to compare sampling between phosphorylation states and replicas. Representative structures along the extremes of the principal components were examined to interpret the structural nature of the dominant motions, particularly interdomain rearrangements involving the A, P, and N domains and their coupling to the transmembrane region. The PCA results were interpreted in conjunction with RMSD and RMSF analyzes to distinguish increased flexibility from directional conformational transitions. Visualization and plotting were performed using Python scripts.

### 3.5 Dynamic Network Analysis

Dynamic network analysis was performed to characterize residue-residue communication pathways based on molecular dynamics trajectories. The networks were constructed using the NetworkView plugin implemented in VMD 1.9.4 [58]. Each residue was represented as a node, defined by its C_*α*_ atom, and edges were established between pairs of residues that satisfy both spatial proximity and correlated motion criteria. The input correlation matrices were computed using the CARMA software package [59], which evaluates atomic fluctuations and generates correlation data based on covariance from MD trajectories. Edge weights were defined as a function of the absolute value of residue-residue correlations, allowing the construction of weighted networks that reflect the strength of dynamical coupling. Community detection was carried out using the Girvan-Newman algorithm [60], which partitions the network into groups of residues that exhibit correlated motions. These communities provide a coarse-grained description of collective dynamics and interdomain communication, as previously described for biomolecular systems [54]. Critical nodes were identified on the basis of the centrality of betweenness, which quantifies the number of shortest paths that pass through a given node.

### 3.6 Structural Analysis and Visualization

The root mean square deviation (RMSD), the root mean square fluctuation (RMSF), and the domain-domain distances were calculated using GROMACS tools. Additional analyzes and plots were performed using custom Python and R scripts. Protein structures and trajectories were visualized and rendered using PyMOL 2.5.

## 4 Conclusions

In the present study, we compared the conformational behaviour of a bacterial and two plant P_1B_- type ATPases under dephosphorylated and phosphorylated conditions by combining microsecond-scale unbiased molecular dynamics with well-tempered metadynamics along collective variables describing metal-ion displacement and transmembrane rearrangement.

Across all six systems, the overall protein fold remained preserved throughout the simulations, indicating that the elevated backbone RMSD values primarily originated from large-scale rearrangements of the cytosolic domains rather than from structural unfolding. Phosphorylation was associated with an increased separation between the monitored N- and P-domain regions in LpCopA, whereas an equivalent response was not consistently observed in the two plant homologues. Together with the principal component and dynamic-network analyses, these results indicate that the conformational response to phosphorylation is homologue-dependent rather than uniformly conserved across the investigated P_1B_-ATPases.

The reconstructed free-energy landscapes projected onto the selected collective variables revealed differences in the extent and connectivity of the accessible regions of the transmembrane pathway among transporters, phosphorylation states, and metal ions. Cu^+^generally explored broader and more connected regions, whereas Ag^+^displayed a more variable distribution and Cd^2+^ preferentially occupied localized minima with limited connectivity. These patterns suggest ion-dependent differences in pore accessibility and local stabilization under the simulated conditions. However, because the results depend on the selected collective variables and on classical nonpolarizable descriptions of the metal ions, they should be interpreted as mechanistic hypotheses rather than as definitive determinants of substrate selectivity.

Overall, our findings provide a comparative atomistic framework connecting phosphorylation-dependent domain rearrangements with changes in the accessibility of the transmembrane pathway in bacterial and plant P_1B_-ATPases. The observed homologue- and ion-dependent landscapes identify structural regions and coordination environments that can be prioritized in future mutational, biochemical, and transport studies, thereby providing testable hypotheses for how conformational dynamics may contribute to metal-ion recognition and translocation in this transporter family.

## 5 Acknowledgments

The authors acknowledge the Laboratório Nacional de Computação Científica (LNCC) for providing computational resources through the Santos Dumont supercomputer. This work was supported by Coordenação de Aperfeiçoamento de Pessoal de Nível Superior (CAPES), Conselho Nacional de Desenvolvimento Científico e Tecnológico (CNPq), and Fundação de Amparo à Pesquisa do Estado do Rio Grande do Sul (FAPERGS).

During the preparation of this manuscript, the authors used OpenAI’s ChatGPT (GPT-5.6) to assist with drafting and revising selected passages, English-language editing and clarification of methodological descriptions. All AI-generated or AI-assisted content was critically reviewed, edited, and verified by the authors.

