## Supplementary material for "Molecular Determinants of Ion Selectivity in Heavy-Metal P-type ATPases"

### 1 Supplementary Information

#### 1.1 Structural Models and Residue Modifications

OsHMA9 and AtHMA5 models were generated using ColabFold v1.6.2, which implements AlphaFold2 with MMseqs2-based multiple-sequence alignment generation. Model confidence was assessed using predicted aligned error (PAE) matrices, multiple-sequence alignment coverage and per-residue predicted local distance difference test (pLDDT) scores, as shown for OsHMA9 and AtHMA5 in Figures S1 and S2, respectively. The LpCopA crystal structures (PDB IDs 3RFU and 4BBJ) were used as input templates in ColabFold and also for post-prediction structural superposition.

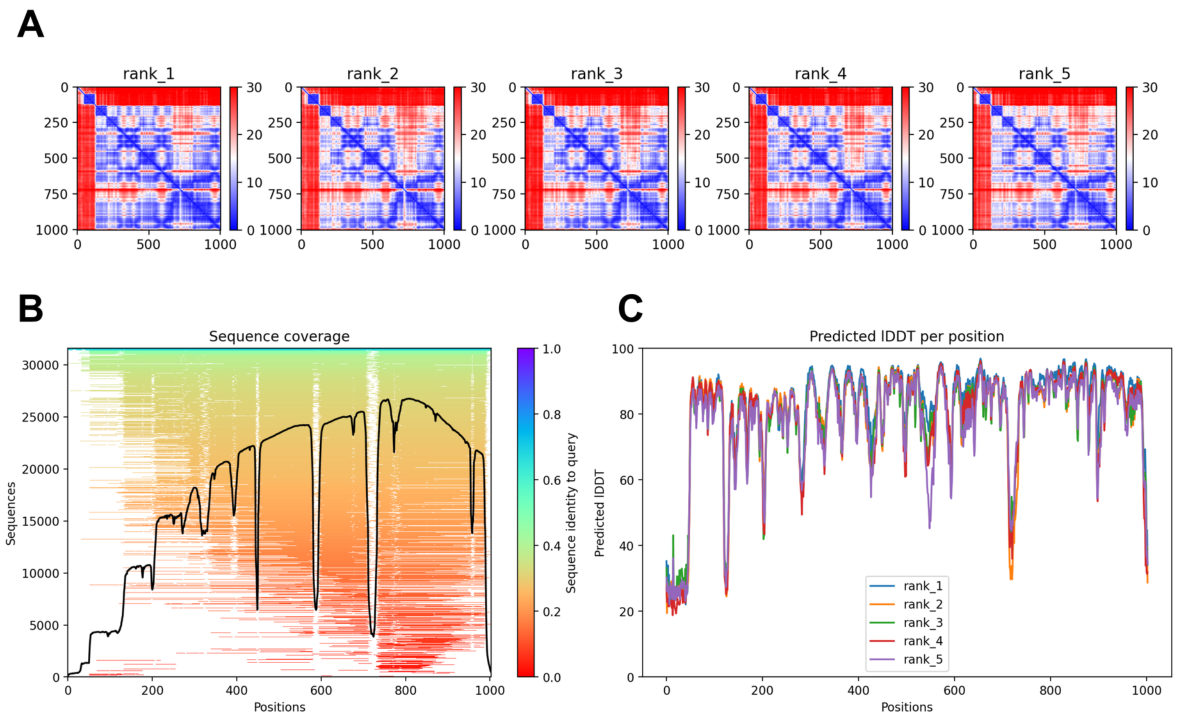


**Figure S1:** Confidence assessment of the AlphaFold2/ColabFold models generated for OsHMA9. **(A)** Predicted aligned error (PAE) matrices for the five ranked models. Lower PAE values indicate greater confidence in the relative positioning of residue pairs. **(B)** Multiple-sequence alignment coverage across the OsHMA9 sequence. The black line represents the number of aligned sequences at each position, while the color scale indicates sequence identity to the query sequence. **(C)** Per-residue predicted local distance difference test (pLDDT) scores for the five ranked models. Higher pLDDT values indicate greater confidence in the local structural prediction.

The crystallographic $AlF_4^-$ group and the associated $Mg^{2+}$ ion were removed from the 3RFU-derived model. In the LpCopA-P model derived from 4BBJ, $BeF_3^-$ was removed, whereas $Mg^{2+}$ was retained. Asp426 was maintained as phosphoaspartate. In the plant models, the conserved catalytic aspartate located in the phosphorylation (P) domain was modeled as phosphoaspartate using CHARMM-GUI.

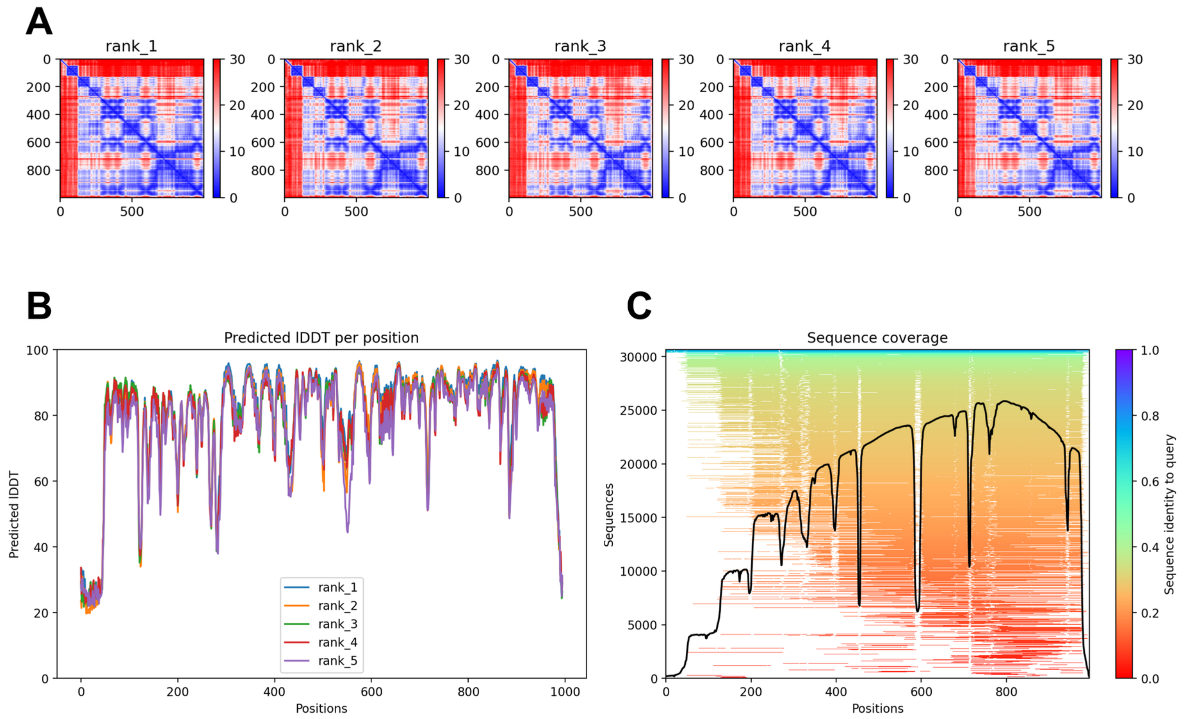


**Figure S2:** Confidence assessment of the AlphaFold2/ColabFold models generated for AtHMA5. **(A)** Predicted aligned error (PAE) matrices for the five ranked models. Lower PAE values indicate greater confidence in the relative positioning of residue pairs. **(B)** Per-residue predicted local distance difference test (pLDDT) scores for the five ranked models. Higher pLDDT values indicate greater confidence in the local structural prediction. **(C)** Multiple-sequence alignment coverage across the AtHMA5 sequence. The black line represents the number of aligned sequences at each position, while the color scale indicates sequence identity to the query sequence.

#### 1.2 Software and Computational Environments

Molecular simulations were carried out using GROMACS installations available on local laboratory workstations and on the Santos Dumont supercomputer. GROMACS 2021.5 was used on the laboratory workstations, whereas calculations performed on Santos Dumont employed the PLUMED-enabled GROMACS 2024.3 installation.

Within each set of comparable simulations, the simulation protocol was kept unchanged across replicas. System-specific topologies were generated using the same force field and preparation procedure, and equivalent integration, nonbonded-interaction, temperature-coupling, and pressure-coupling settings were applied.

#### 1.3 Metadynamics Collective Variables and Parameters

Well-tempered metadynamics simulations were performed using GROMACS coupled to PLUMED. The simulations employed a 2 fs integration time step and were continued from previously equilibrated configurations without reassignment of atomic velocities. Each metadynamics production run was configured for up to 100 ns. Bonds involving hydrogen atoms were constrained using the LINCS algorithm, and the temperature was maintained at 300 K using the stochastic velocity-rescaling thermostat. Pressure was maintained at 1 bar using the Parrinello-Rahman barostat with isotropic coupling.

Two collective variables were biased. The first collective variable, CV1, was defined as the distance between the center of mass of the transported metal ion and the center of mass of the selected pore-end group:

$$\mathrm{CV1} = d(\mathrm{ion}, \mathrm{pore\ end}).$$

The second collective variable described changes in the relative arrangement of three selected transmembrane groups. Two distances were initially calculated,

$$d_2 = d(m_2, m_4)$$

and

$$d_3 = d(m_4, m_6),$$

and CV2 was defined as their sum:

$$\mathrm{CV2} = d_2 + d_3 = d(m_2, m_4) + d(m_4, m_6).$$

Here, $m_2$, $m_4$, and $m_6$ represent the centers of mass of the selected groups associated with the corresponding transmembrane helices. The exact residue and atom selections were defined in the GROMACS index files prepared individually for each protein system.

Gaussian hills with an initial height of 0.6 kJ $\mathrm{mol}^{-1}$ were deposited every 500 simulation steps, corresponding to 1.0 ps. Gaussian widths of 0.10 nm and 0.20 nm were applied to CV1 and CV2, respectively. A bias factor of 20 and a PLUMED reference temperature of 298 K were used.

To restrict sampling to the selected transport region, a lower-wall potential was applied to CV1 at 0.15 nm with a force constant of 2000 kJ $\mathrm{mol}^{-1}$ $\mathrm{nm}^{-2}$, whereas an upper-wall potential was applied at 2.8 nm with a force constant of 4000 kJ $\mathrm{mol}^{-1}$ $\mathrm{nm}^{-2}$. Both wall potentials employed a quadratic exponent.

The values of CV1, the individual $d_2$ and $d_3$ distances, CV2, the metadynamics bias, and the lower- and upper-wall bias contributions were written to the `COLVAR` file every 200 simulation steps, corresponding to 0.4 ps. Six independent metadynamics replicas were performed for each investigated system.

##### 1.4 Initial Metal-Ion Placement and Pre-equilibration

The molecular dynamics simulations were performed in the absence of metal ions, being the metal-containing systems prepared separately for the metadynamics simulations. The initial metal positions were defined based on the putative cytoplasmic pore-entry site ( Met148, Glu205, and Asp337) proposed for LpCopA by Grønberg et al. [1]. In the simulated LpCopA and LpCopA-P structures, this region comprised Met76, Glu133, and Asp265, corresponding to Met148, Glu205, and Asp337, respectively, in the crystallographic numbering. For both phosphorylated/dephosphorylated states, the Cartesian coordinates of one selected donor atom from each residue were averaged, and the metal ion was inserted at the resulting geometric center.

The corresponding pore-entry regions in OsHMA9 and AtHMA5 were identified by structural superposition of each plant model onto the LpCopA structure. For OsHMA9 and OsHMA9-P, the initial ion position was calculated from selected donor atoms of Met239, Glu289, and Asp426. For AtHMA5 and AtHMA5-P, the corresponding position was calculated from selected atoms of Met372, Asp560, and Gly622. The same placement procedure was applied to the phosphorylated and dephosphorylated forms of each homologue. The metal-containing systems were subsequently submitted to energy minimization and NVT equilibration before production of the metadynamics simulations.

##### 1.5 Metal-Ion Parameters

The transported metal ions were represented using nonbonded parameters compatible with the force-field description employed in the simulations. The atom types, formal charges, and Lennard-Jones parameters assigned to $Cu^{+}$, $Ag^{+}$, and $Cd^{2+}$ are summarized in Table S1.

**Table S1:** Nonbonded parameters used for the transported metal ions.

| Ion | Atom type | Charge ($e$) | $\sigma$ | $\epsilon$ |
|---|---|---|---|---|
| $Cu^{+}$ | `CU1P` | +1 | 0.153590939007 | 0.7748768 |
| $Ag^{+}$ | `AG1P` | +1 | 0.216363662688 | 1.2008080 |
| $Cd^{2+}$ | `CAD` | +2 | 0.241789912103 | 0.5020800 |

These parameters were used in the metal-containing systems prepared specifically for the metadynamics simulations.

##### 1.6 Free-Energy Surface Processing

Individual FESs were reconstructed from each HILLS file using PLUMED sum_hills. The surfaces were shifted by subtracting their respective minimum values and interpolated onto a common grid.

Replica-averaged surfaces were calculated over the grid region shared by all three selected from the six replicas of each system studied. Pointwise standard deviations were calculated to quantify between-replica variability. Time-resolved surfaces were reconstructed over successive trajectory blocks using identical grid limits and energy scales.

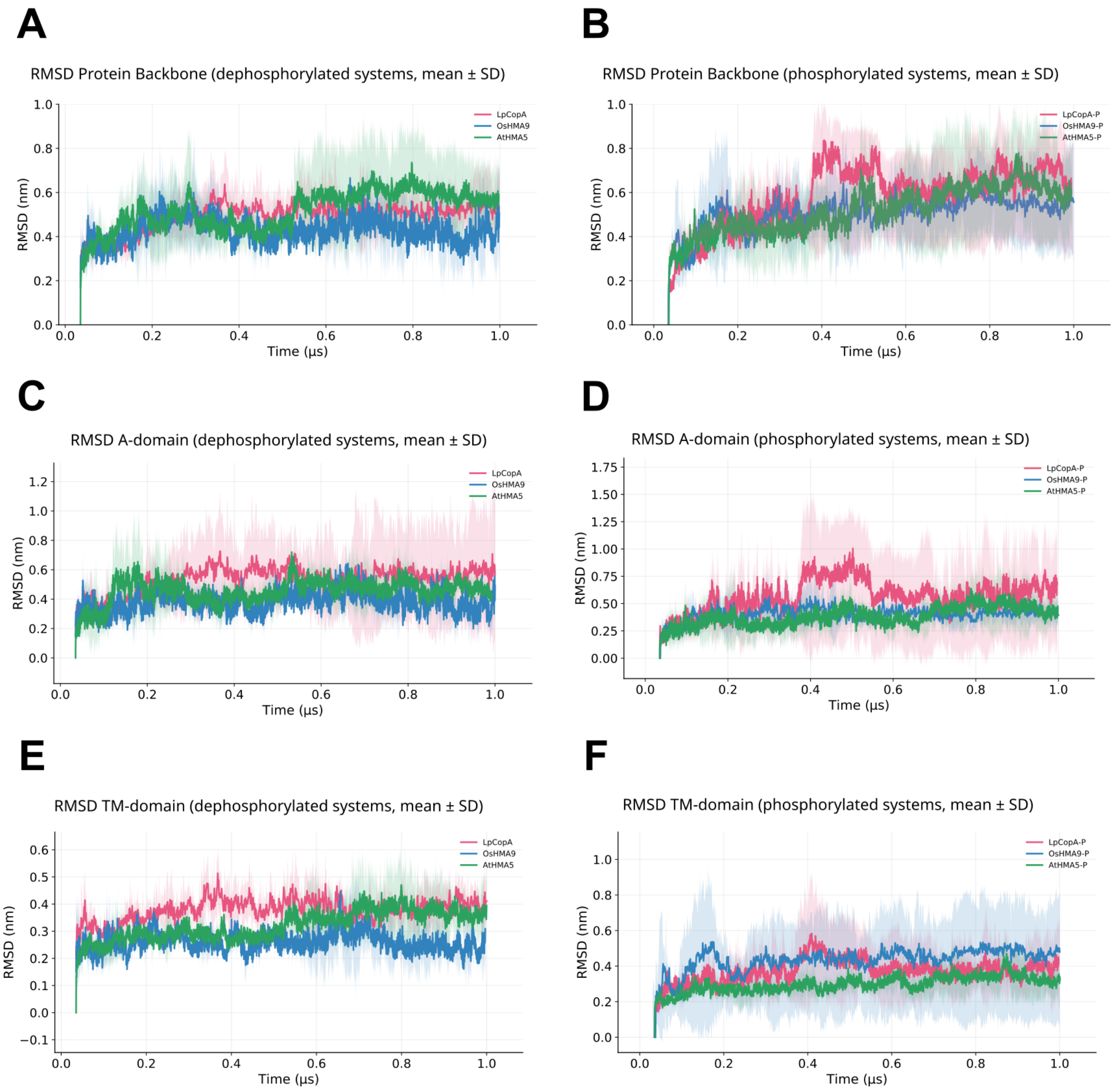


**Figure S3:** Backbone and domain-specific RMSD profiles for the $P_{1B}$-ATPase homologues, shown as mean ± standard deviation. Panels A, C, and E correspond to the dephosphorylated systems, while panels B, D, and F correspond to the phosphorylated systems. Panels A–B show the full protein backbone, C–D the A-domain, and E–F the transmembrane (TM) domain. The TM domain remains comparatively stable in all systems, whereas the A-domain exhibits higher variability. Phosphorylation leads to increased RMSD values and fluctuations, particularly in LpCopA, while plant homologues display more moderate changes.

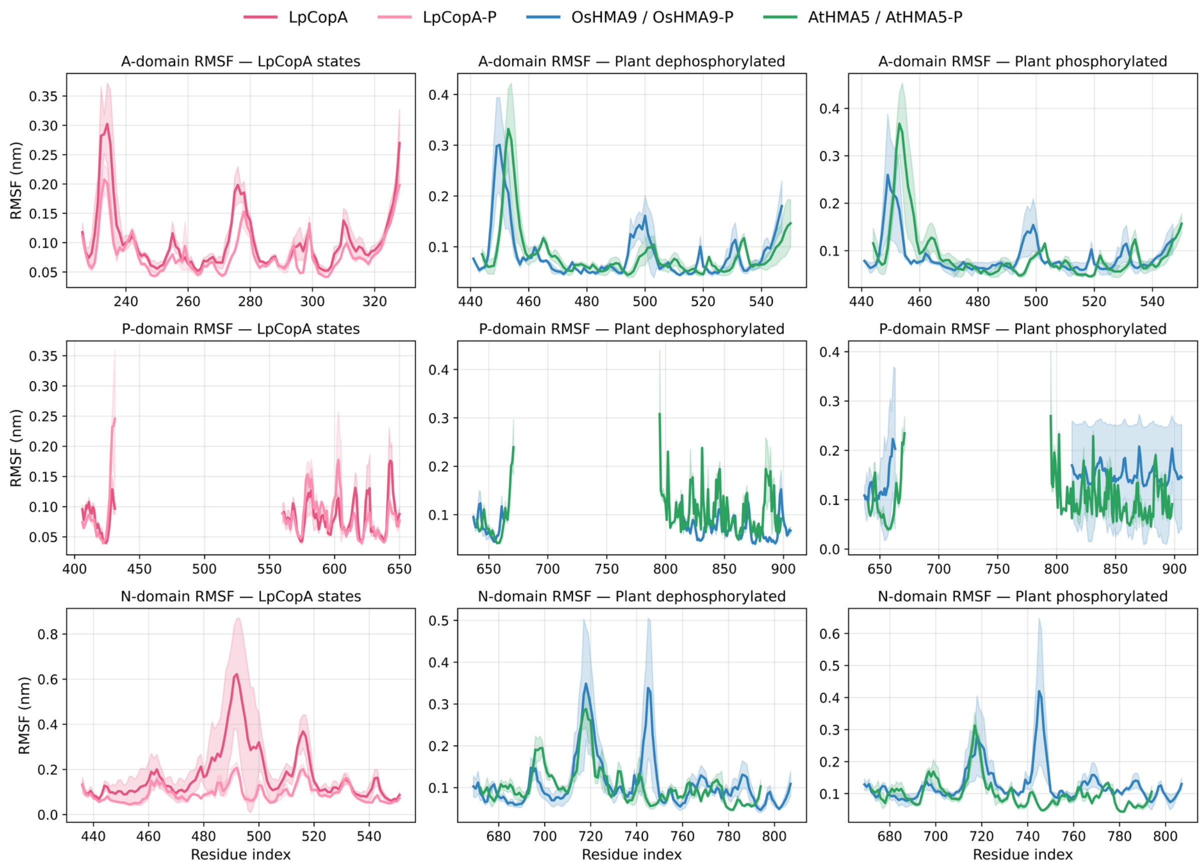


**Figure S4:** Residue-resolved root mean square fluctuation (RMSF) profiles for the cytosolic catalytic domains of $P_{1B}$-ATPases. The figure compares fluctuations of the actuator (A), phosphorylation (P), and nucleotide-binding (N) domains across bacterial and plant homologs in dephosphorylated and phosphorylated states. The left column shows the bacterial reference transporter LpCopA, including both the dephosphorylated (LpCopA) and phosphorylated (LpCopA-P) trajectories. The middle column shows the plant homologs OsHMA9 and AtHMA5 in the dephosphorylated state. The right column shows the same plant systems in the phosphorylated state. Rows correspond to the different cytosolic domains: **top row**, RMSF of the actuator (A) domain; **middle row**, RMSF of the phosphorylation (P) domain; **bottom row**, RMSF of the nucleotide-binding (N) domain. Solid lines represent the average RMSF across three independent molecular dynamics trajectories, while shaded regions correspond to the standard deviation, illustrating the variability between replicas. Color coding follows the convention used throughout the study: LpCopA (red), LpCopA-P (light red), OsHMA9/OsHMA9-P (blue), and AtHMA5/AtHMA5-P (green). The profiles highlight the heterogeneous distribution of flexibility across the catalytic headpiece. While the P-domain remains comparatively restrained in most systems, the A- and N-domains exhibit localized peaks associated with loop regions and interdomain hinge segments. Plant homologs generally display broader fluctuation patterns than the bacterial transporter, consistent with increased conformational plasticity within their cytosolic architectures.

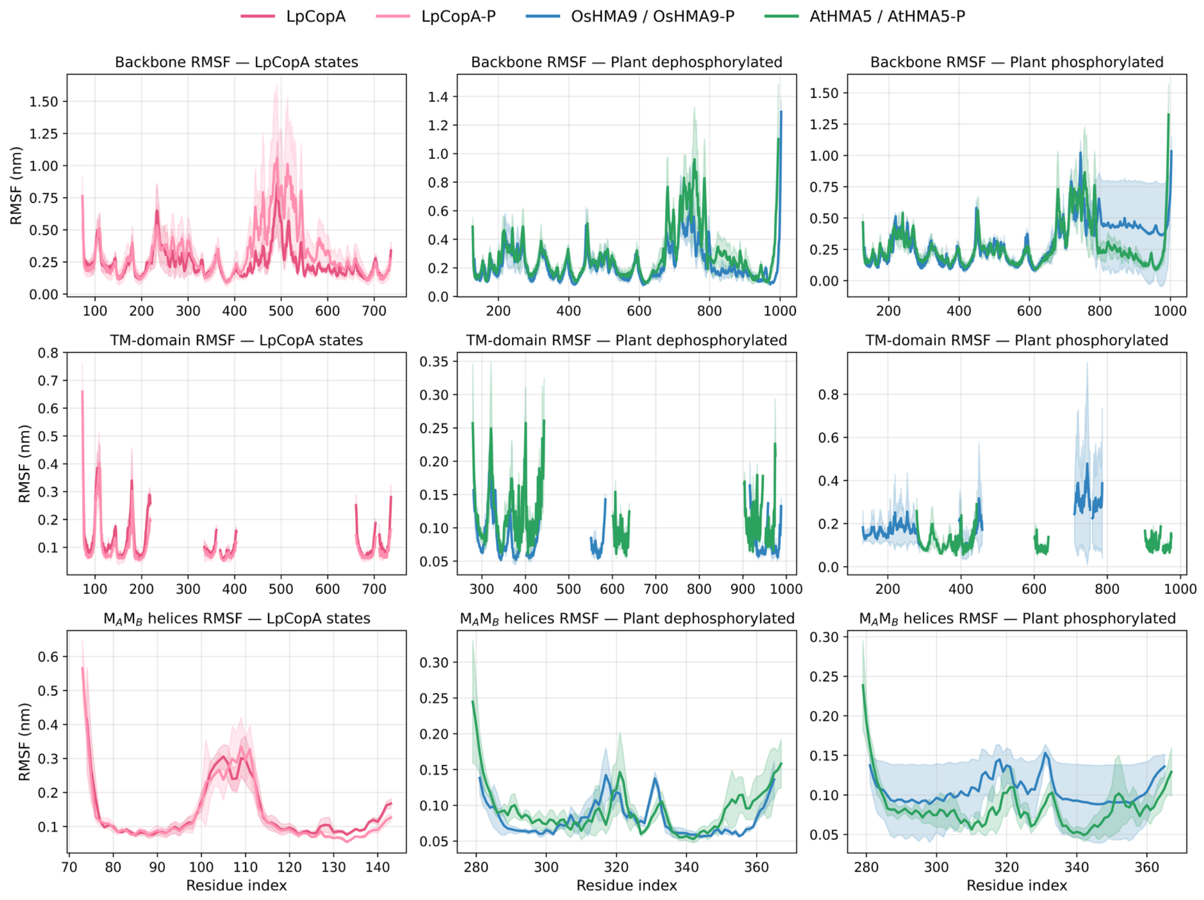


**Figure S5:** Backbone and transmembrane-region RMSF profiles for $P_{1B}$-ATPases. The figure summarizes residue-level fluctuations across the entire backbone, the transmembrane (TM) domain, and the metal-entry helices $M_A$ and $M_B$. The left column corresponds to the bacterial transporter LpCopA, including both the dephosphorylated (LpCopA) and phosphorylated (LpCopA-P) trajectories. The middle column shows the plant homologs OsHMA9 and AtHMA5 in the dephosphorylated state. The right column shows the same plant systems in the phosphorylated state. Rows correspond to different structural regions: **top row**, backbone RMSF calculated for the full protein; **middle row**, RMSF restricted to residues belonging to the transmembrane (TM) domain; **bottom row**, RMSF of the $M_A$ and $M_B$ helices, which form part of the metal-entry pathway. Solid lines represent the mean RMSF across three independent simulations, while shaded envelopes correspond to the standard deviation between replicas. Color coding is consistent with the rest of the study: LpCopA (red), LpCopA-P (light red), OsHMA9/OsHMA9-P (blue), and AtHMA5/AtHMA5-P (green). Across all systems, the transmembrane helices remain comparatively rigid relative to the cytosolic regions, consistent with their role as the structural scaffold for ion transport. In contrast, the backbone RMSF profiles reveal pronounced flexibility in peripheral loops and domain-connecting segments. The $M_A$ and $M_B$ helices show limited fluctuations overall but display localized differences between homologs, suggesting subtle variations in the dynamic behavior of the metal-entry region across bacterial and plant $P_{1B}$-ATPases.

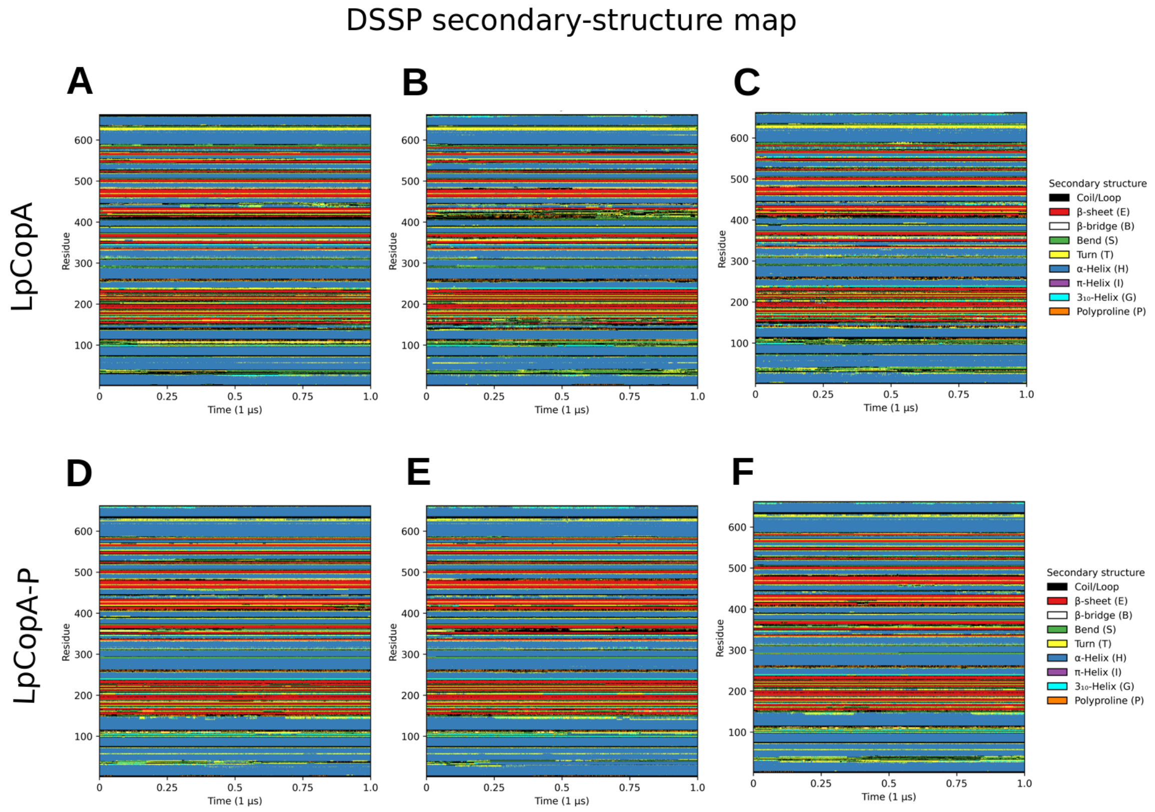


**Figure S6: Secondary-structure stability of LpCopA along 1 µs molecular dynamics simulations.** DSSP secondary-structure maps are shown as a function of simulation time (x-axis, 0–1 µs) and residue index (y-axis). Top row: non-phosphorylated LpCopA in three independent replicas (**A–C**). Bottom row: phosphorylated LpCopA (LpCopA-P) in three independent replicas (**D–F**). Across all systems and replicas, the overall secondary-structure pattern remains highly conserved throughout the trajectories. Extended $\alpha$-helical segments (blue), including the transmembrane helices, persist over time, while $\beta$-sheet elements (red) are maintained with only minor local fluctuations. Structural transitions are predominantly restricted to loop, bend, and turn regions (black/green/yellow), which are intrinsically flexible. No progressive loss of helicity or $\beta$-sheet content is observed, indicating that the elevated backbone RMSD values arise from domain rearrangements rather than global unfolding. The color code corresponds to DSSP assignments: coil/loop (black), $\beta$-sheet (E, red), $\beta$-bridge (B, white), bend (S, green), turn (T, yellow), $\alpha$-helix (H, blue), $\pi$-helix (I, purple), $3_{10}$-helix (G, cyan), and polyproline (P, orange).

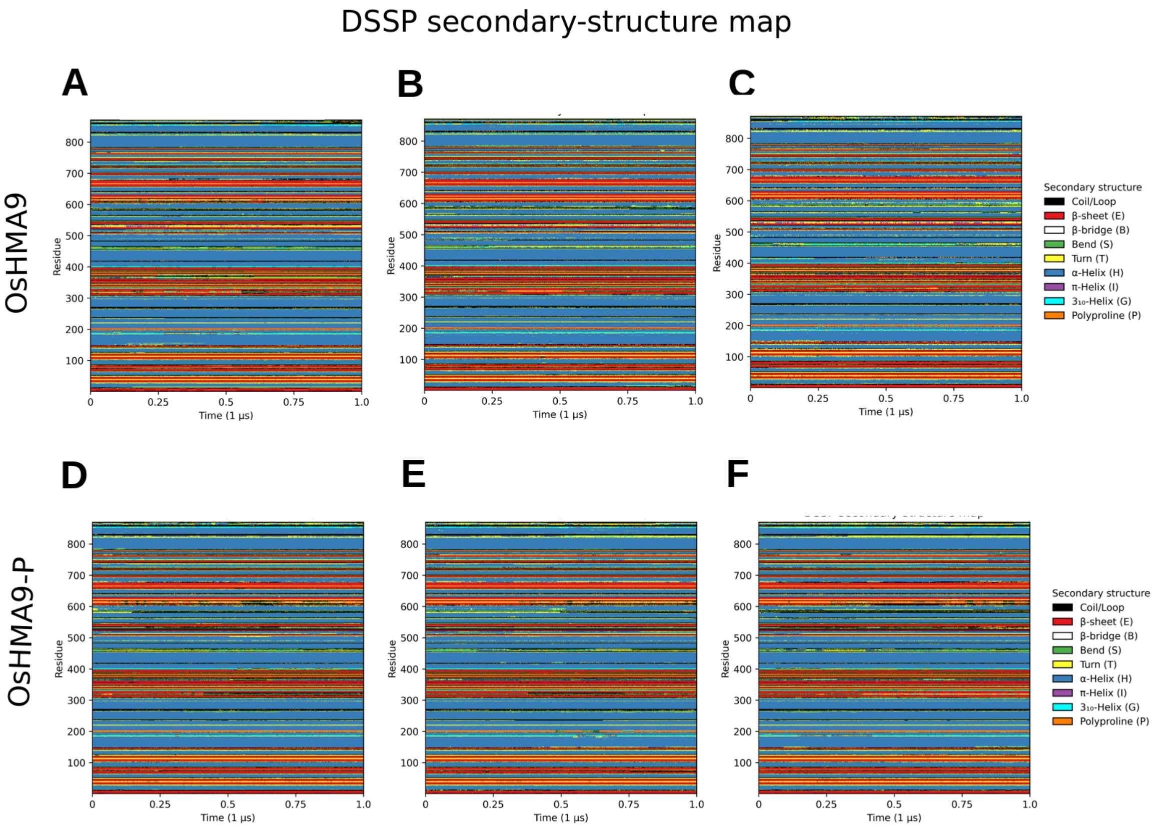


**Figure S7: Secondary-structure stability of OsHMA9 during 1 µs molecular dynamics simulations.** DSSP secondary-structure maps are displayed as a function of simulation time (x-axis, 0–1 µs) and residue index (y-axis). Panels **A–C** correspond to three independent replicas of the non-phosphorylated OsHMA9, while panels **D–F** represent three replicas of the phosphorylated form (OsHMA9-P). Across all replicas and states, the overall secondary-structure pattern remains highly conserved. Extended $\alpha$-helical regions (blue), including the transmembrane helices, persist throughout the trajectories, and $\beta$-sheet elements (red) remain stable. Transient fluctuations are primarily restricted to loop, bend, and turn regions, consistent with localized flexibility rather than global unfolding. The color scheme follows standard DSSP assignments: coil/loop (black), $\beta$-sheet (E, red), $\beta$-bridge (B, white), bend (S, green), turn (T, yellow), $\alpha$-helix (H, blue), $\pi$-helix (I, purple), $3_{10}$-helix (G, cyan), and polyproline (P, orange).

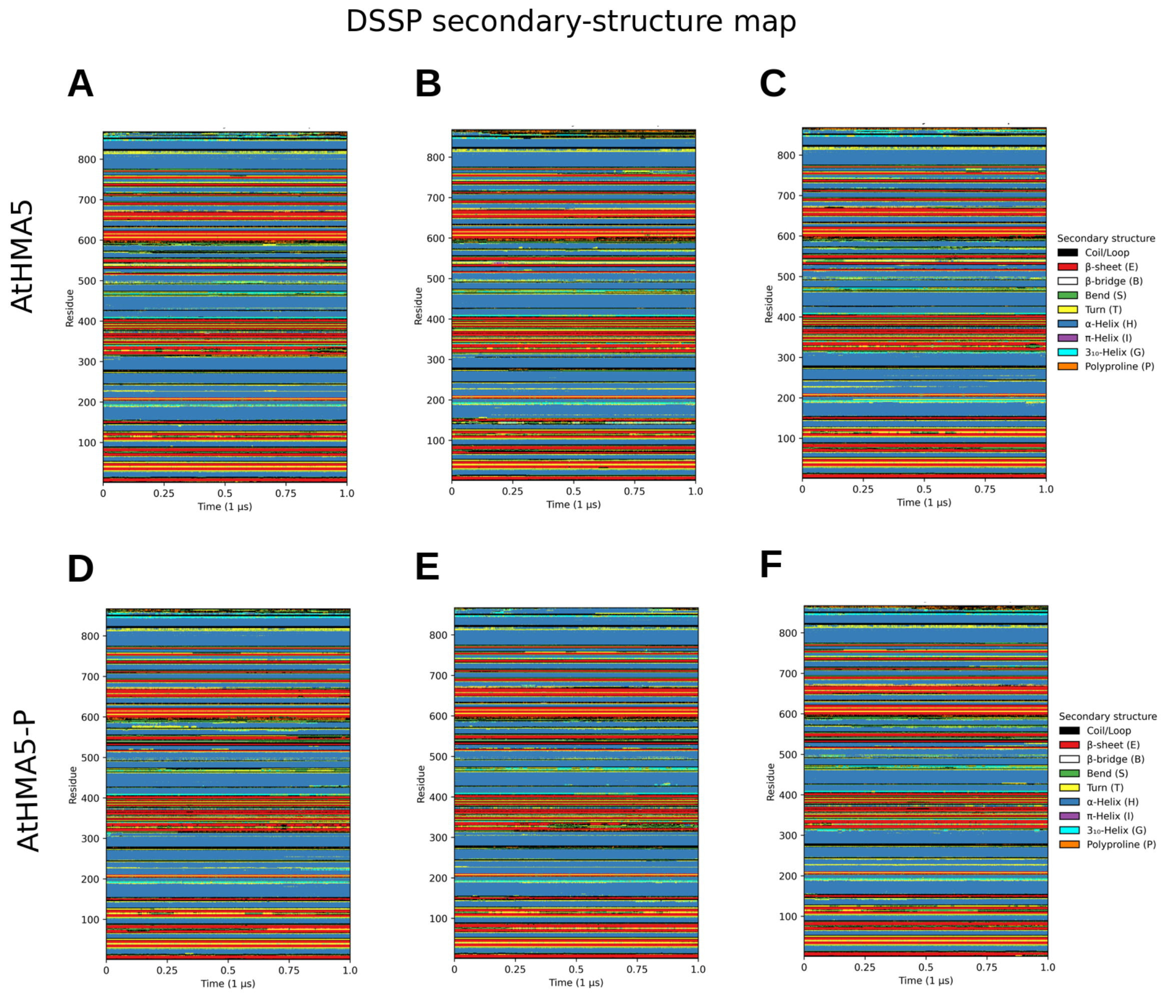


**Figure S8: Secondary-structure stability of AtHMA5 along 1 µs molecular dynamics simulations assessed by DSSP.** DSSP secondary-structure maps are shown as a function of simulation time (x-axis, 0–1 µs) and residue index (y-axis). Panels **A-C** correspond to three independent replicas of non-phosphorylated AtHMA5, whereas panels **D–F** correspond to three replicas of the phosphorylated form (AtHMA5-P). Across all replicas and phosphorylation states, the overall secondary-structure pattern remains highly conserved throughout the trajectories. Extended $\alpha$-helical segments (blue), including the transmembrane helices, persist over time, while $\beta$-sheet elements (red) are maintained with only minor local fluctuations. Structural transitions are predominantly confined to loop/coil, bend, and turn regions (black/green/yellow), consistent with localized flexibility rather than global unfolding. The color code follows DSSP assignments: coil/loop (black), $\beta$-sheet (E, red), $\beta$-bridge (B, white), bend (S, green), turn (T, yellow), $\alpha$-helix (H, blue), $\pi$-helix (I, purple), $3_{10}$-helix (G, cyan), and polyproline (P, orange).

**Table S2:** Mean distances between the centers of mass of selected residues in the N- and P-domain regions of the analyzed $P_{1B}$-ATPases. The mean distance was first calculated independently for each replica. Final values are reported as the mean ± sample standard deviation calculated from the three replica means.

| System | State | Replica 1 (nm) | Replica 2 (nm) | Replica 3 (nm) | Mean ± SD (nm) |
|---|---|---|---|---|---|
| LpCopA | Dephosphorylated | 2.3265 | 2.3763 | 2.3896 | $2.36 \pm 0.03$ |
| OsHMA9 | Dephosphorylated | 2.2112 | 1.8956 | 1.7917 | $1.97 \pm 0.22$ |
| AtHMA5 | Dephosphorylated | 2.0882 | 1.7684 | 2.0948 | $1.98 \pm 0.19$ |
| LpCopA-P | Phosphorylated | 3.0276 | 3.3561 | 2.6505 | $3.01 \pm 0.35$ |
| OsHMA9-P | Phosphorylated | 2.0027 | 1.9638 | 2.0859 | $2.02 \pm 0.06$ |
| AtHMA5-P | Phosphorylated | 2.1930 | 1.7259 | 2.1685 | $2.03 \pm 0.26$ |

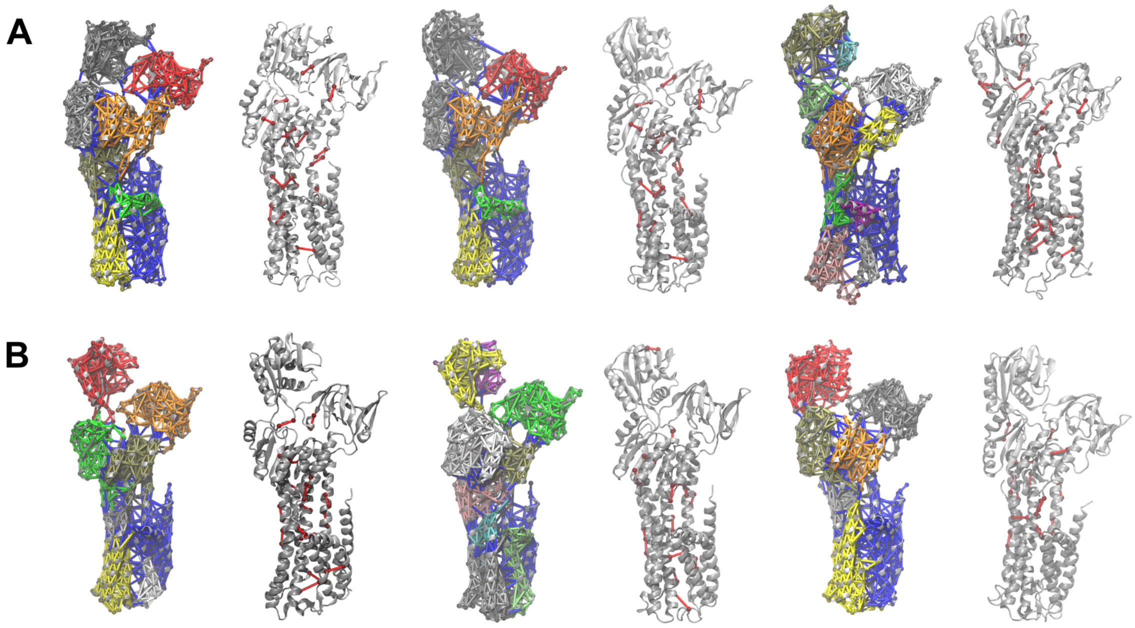


**Figure S9:** Dynamical network analysis of LpCopA systems obtained from molecular dynamics trajectories and computed using the NetworkView plugin in VMD. Column A corresponds to the dephosphorylated state (LpCopA), while column B represents the phosphorylated state (LpCopA-P). Colored network representations depict residue–residue communication pathways and community organization across the protein structure, highlighting interdomain coupling between cytosolic (A, N, P) and transmembrane regions. Structures shown in gray correspond to critical node analysis, identifying residues with high betweenness centrality that play key roles in long-range communication. All analyses were performed using full-length trajectories.

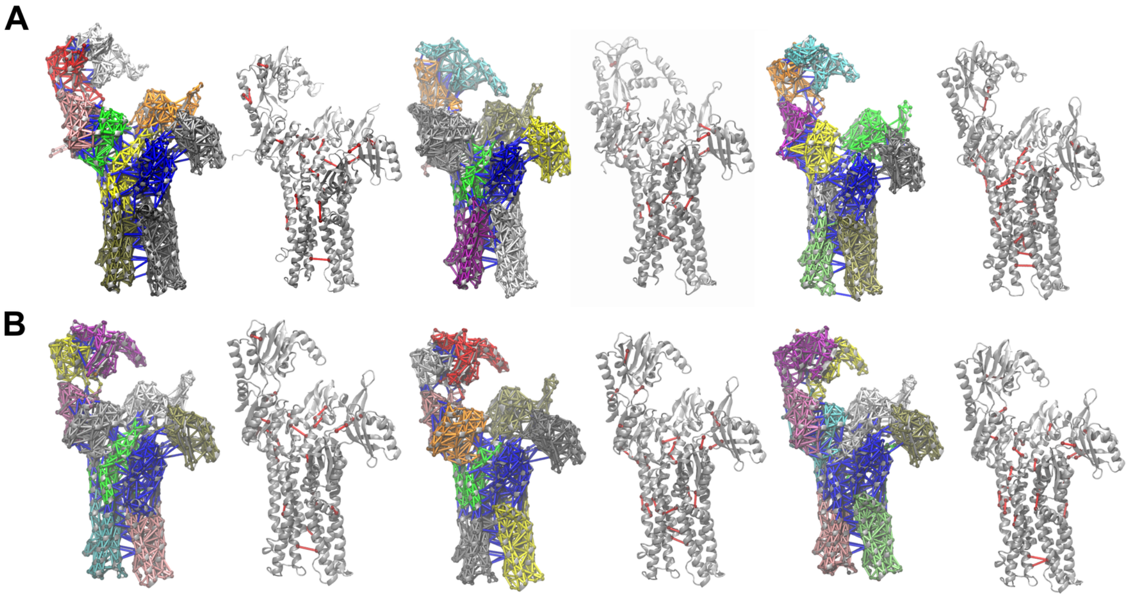


**Figure S10:** Dynamical network analysis of OsHMA9 systems derived from molecular dynamics simulations and evaluated using the NetworkView module in VMD. Column A shows the dephosphorylated state (OsHMA9), whereas column B corresponds to the phosphorylated state (OsHMA9-P). Colored network models represent residue interaction networks and community structures, illustrating communication pathways between cytosolic domains and the membrane-embedded region. Gray structures highlight critical nodes.

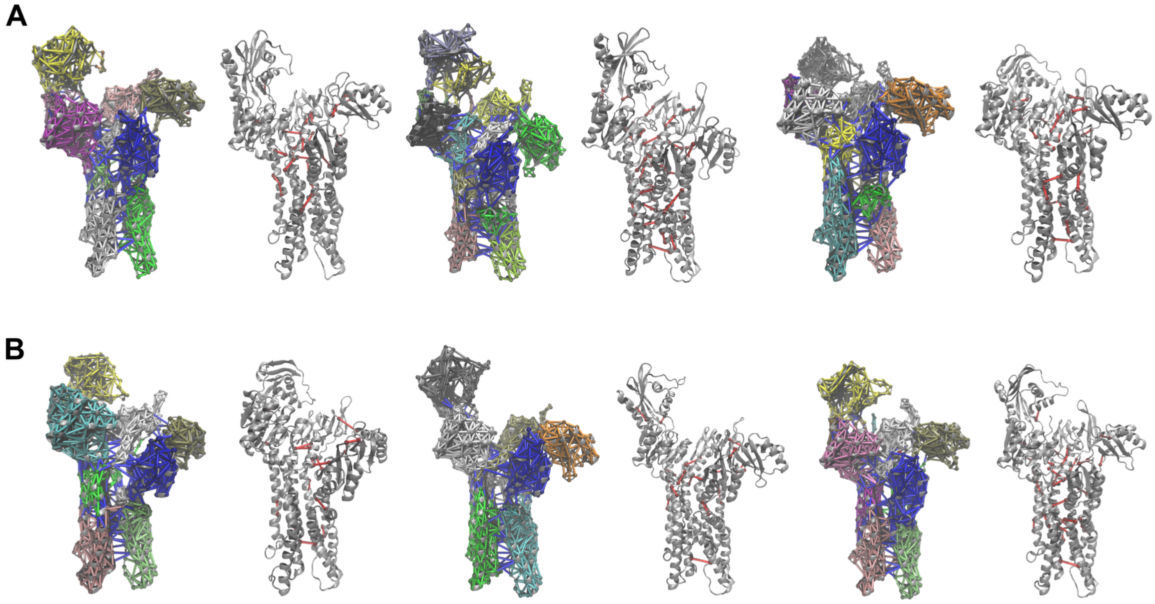


**Figure S11:** Dynamical network analysis of AtHMA5 derived from molecular dynamics simulations and visualized using the NetworkView module in VMD. Column A corresponds to the dephosphorylated state (AtHMA5), while column B represents the phosphorylated state (AtHMA5-P). Colored networks depict residue–residue interaction communities, highlighting pathways of communication between cytosolic domains and the transmembrane region. Gray representations indicate critical nodes identified from the network analysis.

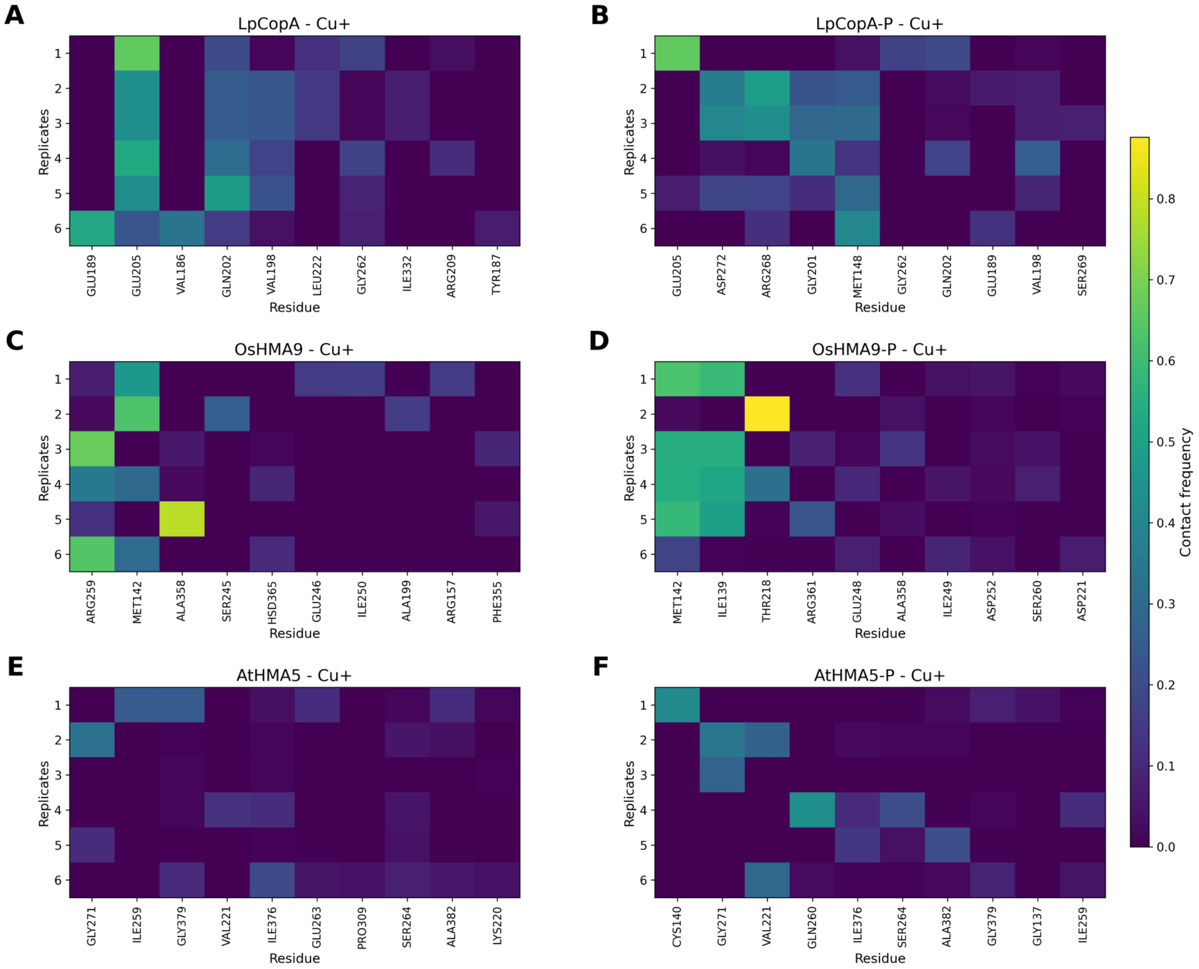


**Figure S12:** Replica-resolved residue–ion contact maps for $Cu^+$ transport in heavy-metal $P_{1B}$-type ATPases. Each heatmap displays the contact frequency between $Cu^+$ and selected channel residues across six independent molecular dynamics replicas (rows). Residues are ranked based on their average contact frequency within each system. Panels A, C, and E correspond to the dephosphorylated states of LpCopA, OsHMA9, and AtHMA5, respectively, whereas panels B, D, and F represent their phosphorylated counterparts (denoted as "-P"). Color intensity reflects the fraction of simulation frames in which the ion is within the contact cutoff distance of a given residue. Across all systems, $Cu^+$ interactions are dominated by a limited set of residues located along the transmembrane transport pathway, with consistent patterns observed across replicas, supporting the robustness of the identified ion conduction route.

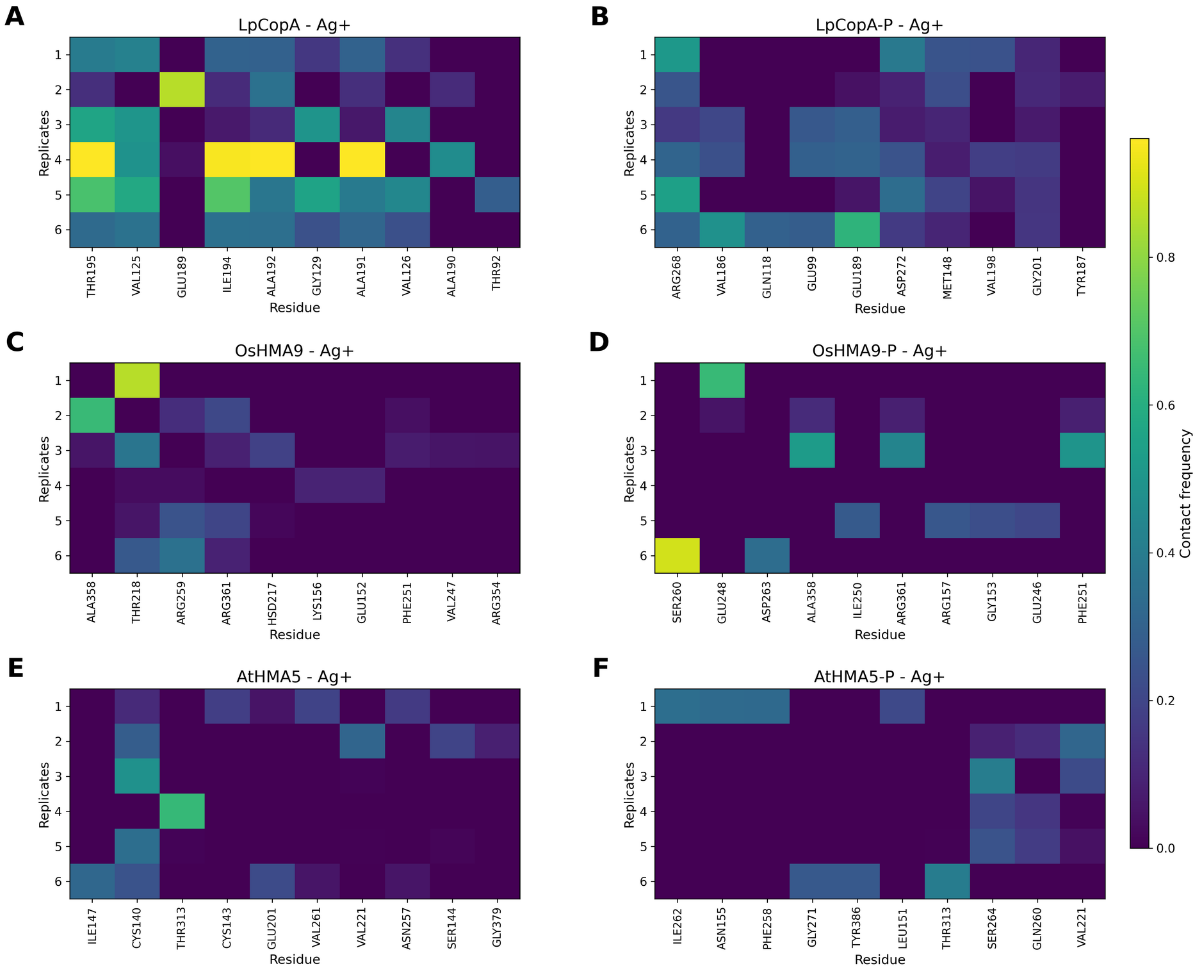


**Figure S13: Replicate-resolved residue–ion contact patterns during $Ag^+$ transport in heavy-metal $P_{1B}$-ATPases.** Heatmaps display the contact frequency between $Ag^+$ ions and protein residues across six independent replicas for each system. Panels correspond to: (A) LpCopA (dephosphorylated), (B) LpCopA-P (phosphorylated), (C) OsHMA9 (dephosphorylated), (D) OsHMA9-P (phosphorylated), (E) AtHMA5 (dephosphorylated), and (F) AtHMA5-P (phosphorylated). For each system, the top 10 residues ranked by average contact frequency across replicas are shown on the x-axis, while replicas are indicated on the y-axis. Color intensity represents the normalized contact frequency, with darker colors indicating low interaction probability and brighter colors indicating persistent or high-frequency contacts. Overall, $Ag^+$ transport exhibits distributed interaction patterns involving multiple residues along the transport pathway, with notable variability across replicas and between phosphorylation states.

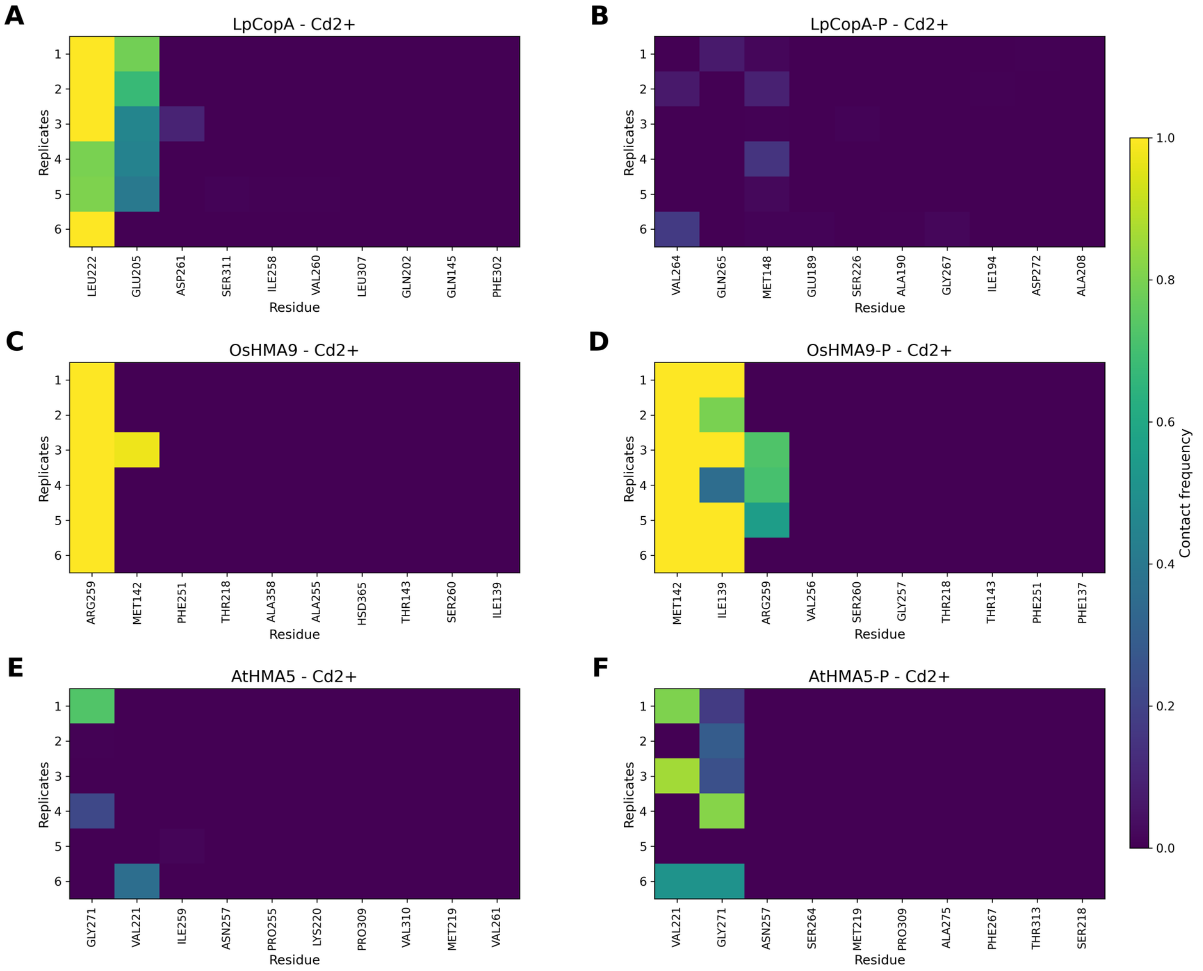


**Figure S14: Replicate-resolved residue–ion contact patterns during $Cd^{2+}$ transport in heavy-metal $P_{1B}$-ATPases.** Heatmaps show the contact frequency between $Cd^{2+}$ ions and protein residues across six independent replicas for each system. Panels correspond to: (A) LpCopA (dephosphorylated), (B) LpCopA-P (phosphorylated), (C) OsHMA9 (dephosphorylated), (D) OsHMA9-P (phosphorylated), (E) AtHMA5 (dephosphorylated), and (F) AtHMA5-P (phosphorylated). In contrast to $Cu^+$ and $Ag^+$, $Cd^{2+}$ interactions are highly localized, with only a small number of residues (typically one to two per replica) exhibiting significant contact frequency. These residues form stable, high-occupancy interactions, as evidenced by the concentrated high-intensity regions in the heatmaps. This behavior indicates that $Cd^{2+}$ transport is dominated by strong and persistent coordination with a limited set of residues, leading to reduced mobility along the transport pathway. The observed interaction pattern suggests a trapping-like mechanism, consistent with the higher charge density and stronger binding affinity of $Cd^{2+}$ compared to monovalent ions.

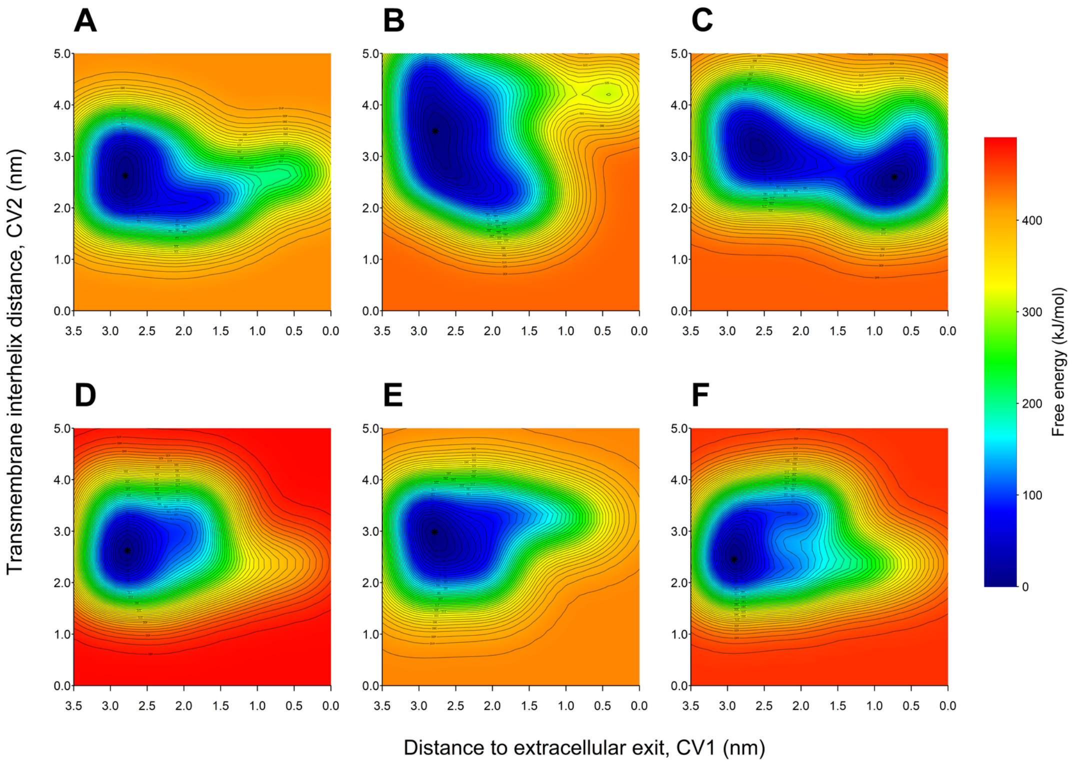


**Figure S15: Free energy surfaces (FES) for $Cu^+$ transport in dephosphorylated LpCopA across six independent replicas.** Two–dimensional free energy landscapes are projected onto the collective variables CV1 (distance to the extracellular exit) and CV2 (transmembrane interhelical distance). Panels A–F correspond to six independent metadynamics replicas of the same transport system. Energies are reported in kJ/mol and normalized such that the global minimum of each replica is set to zero. The color scale represents free energy values up to the global cap used for visualization consistency across replicas. The overall topology of the basins is reproducible across replicas, supporting convergence of the $Cu^+$ transport pathway in dephosphorylated LpCopA.

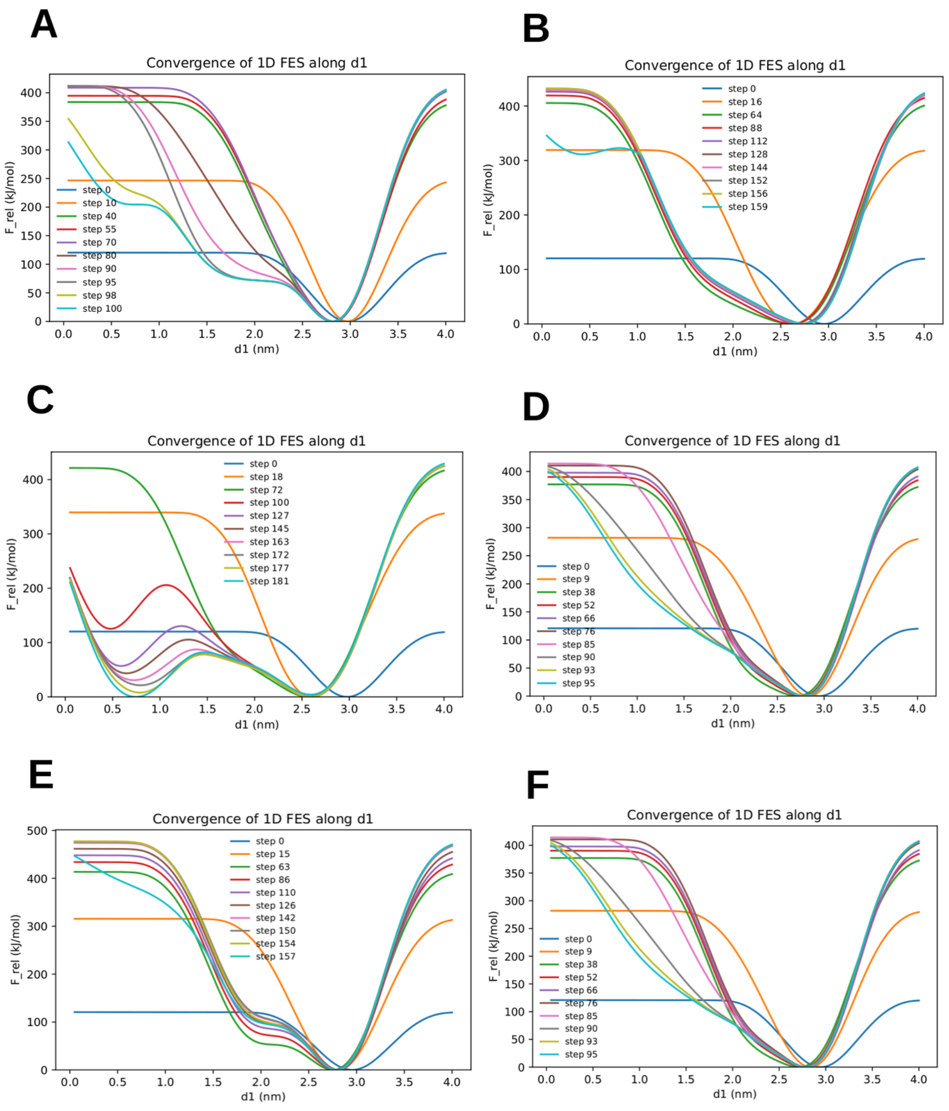


**Figure S16:** Convergence analysis of the one–dimensional free energy surface (FES) along the collective variable $d1$ for $Cu^+$ transport in **LpCopA**. Panels (A–F) correspond to six independent metadynamics replicas. Each colored curve represents the FES reconstructed at different stages of the simulation using accumulated bias potentials. A well defined minimum is consistently observed around $d1 \sim 2.7$–$3.0$ nm, corresponding to the energetically favorable coordination region of the ion along the transport pathway.

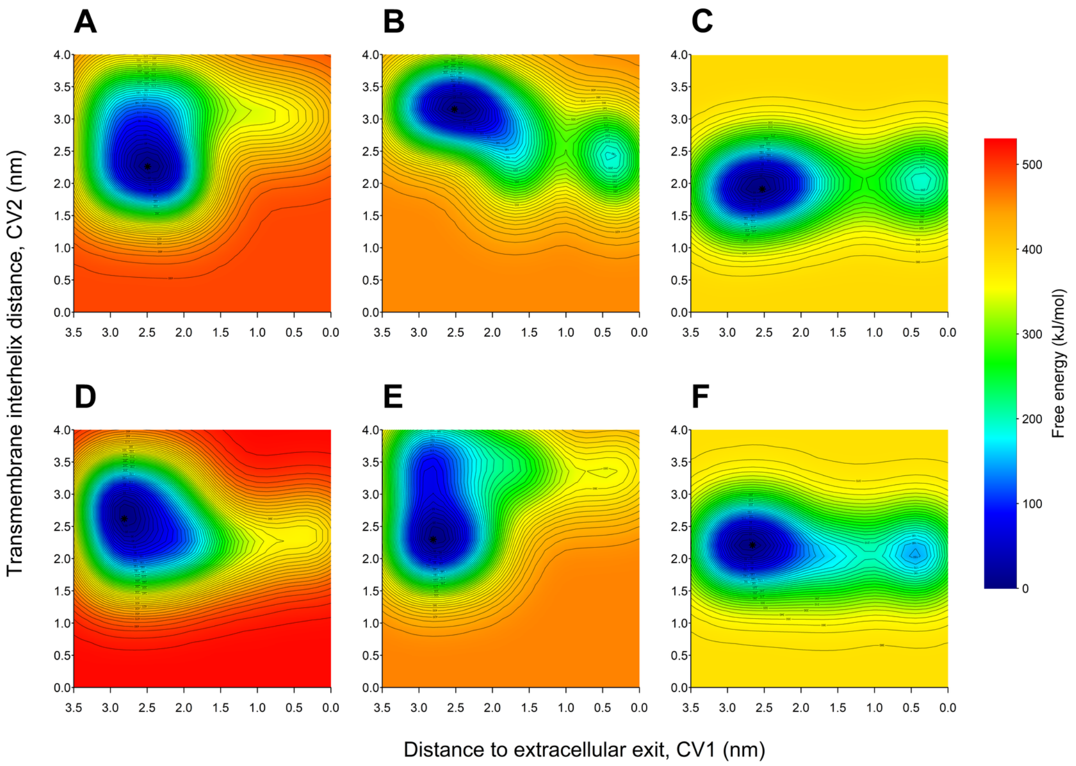


**Figure S17: Free energy surfaces of $Cu^+$ transport in phosphorylated LpCopA (LpCopA-P).** Two-dimensional free energy surfaces (FES) obtained from six independent replicas (A–F) of metadynamics simulations describing $Cu^+$ transport in phosphorylated LpCopA (LpCopA-P). The reaction coordinates correspond to the distance of the ion to the extracellular exit (CV1) and the transmembrane interhelical distance (CV2). Free energies are reported in kJ/mol and shifted so that the global minimum of each replica is set to zero. Black contour lines represent isoenergetic levels, and the star marks the location of the global minimum in each surface. Overall, the replicas display a consistent low-energy basin associated with the inward coordination region and a secondary minimum toward the extracellular side, indicating reproducible sampling of the transport pathway in the phosphorylated state.

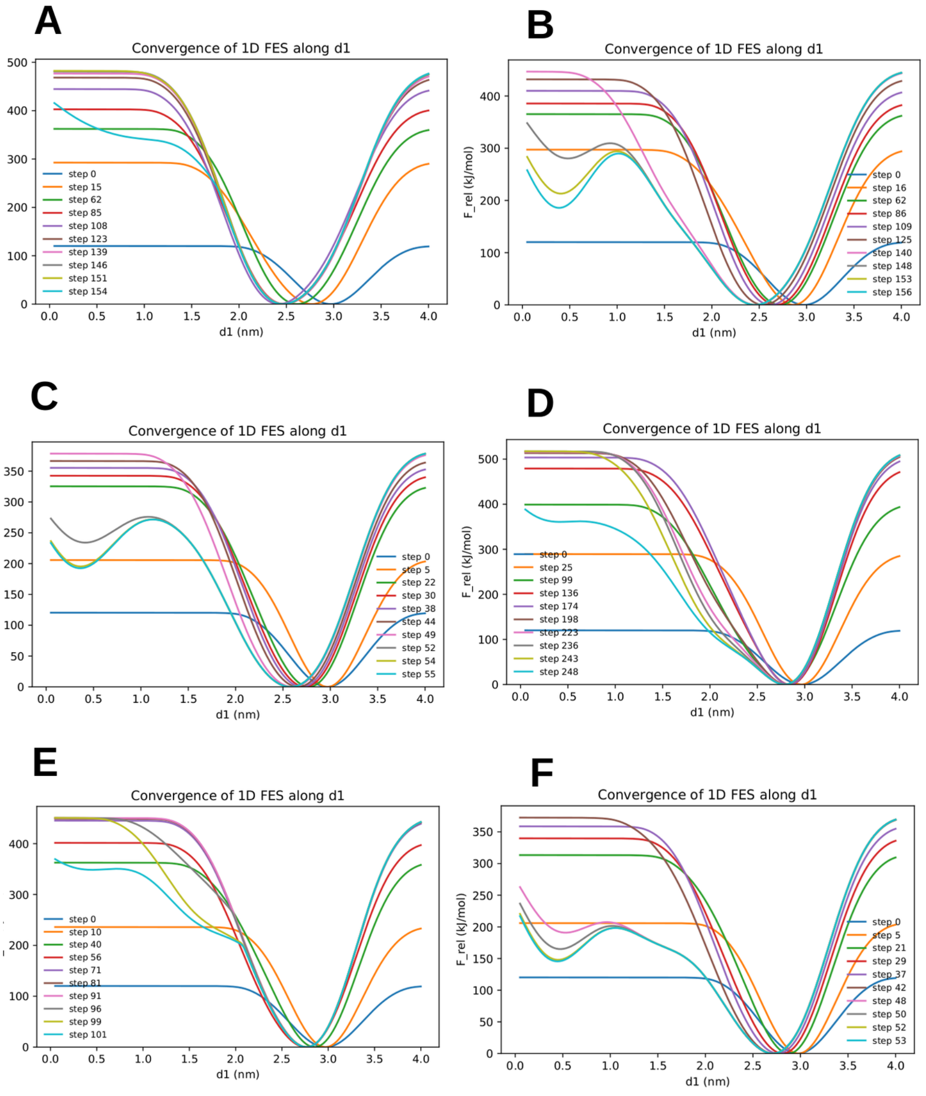


**Figure S18:** Convergence analysis of the one–dimensional free energy surface (FES) along the collective variable $d1$ for $Cu^{+}$ transport in the phosphorylated transporter **LpCopA-P**. Panels (A–F) correspond to six independent metadynamics replicas. Each colored curve represents the FES reconstructed at different stages of the simulation using accumulated bias potentials. As sampling progresses, the profiles progressively stabilize. A consistent free energy minimum emerges around $d1 \sim 2.7$–$3.0$ nm, corresponding to the preferred ion coordination region along the transport pathway.

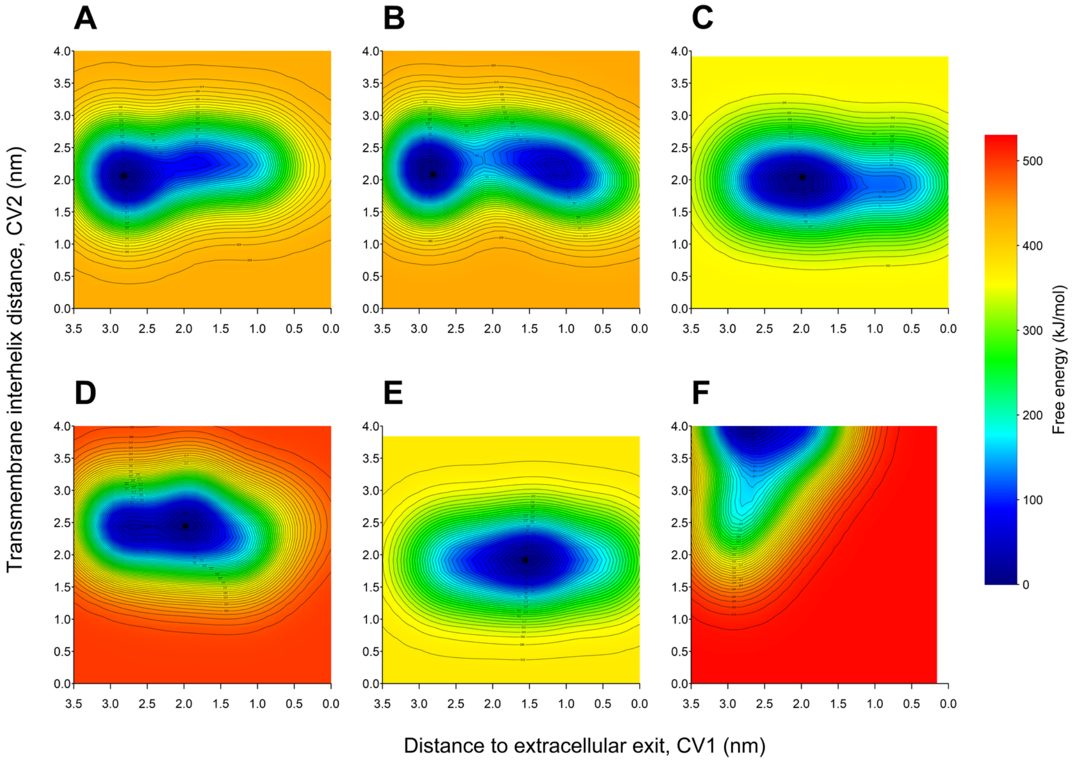


**Figure S19:** Two–dimensional free energy surfaces (FES) describing $Cu^+$ transport in dephosphorylated OsHMA9 obtained from six independent metadynamics replicas (A–F). The collective variables correspond to the distance to the extracellular exit (CV1) and the transmembrane interhelical distance (CV2). Free energies are shown in kJ/mol and are referenced to the global minimum of each replica. The overall topology reveals a dominant low-energy basin centered around CV2 $\sim$ 2.0–2.5 nm, with replica-dependent variations in basin width and extension along CV1, reflecting the intrinsic flexibility of the transport pathway in the dephosphorylated state.

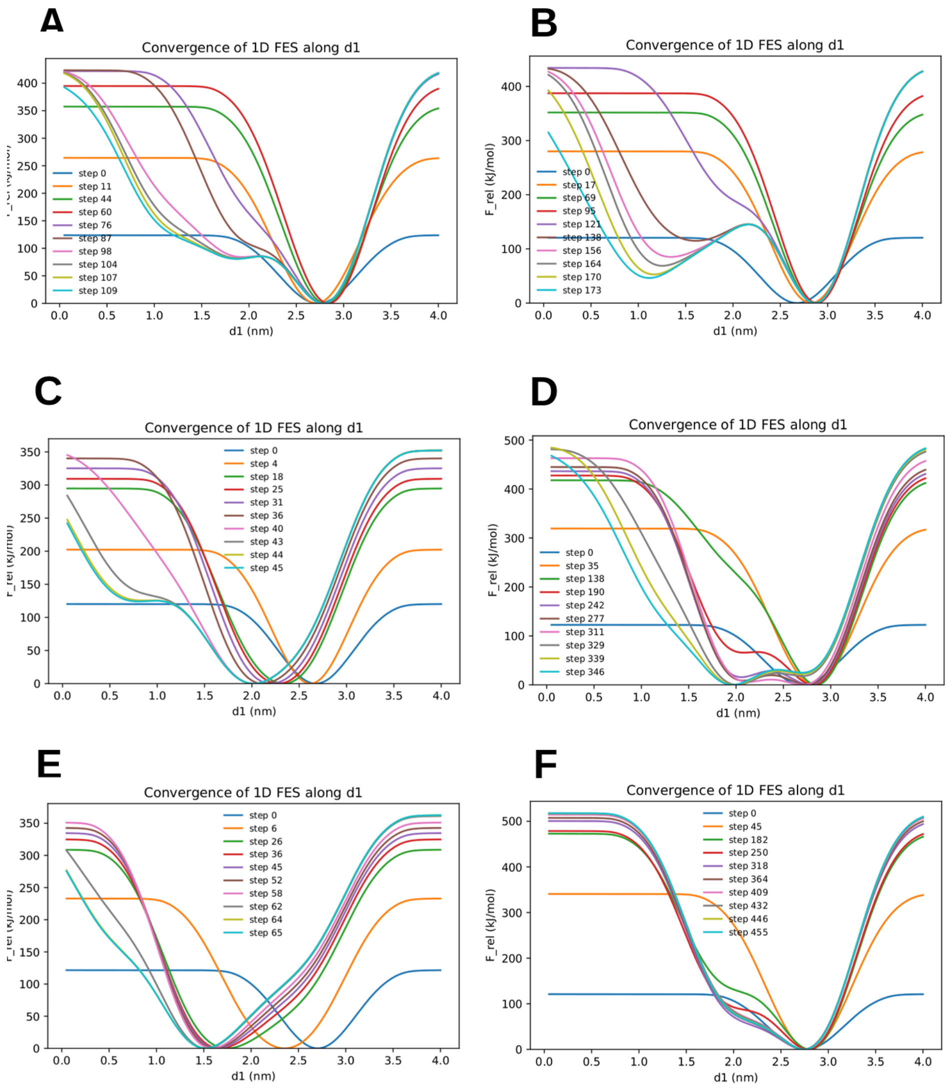


**Figure S20:** Convergence analysis of the one–dimensional free energy surface (FES) along the collective variable $d1$ for $Cu^+$ transport in **OsHMA9**. Panels (A–F) correspond to six independent metadynamics replicas. Each colored curve represents the FES reconstructed at different stages of the simulation using accumulated bias potentials. The progressive overlap of the profiles indicates stabilization of the free energy landscape as sampling proceeds. A dominant minimum appears around $d1 \sim 2.7$–$3.0$ nm, corresponding to the energetically favorable coordination region for the ion along the transport pathway.

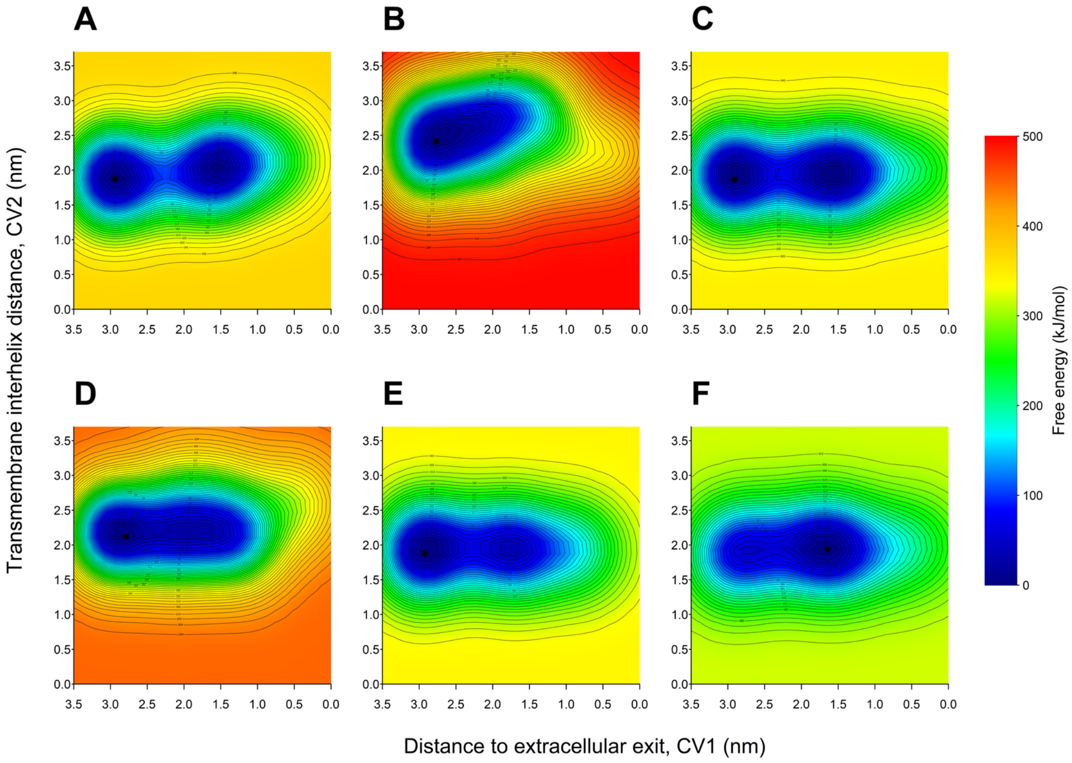


**Figure S21:** Two–dimensional free energy surfaces (FES) describing $Cu^+$ transport in phosphorylated OsHMA9 (OsHMA9-P) obtained from six independent metadynamics replicas (A–F). The collective variables correspond to the distance to the extracellular exit (CV1) and the transmembrane interhelical distance (CV2). Free energies are reported in kJ/mol and referenced to the global minimum of each replica. Across replicas, a dominant low-energy basin is observed within the transmembrane region, with variations in basin shape and extension along CV1 reflecting differences in sampling of intermediate ion coordination states in the phosphorylated conformation.

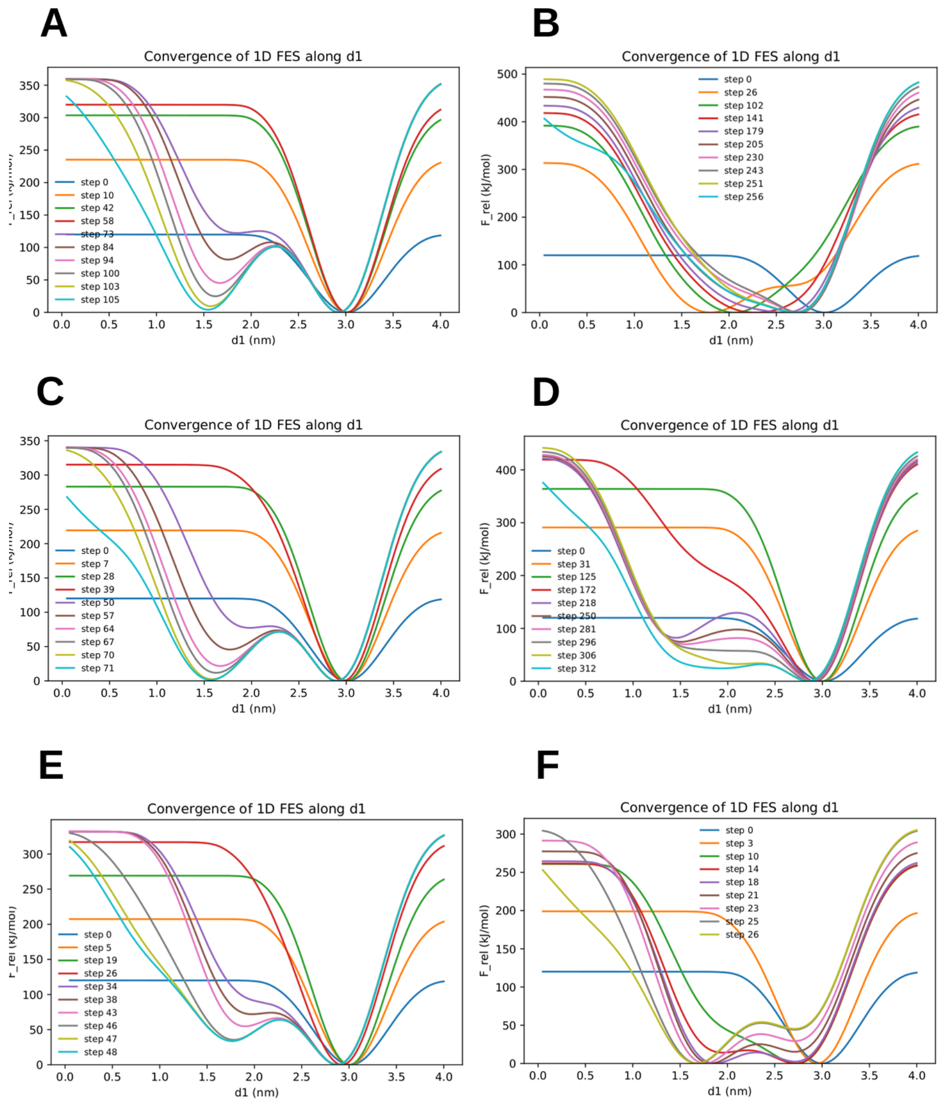


**Figure S22:** Convergence analysis of the one–dimensional free energy surface (FES) along the collective variable $d1$ for $Cu^+$ transport in the phosphorylated transporter **OsHMA9-P**. Panels (A–F) correspond to six independent metadynamics replicas. Each colored curve represents the FES reconstructed at successive stages of the simulations as bias potentials accumulate.

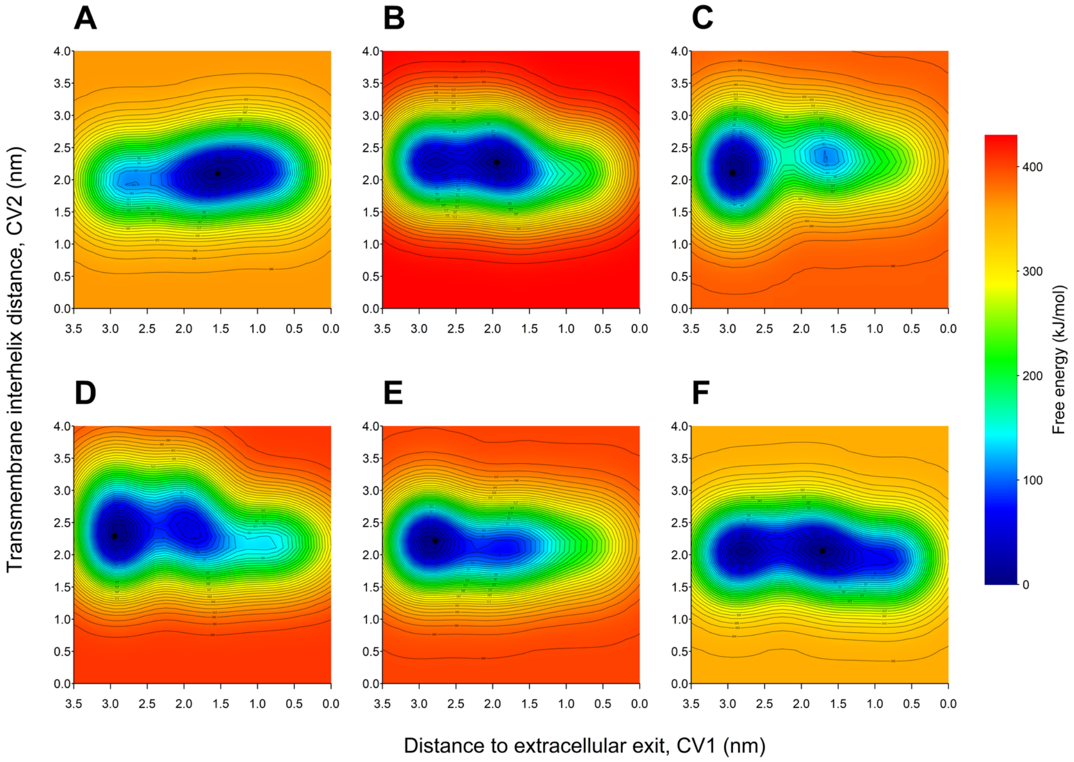


**Figure S23: Free energy surfaces of $Cu^+$ transport in dephosphorylated AtHMA5.** Two-dimensional free energy surfaces (FES) obtained from six independent replicas (A–F) of metadynamics simulations describing $Cu^+$ transport in the dephosphorylated state of AtHMA5, a $P_{1B}$-type ATPase from *Arabidopsis thaliana*. All simulations consistently reveal a dominant low-energy basin associated with the inward coordination region and a continuous free energy pathway toward the extracellular side, supporting a reproducible transport profile in the dephosphorylated state.

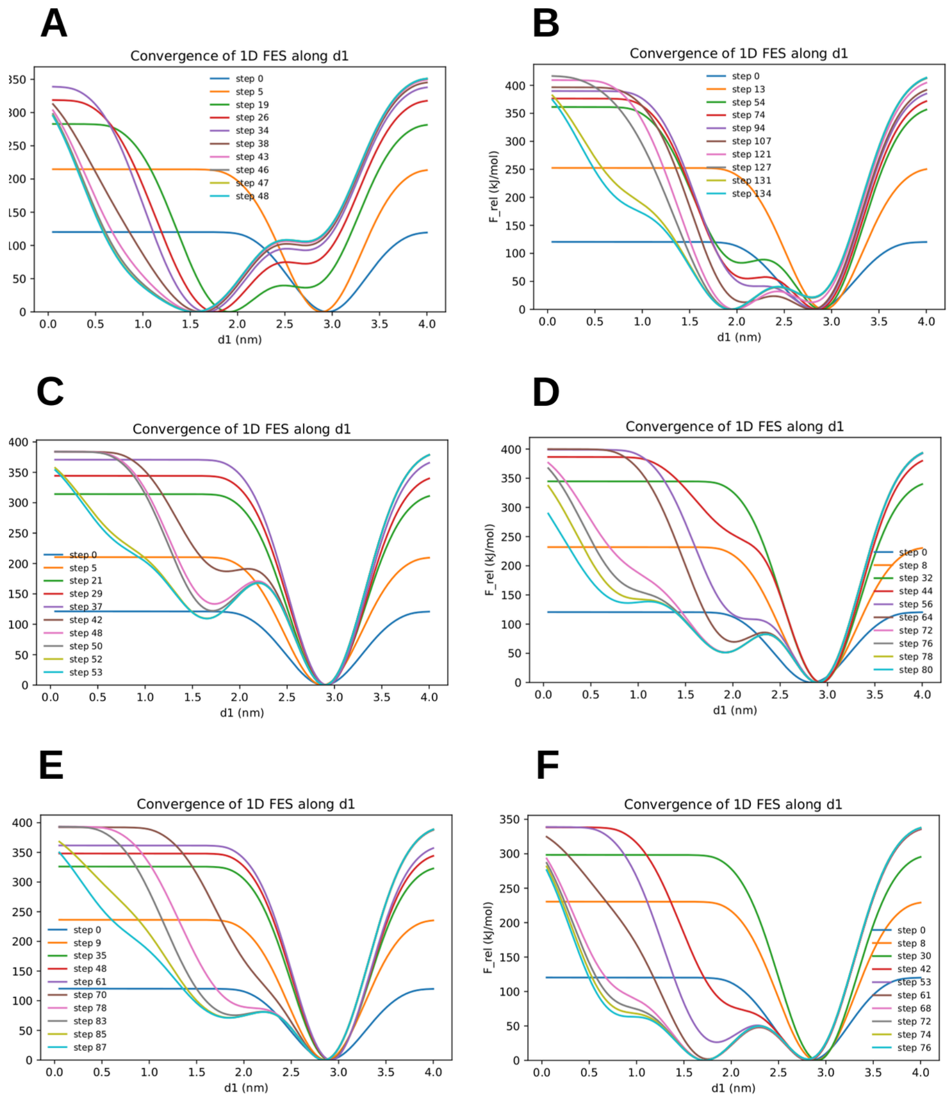


**Figure S24:** Convergence analysis of the one–dimensional free energy surface (FES) along the collective variable $d1$ for $Cu^{+}$ transport in **AtHMA5**. Panels (A–F) correspond to six independent metadynamics replicas. Each colored curve represents the FES reconstructed at different stages of the simulations during bias accumulation.

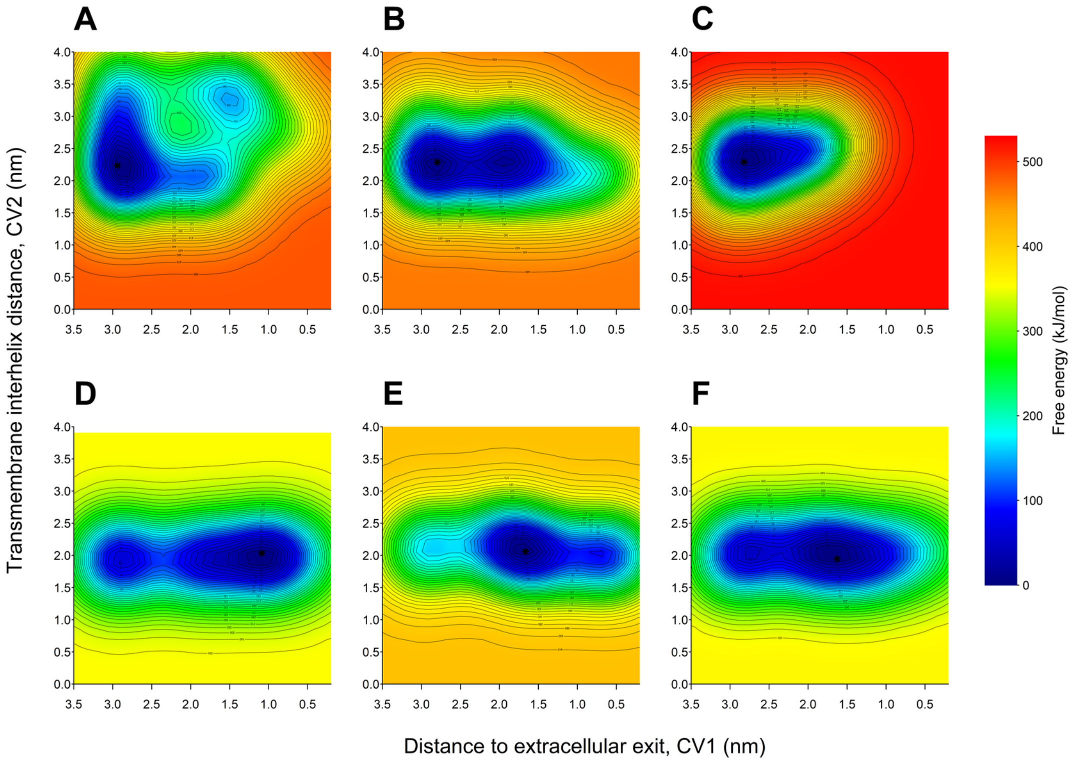


**Figure S25:** Two–dimensional free energy surfaces (FES) describing $Cu^+$ transport in phosphorylated AtHMA5 (AtHMA5-P) obtained from six independent metadynamics replicas (A–F). The collective variables correspond to the distance to the extracellular exit (CV1) and the transmembrane interhelical distance (CV2). Free energies are reported in kJ/mol and referenced to the global minimum of each replica. The replicas consistently display a dominant low-energy basin within the transmembrane region and replica-dependent variations in the extension of the basin along CV1, reflecting differences in sampling of intermediate $Cu^+$ coordination states in the phosphorylated conformation.

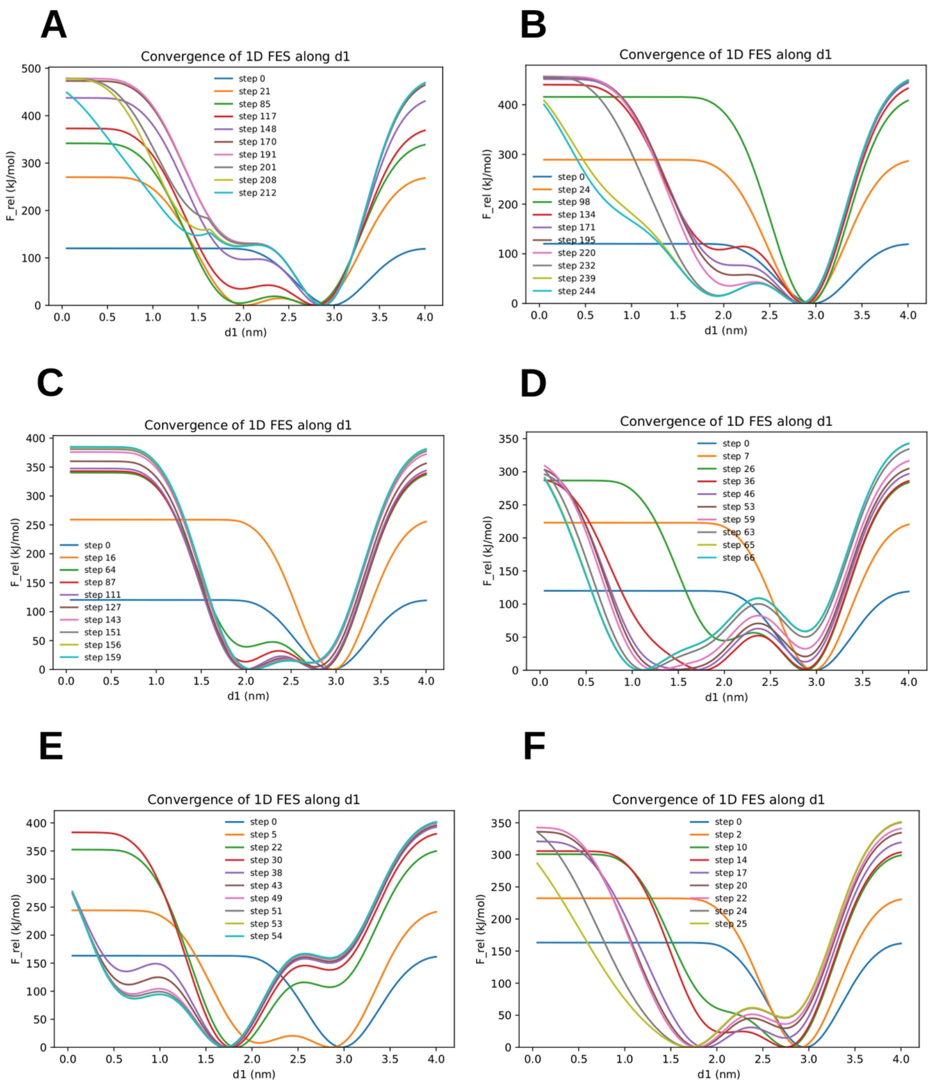


**Figure S26:** Convergence analysis of the one–dimensional free energy surface (FES) along the collective variable $d1$ for $Cu^{+}$ transport in the phosphorylated transporter **AtHMA5-P**. Panels (A–F) correspond to six independent metadynamics replicas. A stable minimum is consistently observed around $d1 \sim 2.7$–$3.0$ nm, corresponding to the preferred ion coordination region along the transport pathway.

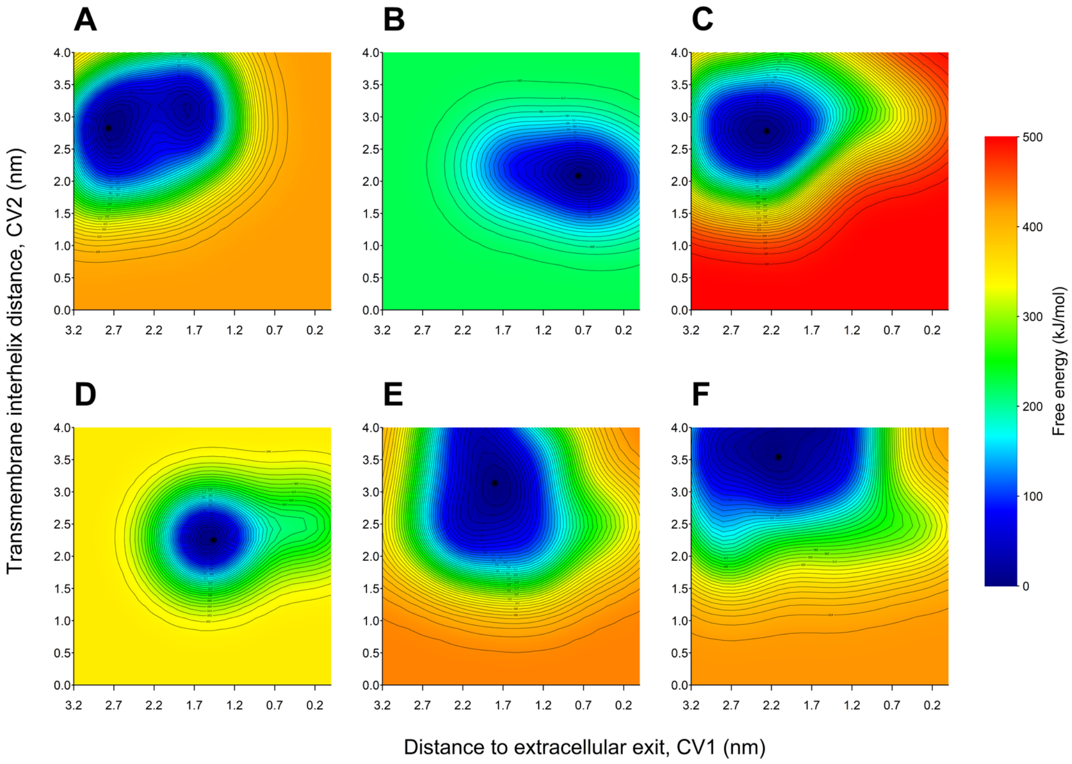


**Figure S27:** Two–dimensional free energy surfaces (FES) describing $Ag^+$ transport in dephosphorylated LpCopA obtained from six independent metadynamics replicas (A–F). The replicas reveal a dominant low-energy basin within the transmembrane region, with variations in basin topology and extension along CV1 reflecting differences in sampling of intermediate $Ag^+$ coordination states in the dephosphorylated conformation.

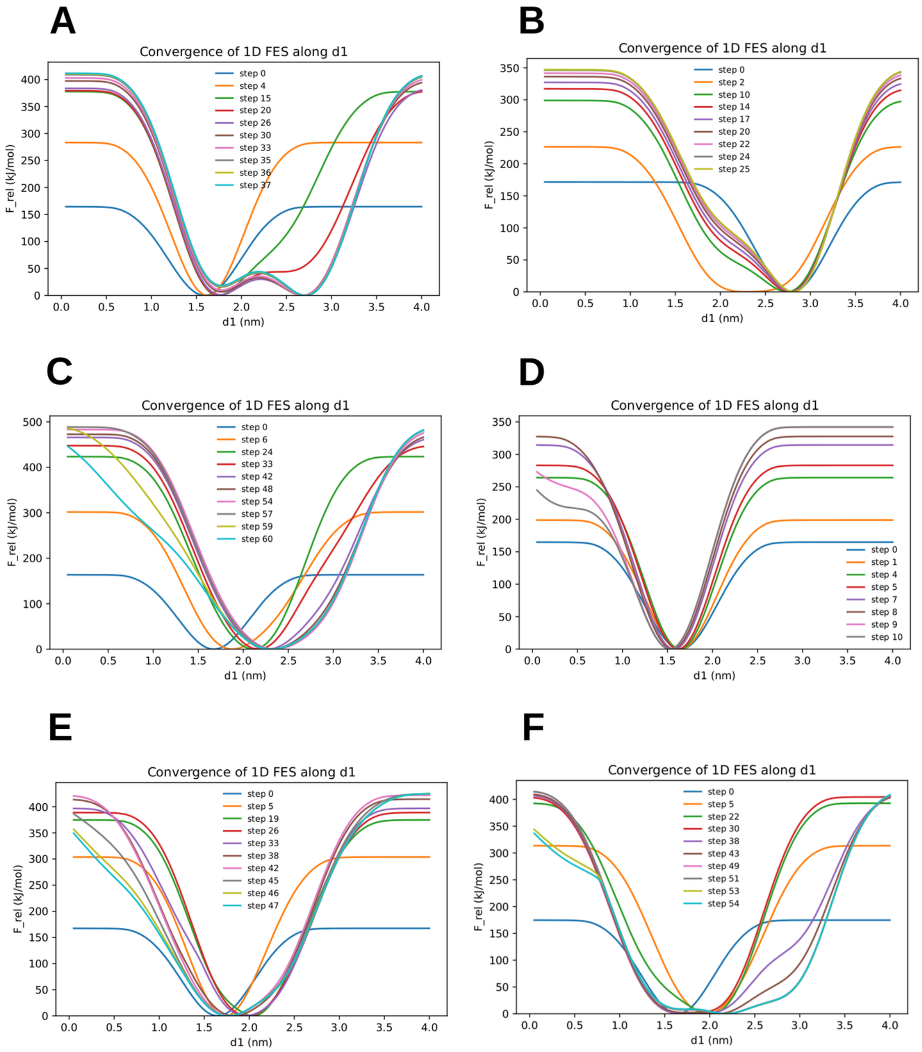


**Figure S28:** Convergence analysis of the one–dimensional free energy surface (FES) along the collective variable $d1$ for $Ag^{+}$ transport in **LpCopA**. Panels (A–F) correspond to six independent metadynamics replicas. Each colored curve represents the FES reconstructed at different stages of the simulation using accumulated bias potentials. As sampling progresses, the profiles gradually stabilize, indicating convergence of the free energy landscape.

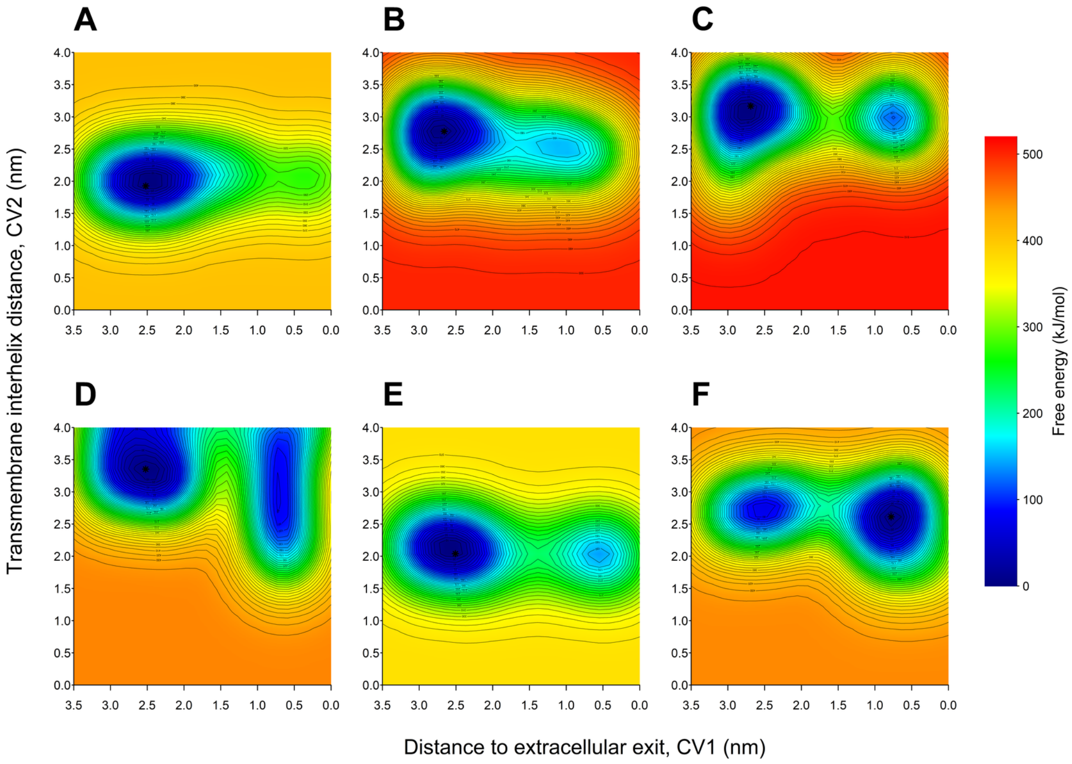


**Figure S29:** Two–dimensional free energy surfaces (FES) describing $Ag^{+}$ transport in phosphorylated LpCopA (LpCopA-P) obtained from six independent metadynamics replicas (A–F). The replicas exhibit a dominant low-energy basin within the pore region and replica-dependent variations in the relative stabilization of intermediate states along CV1, reflecting differences in sampling of $Ag^{+}$ coordination during the phosphorylated transport cycle.

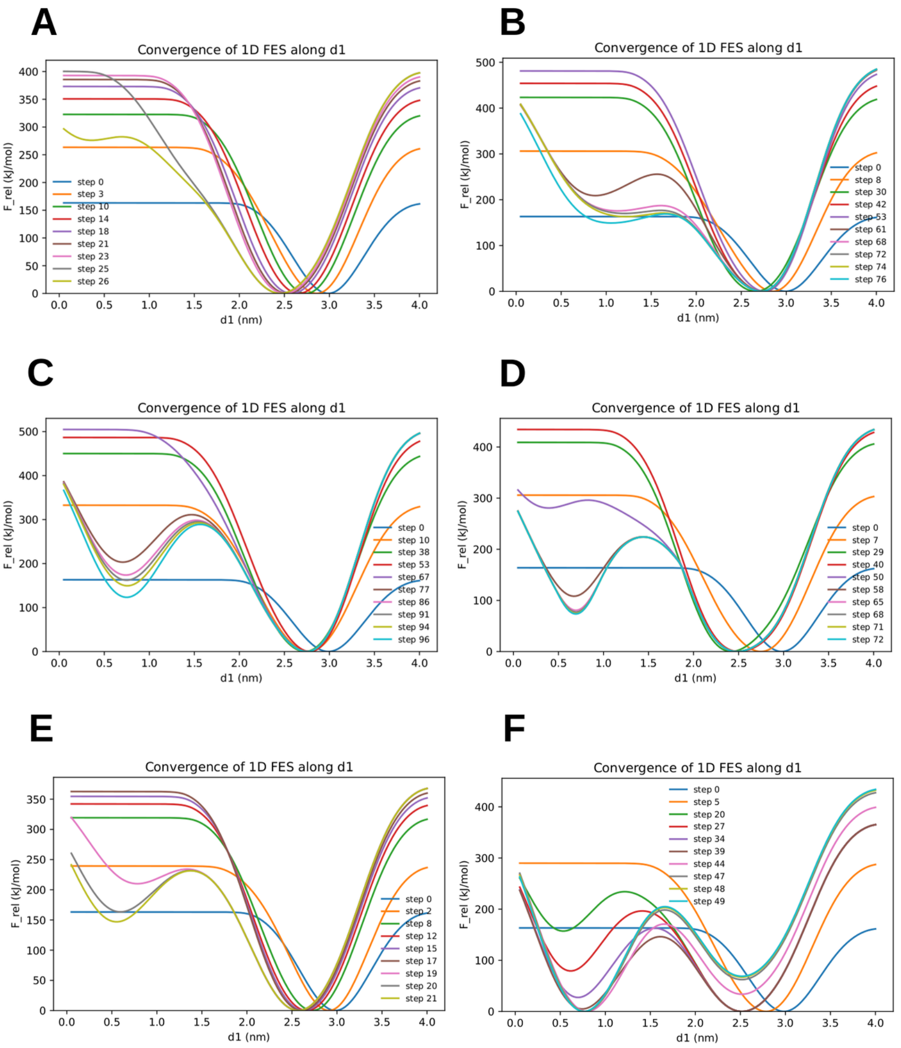


**Figure S30:** Convergence analysis of the one–dimensional free energy surface (FES) along the collective variable $d1$ for $Ag^+$ transport in the phosphorylated transporter **LpCopA-P**. Panels (A–F) correspond to six independent metadynamics replicas. Each colored curve represents the FES reconstructed at different stages of the simulation using accumulated bias potentials. A consistent free-energy minimum emerges along $d1$, corresponding to the preferred ion coordination region along the transport pathway.

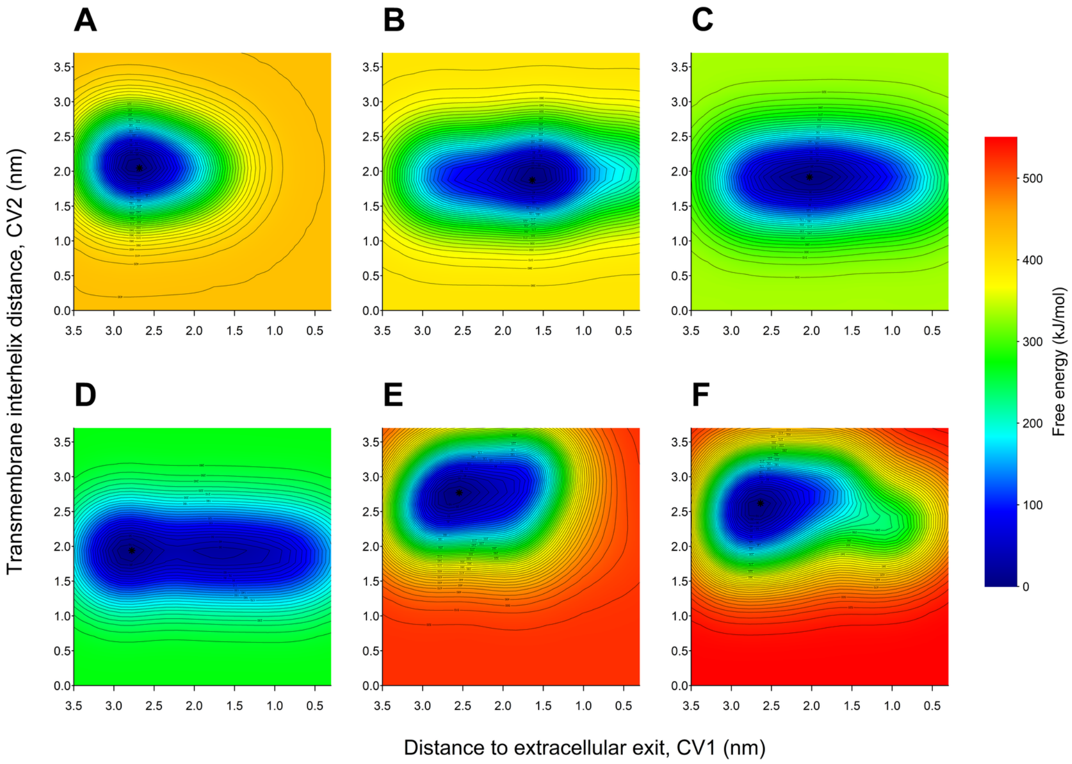


**Figure S31:** Two–dimensional free energy surfaces (FES) describing $Ag^{+}$ transport in dephosphorylated OsHMA9 obtained from six independent metadynamics replicas (A–F).

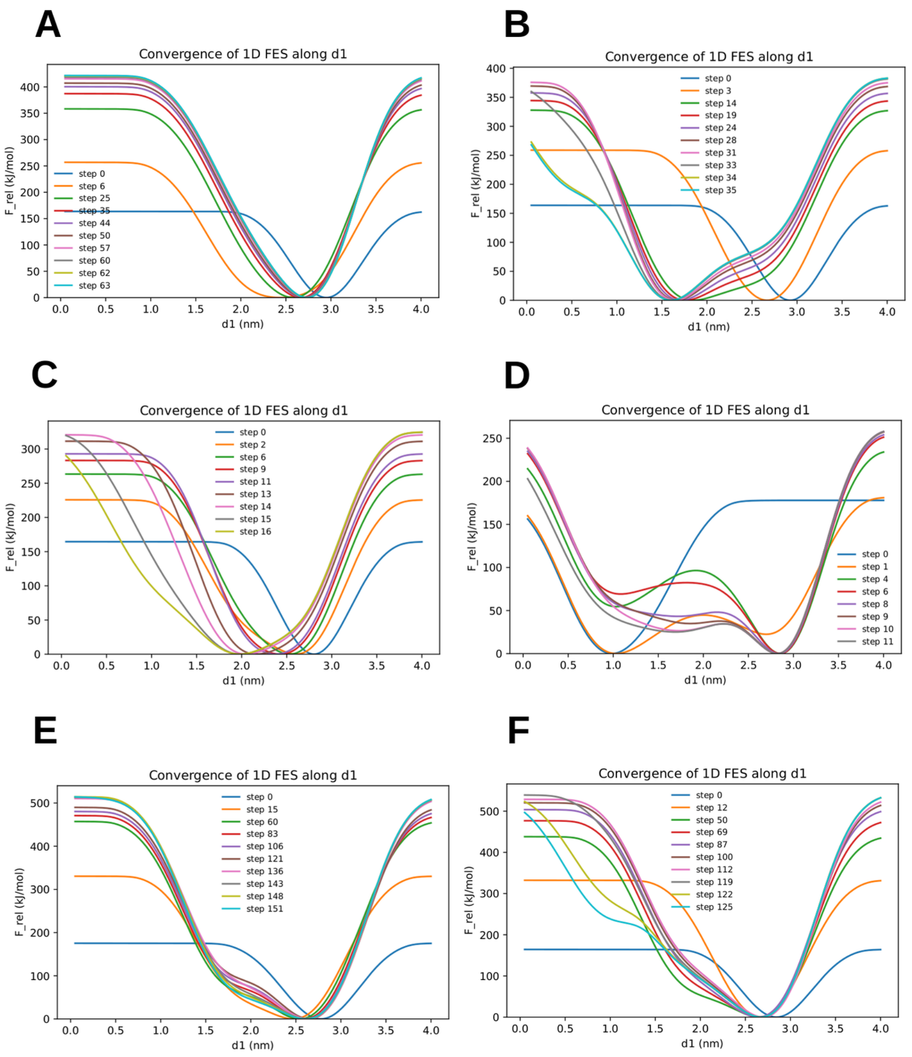


**Figure S32:** Convergence analysis of the one–dimensional free energy surface (FES) along the collective variable $d1$ for $Ag^{+}$ transport in **OsHMA9**. Panels (A–F) correspond to six independent metadynamics replicas. Each colored curve represents the FES reconstructed at different stages of the simulation using accumulated bias potentials.

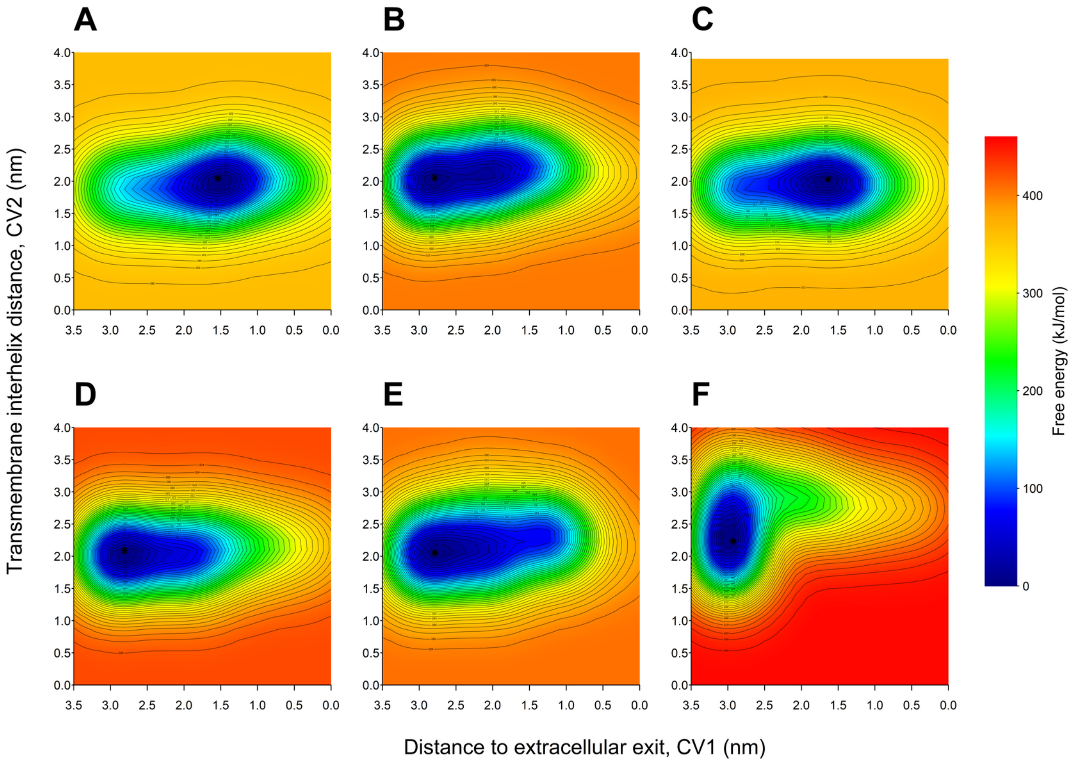


**Figure S33:** Two–dimensional free energy surfaces (FES) describing $Ag^{+}$ transport in phosphorylated OsHMA9 (OsHMA9-P) obtained from six independent metadynamics replicas (A–F). The collective variables correspond to the distance to the extracellular exit (CV1) and the transmembrane interhelical distance (CV2). Free energies are reported in kJ/mol and referenced to the global minimum of each replica.

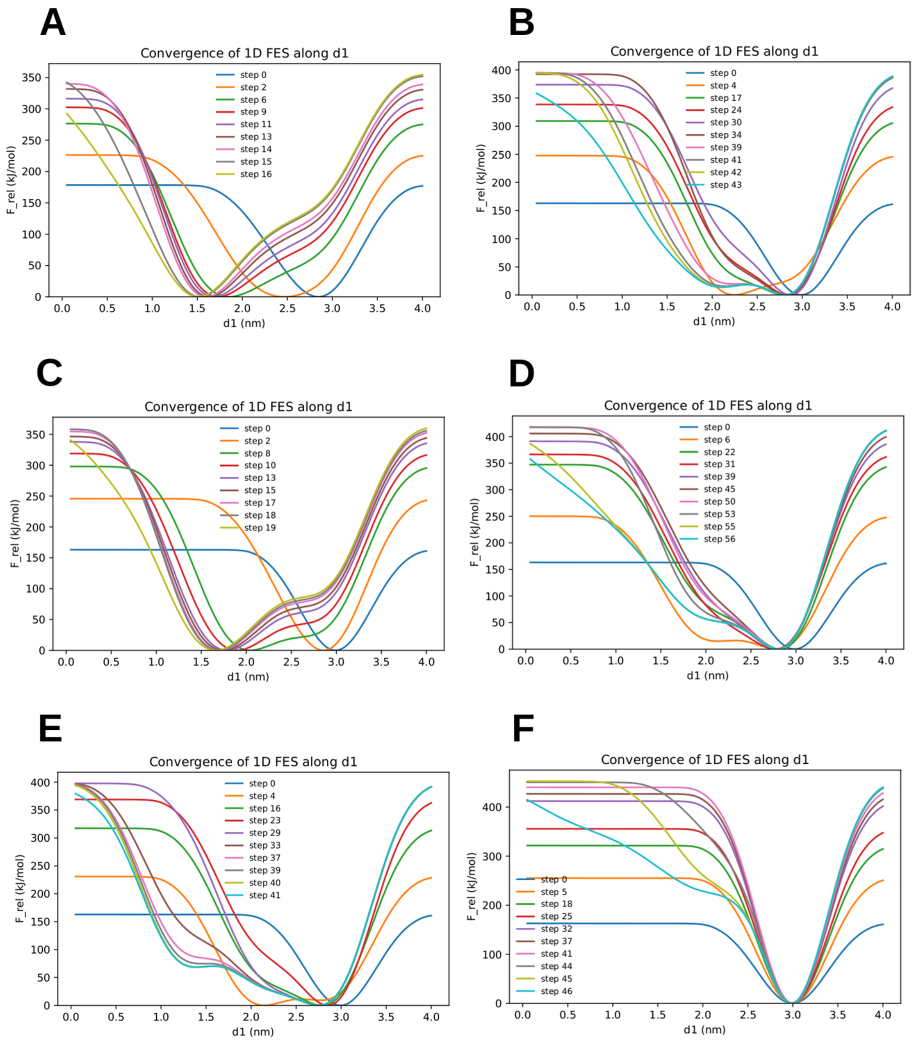


**Figure S34:** Convergence analysis of the one–dimensional free energy surface (FES) along the collective variable $d1$ for $Ag^{+}$ transport in the phosphorylated transporter **OsHMA9-P**. Panels (A–F) correspond to six independent metadynamics replicas. Each colored curve represents the FES reconstructed at successive simulation stages as bias potentials accumulate. A more stable minimum is observed along $d1$, corresponding to a preferred ion coordination region along the transport pathway.

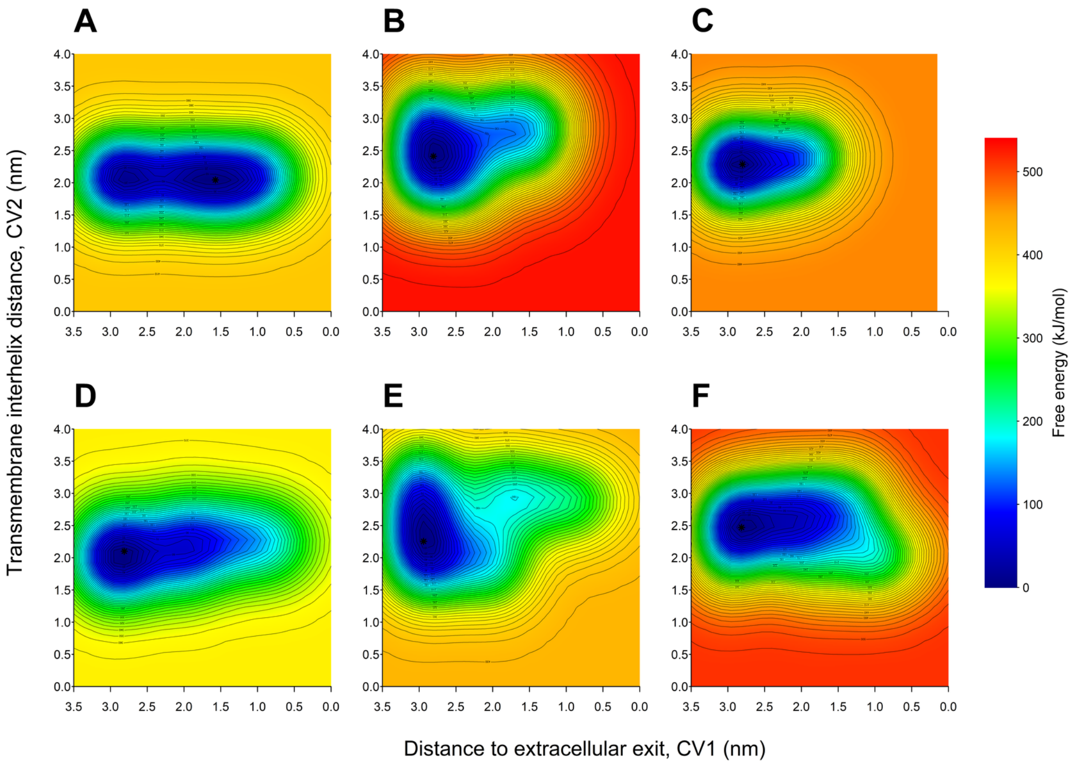


**Figure S35:** Two–dimensional free energy surfaces (FES) describing $Ag^{+}$ transport in dephosphorylated AtHMA5 obtained from six independent metadynamics replicas (A–F). Free energies are reported in kJ/mol and referenced to the global minimum of each replica.

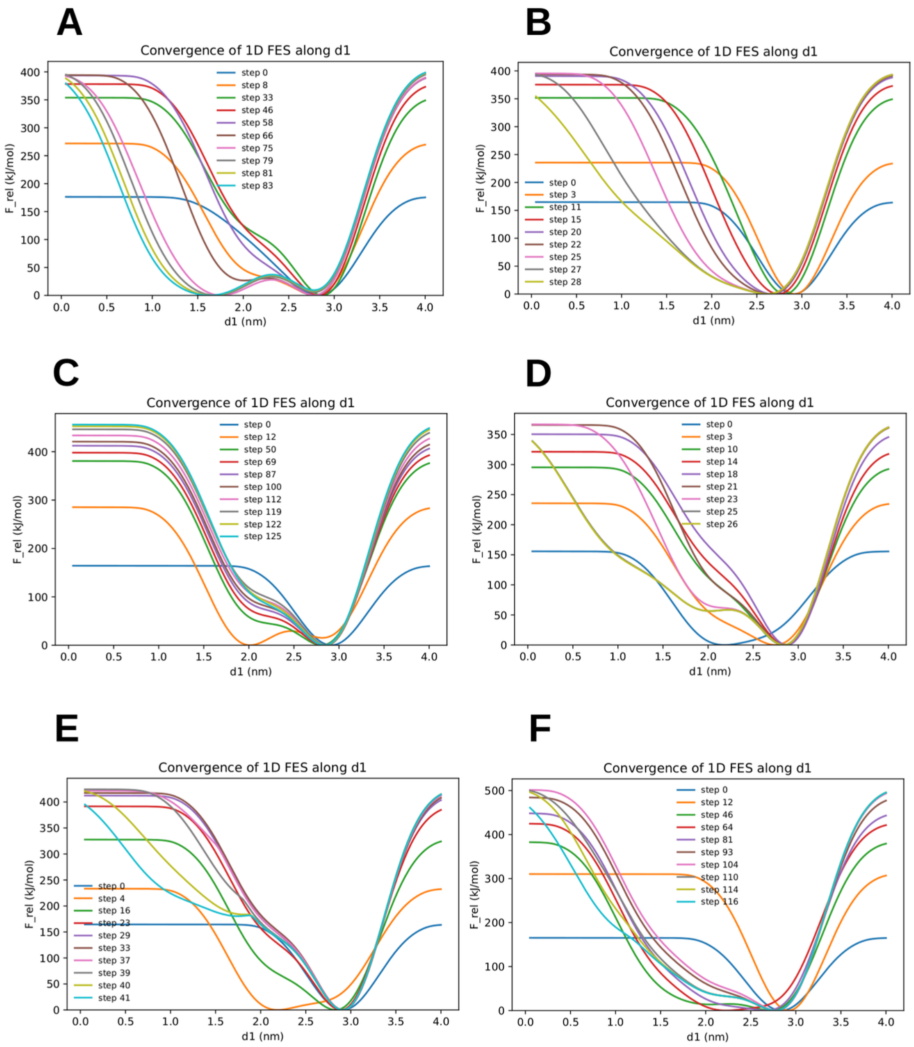


**Figure S36:** Convergence analysis of the one–dimensional free energy surface (FES) along the collective variable $d1$ for $Ag^+$ transport in **AtHMA5**. Panels (A–F) correspond to six independent metadynamics replicas. Each colored curve represents the FES reconstructed at different stages of the simulations during bias accumulation. A consistent free-energy minimum is observed along $d1$, corresponding to an energetically favorable coordination region for the transported ion.

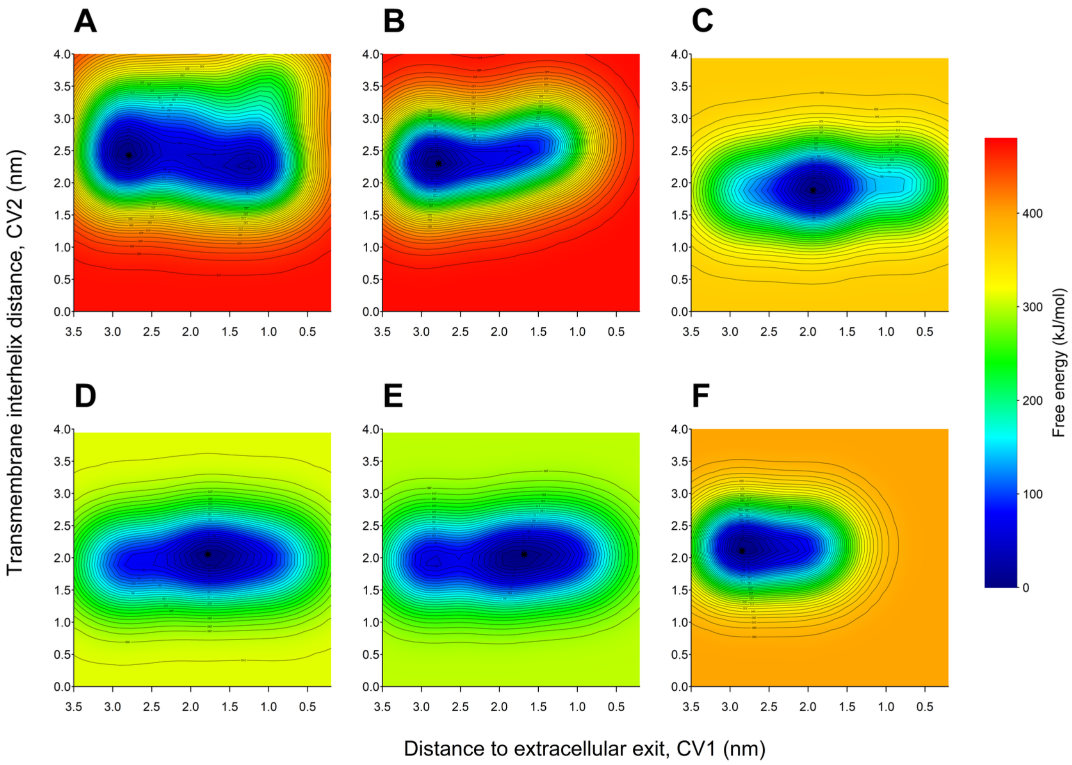


**Figure S37:** Two–dimensional free energy surfaces (FES) describing $Ag^{+}$ transport in phosphorylated AtHMA5 (AtHMA5-P) obtained from six independent metadynamics replicas (A–F). Free energies are reported in kJ/mol and referenced to the global minimum of each replica.

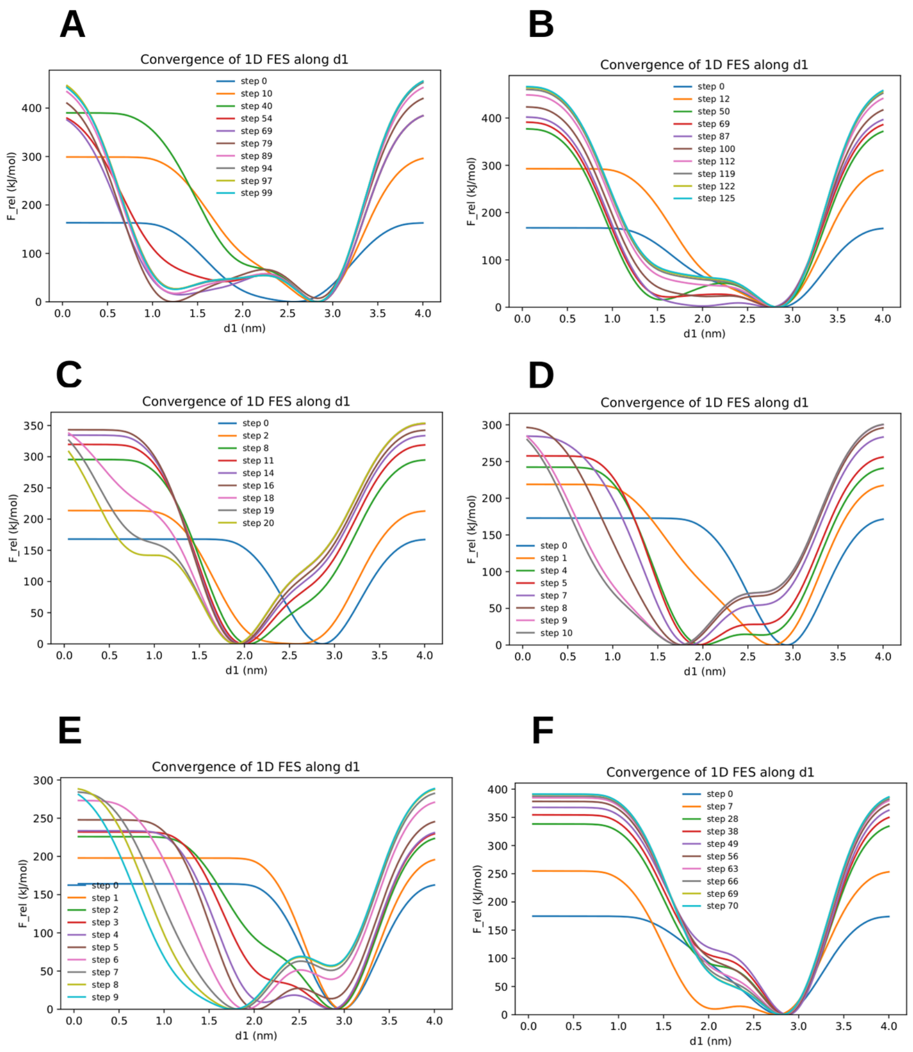


**Figure S38:** Convergence analysis of the one–dimensional free energy surface (FES) along the collective variable $d1$ for $Ag^{+}$ transport in the phosphorylated transporter **AtHMA5-P**. Panels (A–F) correspond to six independent metadynamics replicas. Each colored curve represents the FES reconstructed at different stages of the simulations as bias potentials accumulate.

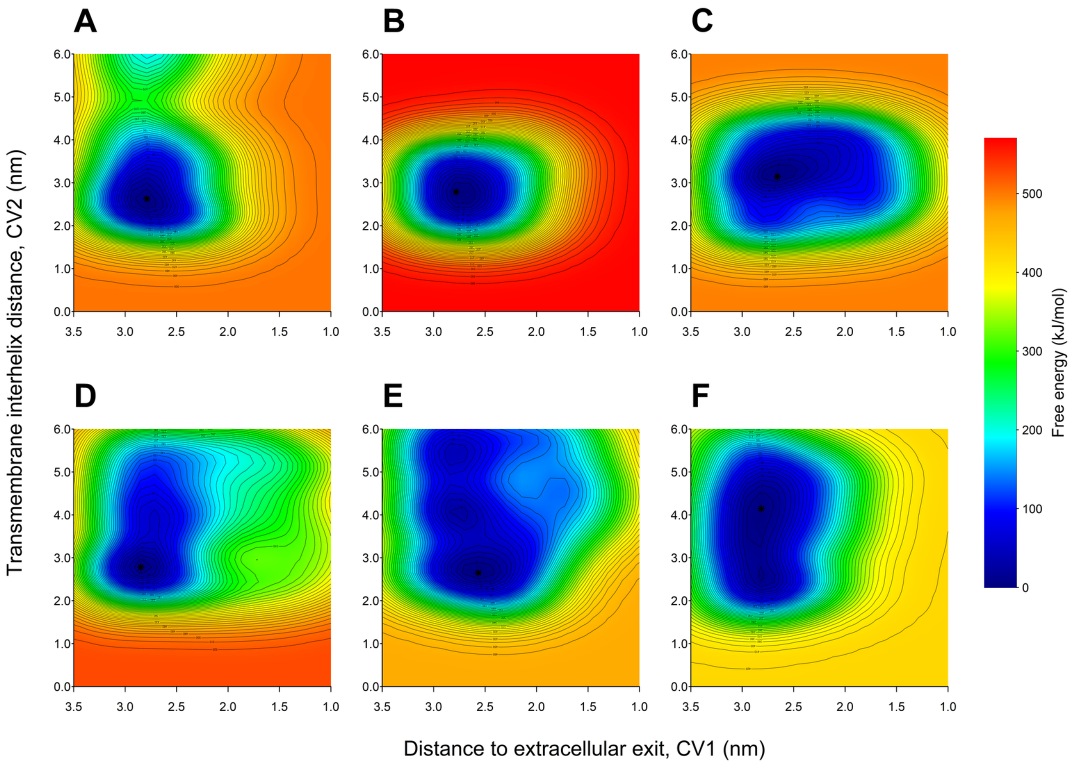


**Figure S39:** Two–dimensional free energy surfaces (FES) describing $Cd^{2+}$ transport in dephosphorylated LpCopA obtained from six independent metadynamics replicas (A–F). Free energies are reported in kJ/mol and referenced to the global minimum of each replica.

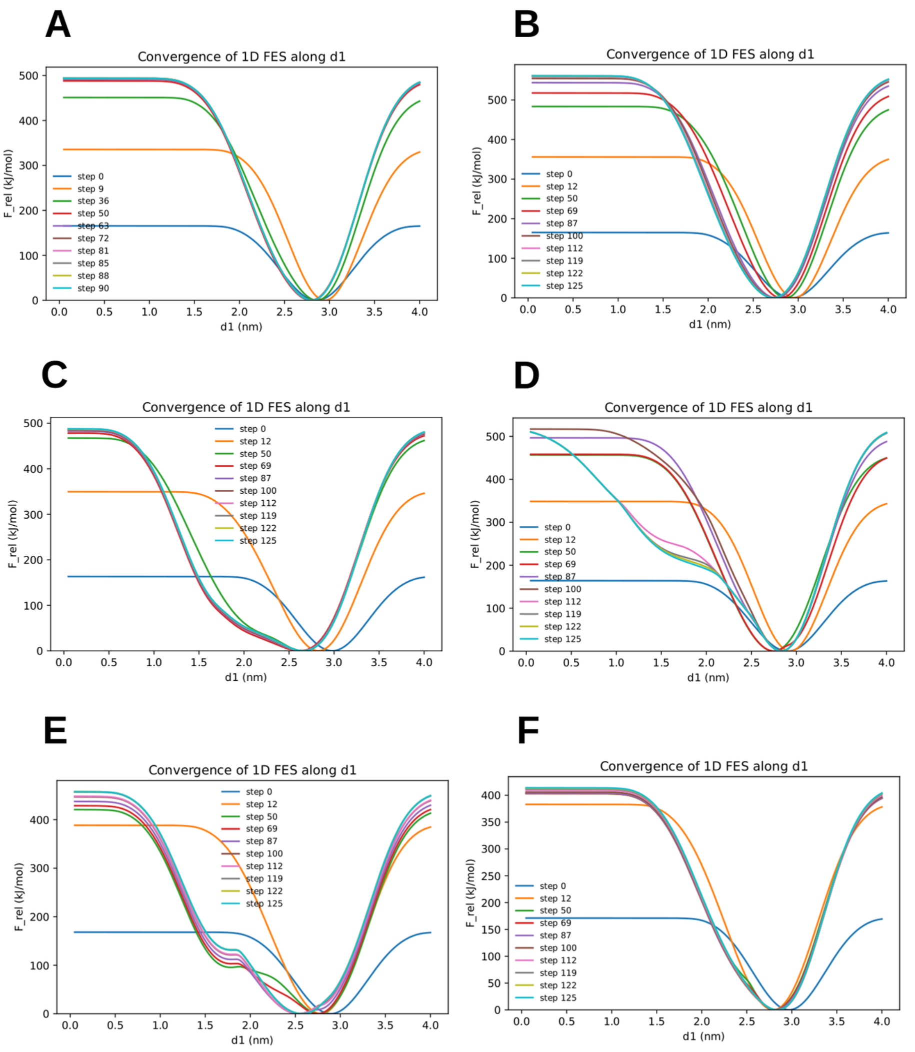


**Figure S40:** Convergence analysis of the one–dimensional free energy surface (FES) along the collective variable $d1$ for $Cd^{2+}$ transport in **LpCopA**. Panels (A–F) correspond to six independent metadynamics replicas. Each colored curve represents the FES reconstructed at different stages of the simulation using accumulated bias potentials. A

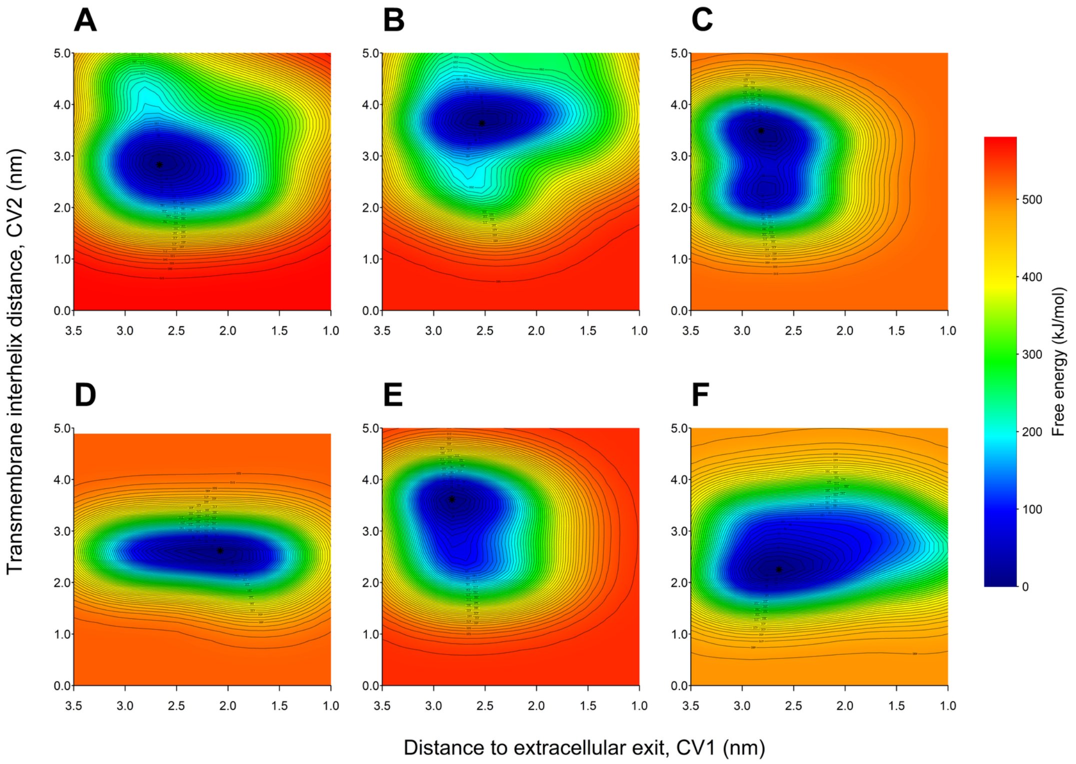


**Figure S41:** Two–dimensional free energy surfaces (FES) describing $Cd^{2+}$ transport in phosphorylated LpCopA (LpCopA-P) obtained from six independent metadynamics replicas (A–F).

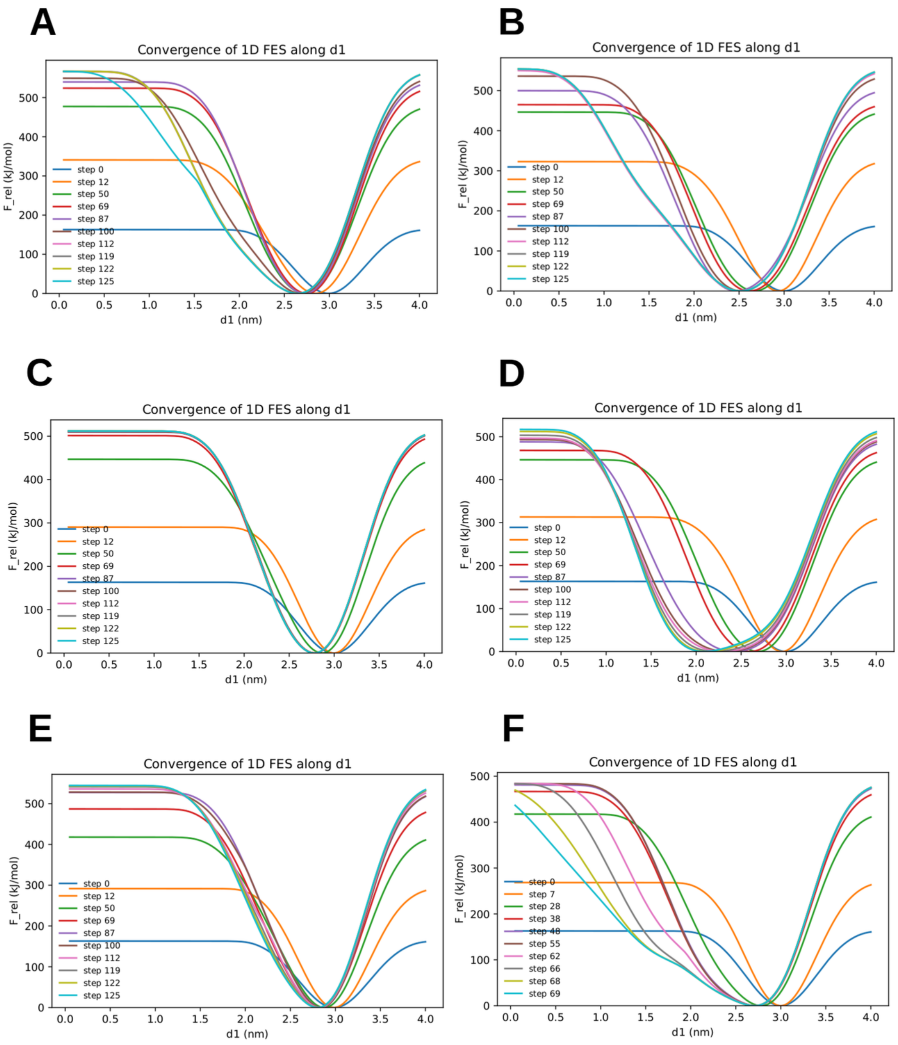


**Figure S42:** Convergence analysis of the one–dimensional free energy surface (FES) along the collective variable $d1$ for $Cd^{2+}$ transport in the phosphorylated transporter **LpCopA-P**. Panels (A–F) correspond to six independent metadynamics replicas. Each colored curve represents the FES reconstructed at different stages of the simulation using accumulated bias potentials.

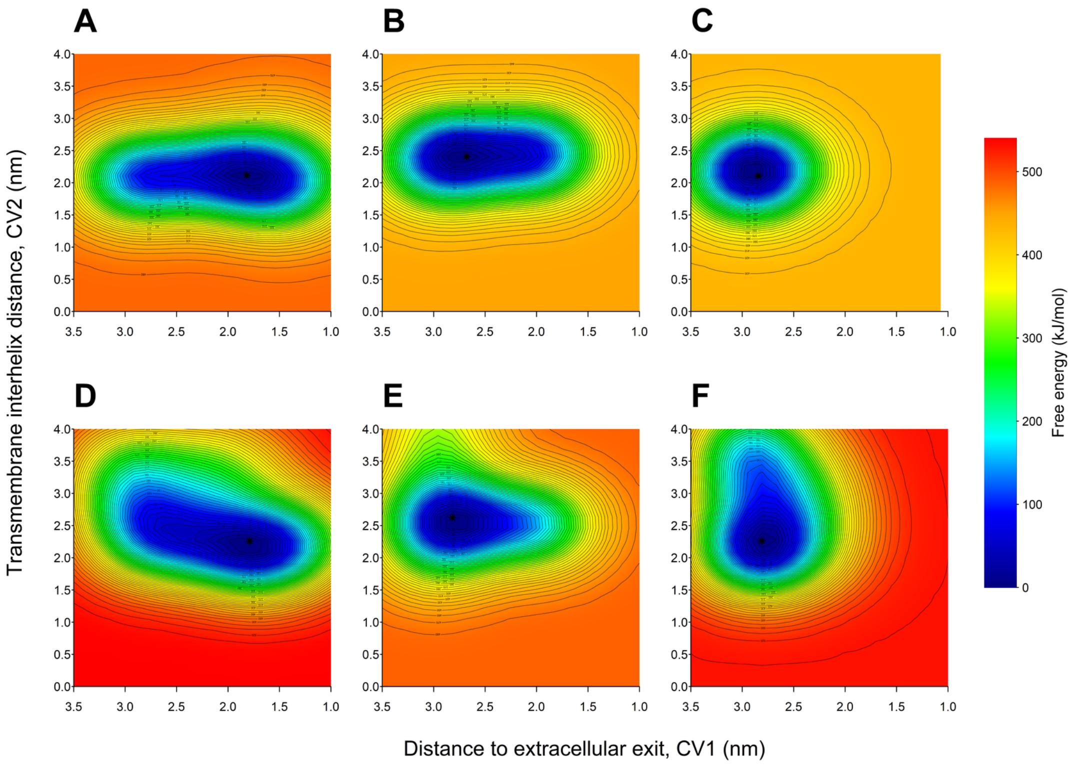


**Figure S43:** Two–dimensional free energy surfaces (FES) describing $Cd^{2+}$ transport in dephosphorylated OsHMA9 obtained from six independent metadynamics replicas (A–F). Free energies are reported in kJ/mol and referenced to the global minimum of each replica.

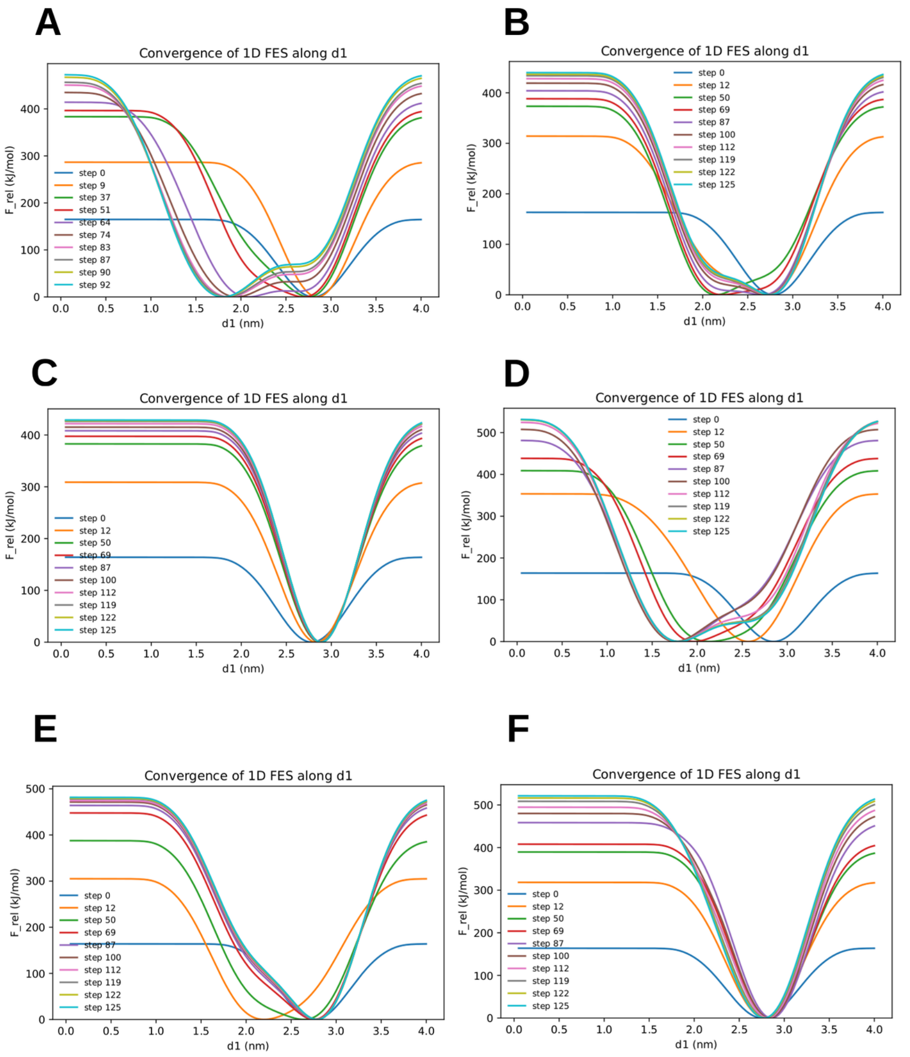


**Figure S44:** Convergence analysis of the one–dimensional free energy surface (FES) along the collective variable $d1$ for $Cd^{2+}$ transport in **OsHMA9**. Panels (A–F) correspond to six independent metadynamics replicas. Each colored curve represents the FES reconstructed at different stages of the simulation using accumulated bias potentials. A dominant minimum is consistently observed along $d1$, corresponding to the energetically favorable coordination region of $Cd^{2+}$ along the transport pathway.

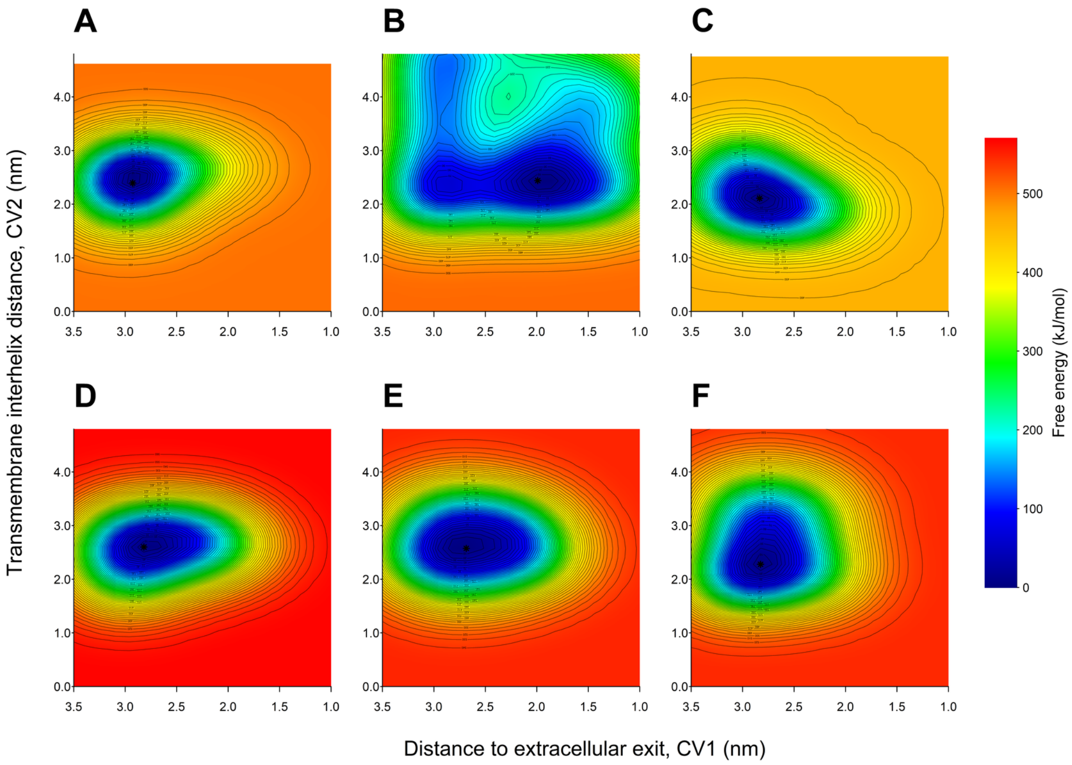


**Figure S45:** Two–dimensional free energy surfaces (FES) describing $Cd^{2+}$ transport in phosphorylated OsHMA9 (OsHMA9-P) obtained from six independent metadynamics replicas (A–F). Free energies are reported in kJ/mol and referenced to the global minimum of each replica. The replicas exhibit dominant low-energy basins within the transmembrane region, with broader distributions along CV2 and increased basin heterogeneity compared to monovalent ions, consistent with distinct coordination behavior of $Cd^{2+}$ in the phosphorylated conformational state.

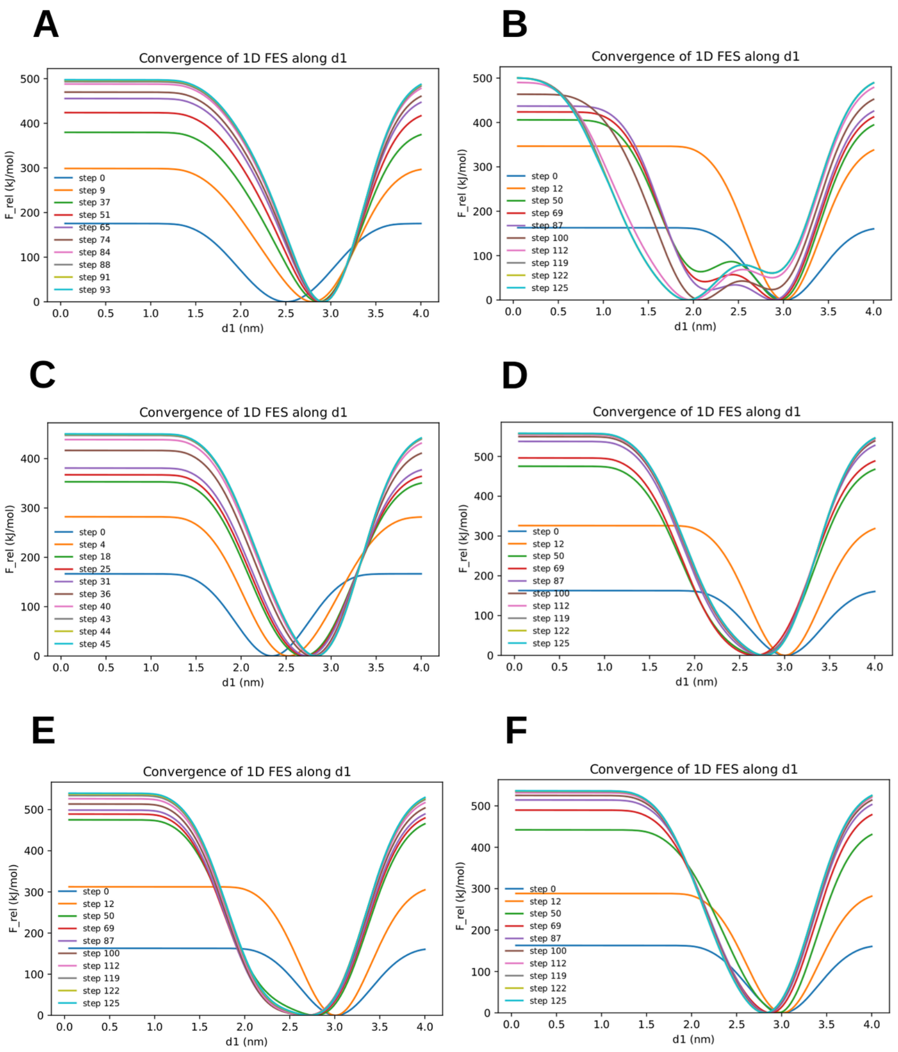


**Figure S46:** Convergence analysis of the one–dimensional free energy surface (FES) along the collective variable $d1$ for $Cd^{2+}$ transport in the phosphorylated transporter **OsHMA9-P**. Panels (A–F) correspond to six independent metadynamics replicas. Each colored curve represents the FES reconstructed at successive simulation stages as bias potentials accumulate.

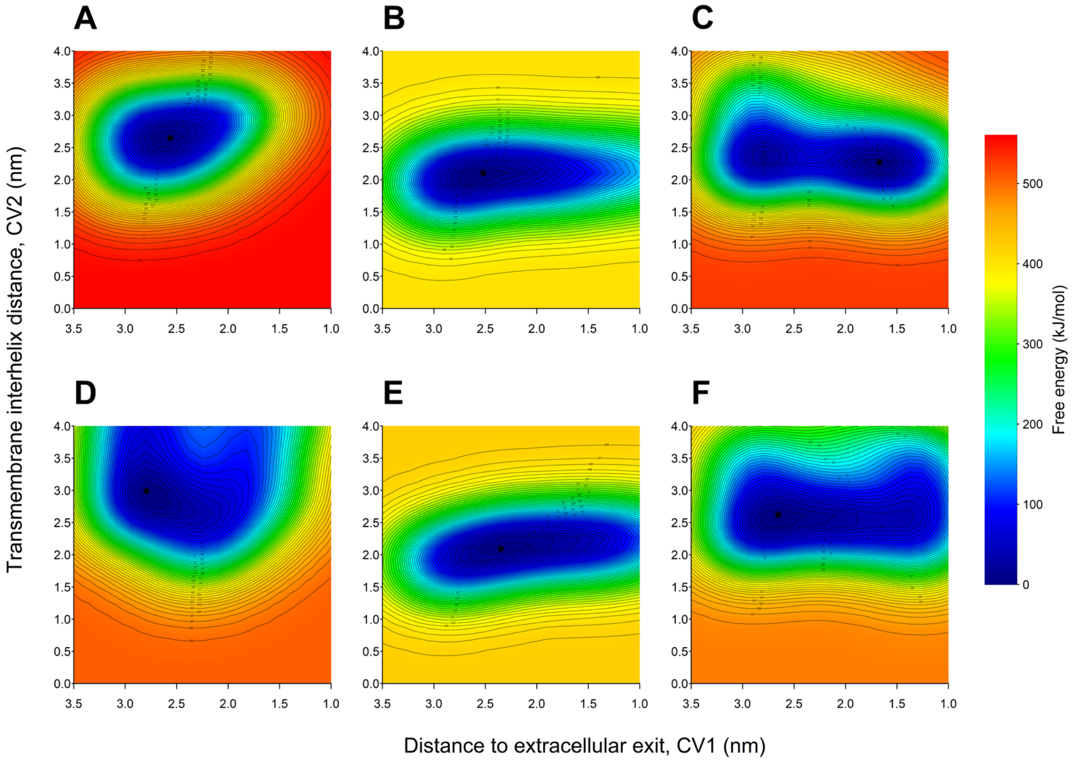


**Figure S47:** Two–dimensional free energy surfaces (FES) describing $Cd^{2+}$ transport in dephosphorylated AtHMA5 obtained from six independent metadynamics replicas (A–F). Free energies are reported in kJ/mol and referenced to the global minimum of each replica.

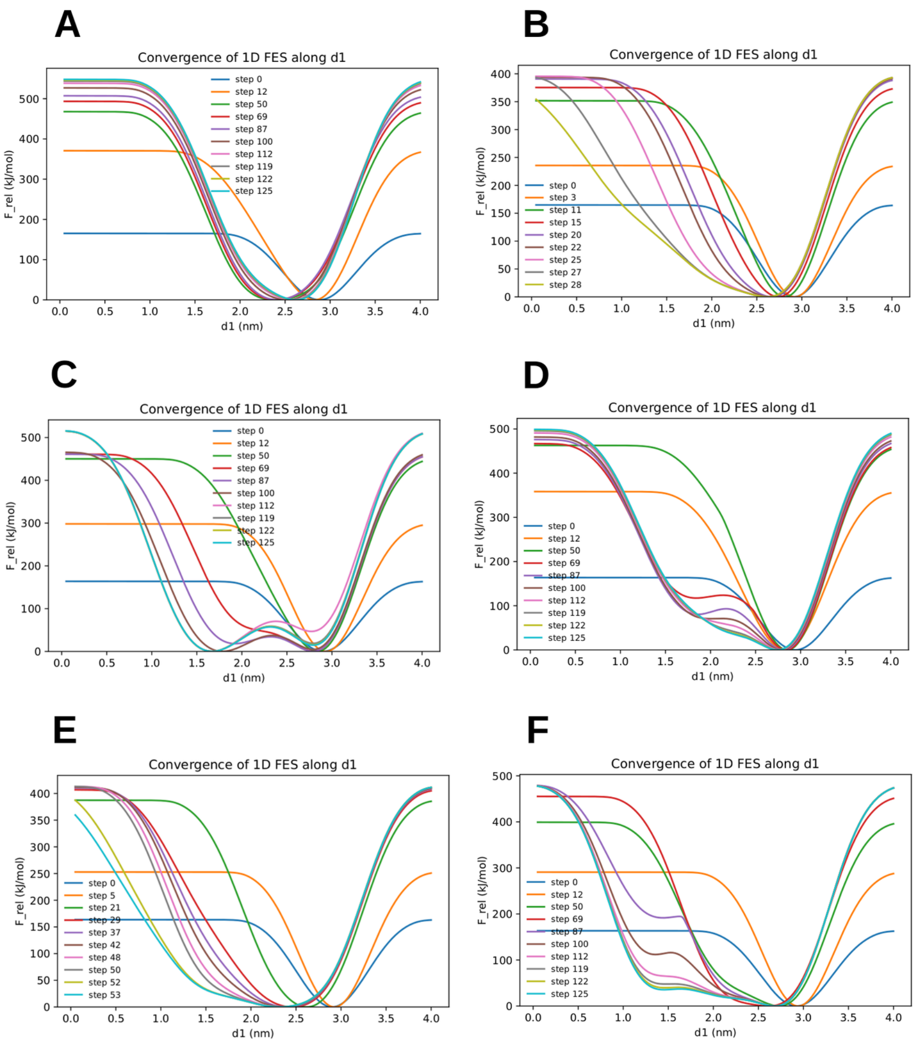


**Figure S48:** Convergence analysis of the one–dimensional free energy surface (FES) along the collective variable $d1$ for $Cd^{2+}$ transport in **AtHMA5**. Panels (A–F) correspond to six independent metadynamics replicas. Each colored curve represents the FES reconstructed at different stages of the simulations during bias accumulation.

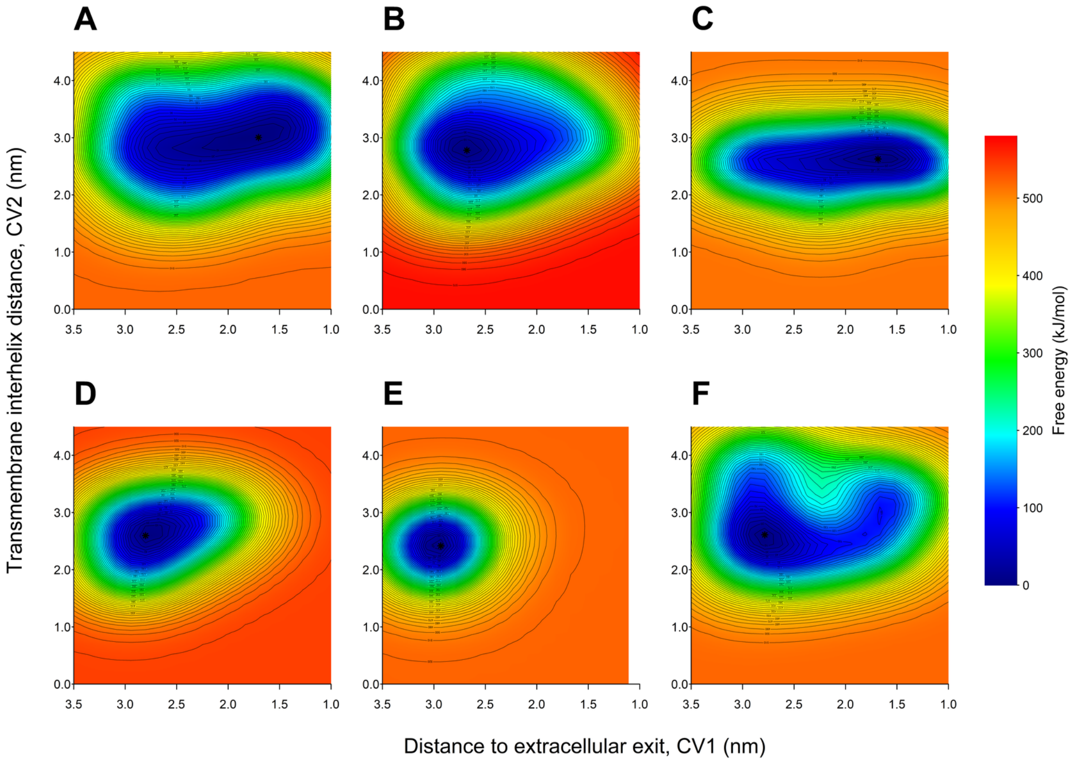


**Figure S49:** Two–dimensional free energy surfaces (FES) describing $Cd^{2+}$ transport in phosphorylated AtHMA5 (AtHMA5-P) obtained from six independent metadynamics replicas (A–F). The collective variables correspond to the distance to the extracellular exit (CV1) and the transmembrane interhelical distance (CV2). Free energies are reported in kJ/mol and referenced to the global minimum of each replica. The replicas reveal dominant low-energy basins within the pore region, with broader sampling along CV2 and increased basin heterogeneity compared to monovalent ions, consistent with distinct coordination and transport behavior of $Cd^{2+}$ in the phosphorylated conformational state.

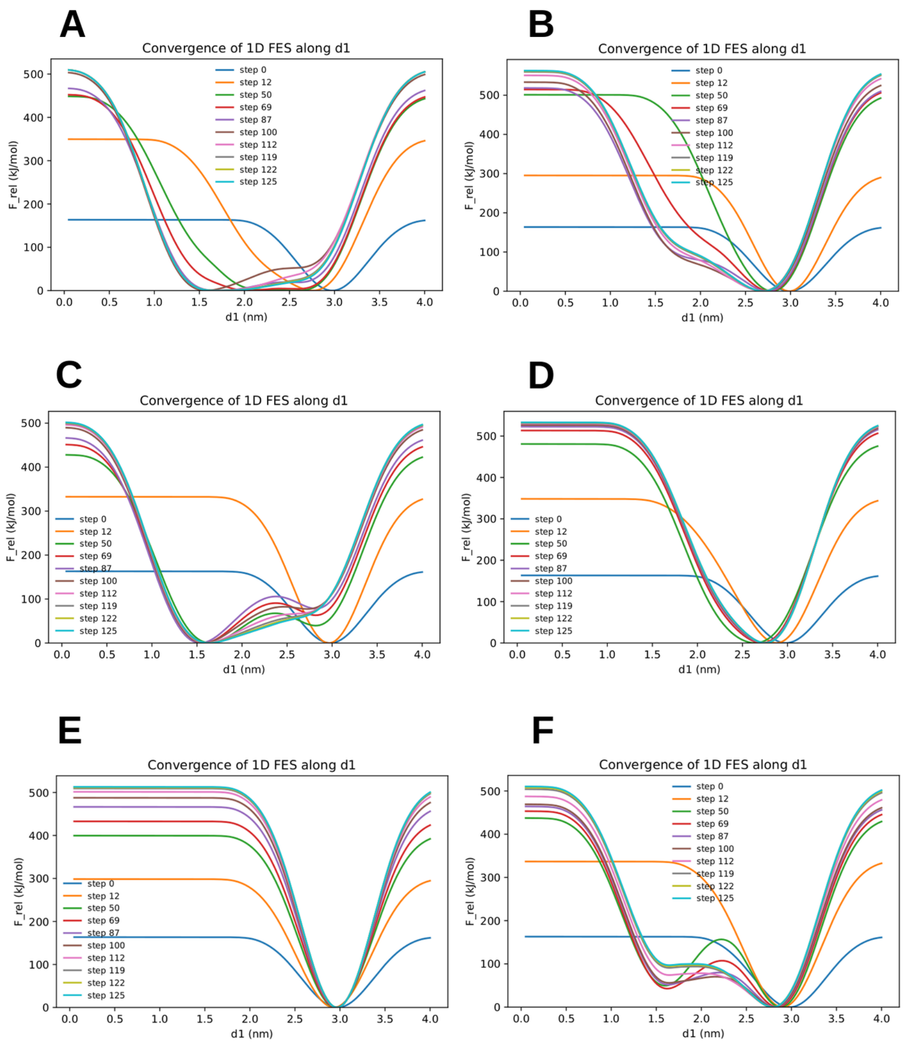


**Figure S50:** Convergence analysis of the one–dimensional free energy surface (FES) along the collective variable $d1$ for $Cd^{2+}$ transport in the phosphorylated transporter **AtHMA5-P**. Panels (A–F) correspond to six independent metadynamics replicas. Each colored curve represents the FES reconstructed at different stages of the simulations as bias potentials accumulate.
